# Nuclear receptor-neurotransmitter coupling links behavior to metabolic state

**DOI:** 10.1101/2025.10.20.683577

**Authors:** Porhathai Malaiwong, Allen F. Schroeder, Tia Brown, Chester J. Wrobel, Madhumanti Dasgupta, Nawaphat Malaiwong, Frank C. Schroeder, Michael P. O’Donnell

## Abstract

Animals must flexibly respond to environmental stimuli to survive, and optimal responses critically depend on the organism’s current needs. Many organisms have evolved both cell-intrinsic and intertissue signaling pathways that integrate metabolic status. However, how this information is encoded in molecular signals is currently not well understood. Here we show that the nematode *C. elegans* employs lipidated neurohormones that combine the neurotransmitter octopamine and fat metabolism-derived building blocks to relay information about lipid metabolic status and drive inhibition of aversive olfactory responses during food removal. Using targeted metabolomics, we show that lipidated neurohormone synthesis requires the carboxylesterase CEST-2.1, which links octopamine-glucosides with endogenous methyl-branched or diet-derived cyclopropane fatty acids that act as agonists of the nuclear receptor and master regulator of fat metabolism, NHR-49/PPARα. Loss of *cest-2.1,* loss of bacterial cyclopropane fatty acid production, or loss of endogenous biosynthesis of the methyl-branched fatty acid substrates of CEST-2.1 mimics the behavioral responses of animals lacking octopamine, indicating that regulation of neurotransmitter-dependent behavior is linked to the coordination of fat metabolism via NHR-49/PPARα. Biosynthesis and subsequent neuromodulation via lipidated neurohormone relies on an intertissue trafficking pathway in which octopamine is shuttled first into the intestine where it is chemically modified, which is likely followed by neuronal import and intracellular hydrolysis to finally release free octopamine. We propose that esterase-dependent synthesis and subsequent hydrolysis of lipidated neurohormones represents a chemical encoding mechanism by which animals integrate information from neurotransmitter signaling and lipid homeostasis to direct appropriate behaviors.

## Main

Responses to sensory stimuli depend on an animal’s physiological state, such that prioritizing, e.g. the pursuit of food over other needs such as rest or reproduction is likely beneficial when nutrients are limiting^1^. Most animals encode highly conserved nutrient sensing pathways that respond to organismal or intracellular changes in levels of sugars, amino acids and fatty acids, which, together with intertissue hormonal signals, transmit information about metabolic state to distant organs^2–4^. Thus, animals must integrate information about past feeding experience to infer the source and value of chemosensory stimuli. In invertebrates, the monoamine neurotransmitter octopamine and its metabolic precursor, tyramine, are key regulators of diverse stress-and satiety-related behavioral and physiological responses. Octopamine and tyramine are structurally and functionally related to the two vertebrate adrenergic neurotransmitters, epinephrine and norepinephrine^5^, which coordinate behaviors, such as increasing locomotion and food seeking while attenuating chemosensory aversion^6–11^, along with metabolic processes, such as increased lipolysis, carbohydrate mobilization and release of pro-catabolic hormones^12–16^, indicating that neurotransmitter signaling integrates metabolic state to regulate behaviors. However, it remains unclear whether there exist specific biochemical mechanisms that link the regulation of central metabolism with monoamine neurotransmitter-dependent behaviors.

### *Cest-2.1* is required for octopamine-dependent behaviors

We hypothesized that chemical modifications to monoamines may be used to integrate information about metabolic state, based on the recent identification of a large family of modular glucosides (MOGLs), which combine a monoamine neurotransmitter moiety (e.g. octopamine, tyramine or serotonin) with building blocks from amino acid and fatty acid metabolism^17–19^ (Fig. 1a). Neurotransmitter-based MOGLs are produced via carboxylesterases orthologous to human hCES1, e.g. encoded by *cest-1.2* and *cest-4*, and production of certain MOGLs is important for survival after starvation in *C. elegans*^18,19^. Given that octopamine plays an important role in satiety, we hypothesized that disruption of octopamine-linked MOGLs may alter feeding-state-dependent behaviors.

**Fig. 1.**
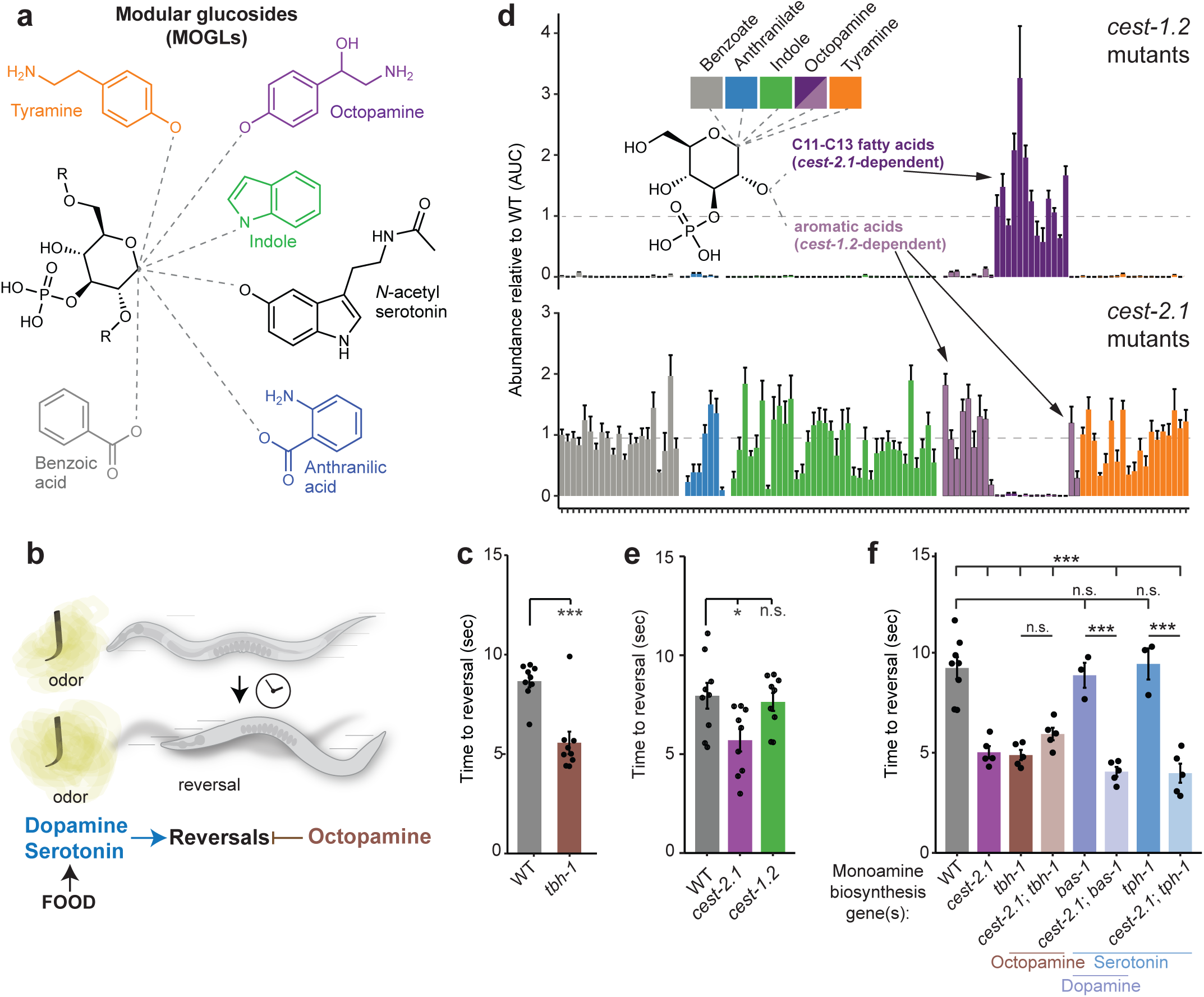
*Cest-2.1* is required for octopamine glucosides and octopamine-dependent behaviors. **a,** Structures of a subset of <u>mo</u>dular <u>gl</u>ucosides identified in *C. elegans* with emphasis on the 1° head group moieties. **b,** Cartoon depicting measurement of aversive olfactory latency in *C. elegans,* referred to as the “smell-on-a-stick” SOS assay. Animals are presented with an odorant in front of the nose and the time until a reversal behavior occurs is recorded. Monoamine pathways that either promote reversals (dopamine, serotonin) or inhibit reversals (octopamine) are depicted. **c, e, f,** SOS reversal latency of animals of the indicated genotypes in response to 33% octanol following 20 minutes of food removal. Each dot indicates an independent assay performed on separate days with at least 20 animals each. Error bars are SEM. Top *** and *, p<0.001 and p<0.05 relative to WT, respectively. Bottom in (**f**) ***, p<0.001 relative to WT in the genetic background of the indicated monoamine biosynthesis mutant. n.s. not significant. ANOVA with TUKEY post hoc correction. **d,** Relative abundance of various putative modular glucosides identified via HPLC-MS in animals of the indicated genotypes. Each bar indicates the abundance (area under the HPLC-MS curve) of an individual metabolite in either *cest-1.2* mutants (top) or *cest-2.1* mutants (bottom), relative to levels found in WT animals. Color indicates the putative molecular identity of the ‘head group’ or the chemical group linked to the anomeric (1°) carbon of glucopyranose. Error bars indicate SEM of 4 independent replicates.

To investigate this hypothesis, we focused on the role of octopamine in the regulation of aversive olfactory responses (locomotory reversals) in animals removed from food^20,21^. Acute avoidance of dilute 1-octanol occurs more rapidly when animals have access to food relative to animals not exposed to food^22^. This avoidance behavior is regulated partly via serotonin^22,23^ and dopamine^24^ signaling and is opposed by octopamine^21,25^ (Fig. 1b). Animals deficient in octopamine production due to the loss of tyramine β-hydroxylase (TBH-1) exhibit faster reversals than wildtype (WT) animals in the absence of food, suggesting that octopamine signaling contributes to the transition from ‘on-food’ to ‘off-food’ response status^21^. We confirmed that animals lacking octopamine signaling exhibited more rapid reversals only after sustained food removal using the well-established ‘smell-on-a-stick’ olfactory behavior assay^26^ (Fig. 1c, Extended Data Fig. 1a).

Next we asked whether animals lacking subsets of octopamine-containing MOGLs exhibited altered acute avoidance responses in the absence of food. To identify mutants defective in the biosynthesis of octopamine-derived MOGLs, we screened a library of 9 available carboxylesterase (*cest*) mutants using HPLC-high resolution mass spectrometry (HPLC-HRMS). This screen revealed that the carboxylesterase gene, *cest-2.1*, was required for the production of a series of MOGLs whose mass spectra indicated that they represent octopamine glucosides (Fig. 1d, Extended Data Fig. 1b). These *cest-2.1*-dependent MOGLs were distinct from the previously reported series of *cest-1.2*-dependent octopamine glucosides^18^ (Fig. 1d). Based on these results we tested *cest-1.2* and *cest-2.1* mutants in the olfactory avoidance assay. We found that *cest-2.1* mutants, but not *cest-1.2* mutants, displayed rapid reversals in response to dilute octanol, suggesting that specific loss of *cest-2.1-*dependent metabolites may lead to these olfactory defects (Fig. 1e).

In principle, the phenotype of *cest-2.1* mutants could result from loss of modulatory antagonists of aversive responses such as octopamine, or alternatively, via increases in the serotonergic or dopaminergic signaling pathway(s). To distinguish between these possibilities, we examined animals lacking both *cest-2.1* and *tph-1* (encoding a tryptophan hydroxylase required for neuronal serotonin production)^27^, *bas-1* (encoding an aromatic amino acid decarboxylase required for serotonin and dopamine production)^28,29^ or *tbh-1*, respectively. If CEST-2.1 normally acts to limit either serotonin or dopamine signaling, we would expect *cest-2.1* response latency defects to be suppressed in these double mutants. We found that rapid response latency of *cest-2.1* mutants was not suppressed in *cest-2.1; tph-1* and *cest-2.1; bas-1* mutants, and that *cest-2.1; tbh-1* mutants exhibited no further decreases in response rate from *cest-2.1* single mutants (Fig. 1f). Together, these results are consistent with CEST-2.1 acting specifically via the octopamine signaling pathway and not via inhibition of serotonin or dopamine signaling. These results suggest that *cest-2.1* is necessary for octopamine-dependent olfactory avoidance responses.

### *Cest-2.1* is required for lipidated octopamine glucosides

Previous studies showed that *cest-1.2* is involved in the biosynthesis of a diverse range of MOGLs, of which only some incorporate octopamine, whereas most are derived from other metabolic building blocks, e.g., amino acid degradation products such as indole^18^. In contrast, the metabolic role of *cest-2.1* has not been studied previously. To fully profile and identify *cest-2.1*-dependent metabolites, we analyzed the metabolomes of *cest-2.1* mutant and WT animals grown in parallel under identical conditions using HPLC-HRMS. Untargeted comparative analysis of the resulting datasets using the Metaboseek platform^30^ revealed a single family of compounds, whose tandem-mass spectrometry (MS^2^) spectra suggested that they represent phosphorylated octopamine glucosides bearing predominantly 11-and 13-carbon fatty acyl moieties (Fig. 1d, Table S1-S2). While the MS^2^ spectra did not allow us to determine the point of attachment of the phosphate and fatty acyl moieties in the *cest-2.1*-dependent octopamine glucosides, we hypothesized that the phosphate is attached to position 3 of the glucose and the fatty acyl moiety to position 2, in analogy to previously identified MOGLs dependent on the closely related *cest-1.2*^18^ (Fig. 2a). With regard to the 11-and 13-carbon fatty acyl moieties in the *cest-2.1*-dependent metabolites, we considered that they could be derived from canonical iso-branched fatty acids, which are abundant in *C. elegans*^31–33^. However, recent studies showed that *C. elegans* additionally accumulates 11-carbon fatty acids derived from bacterial cyclopropyl fatty acids^34,35^ and endogenously produces an unusual C_11_-β-methyl fatty acid, named bemeth#1^35^ (Fig. 2b), which both function as potent agonists of the *C. elegans* HNF4/PPAR-α orthologue, encoded by *nhr-49*^35–38^.

**Fig. 2.**
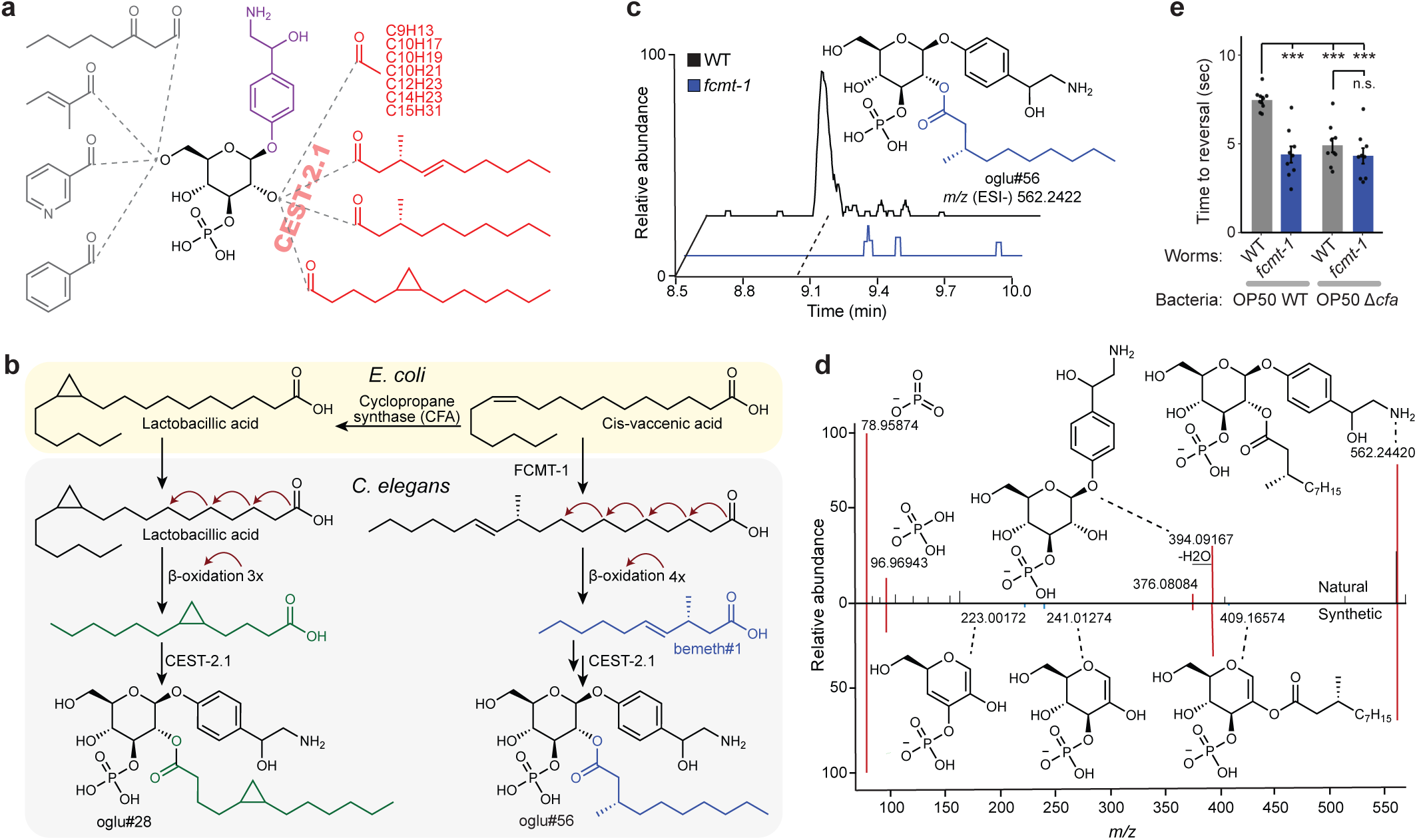
Octopamine glucosides encode a metabolism-dependent neuromodulatory signal. **a**, Putative structural depiction of identified *cest-2.1* dependent octopamine glucosides. **b,** Dietary cis-vaccenic acid is converted to cyclopropyl fatty acids by *E. coli* cyclopropane fatty acid synthase (CFA), or to methyl branched fatty acids by *C. elegans* FCMT-1. These structurally related fatty acids are then β-oxidized and incorporated into octopamine glucosides. **c,** Ion chromatograms of oglu#56 in WT *C. elegans* (black) and in *fcmt-1* mutant *C. elegans* (blue). **d,** Annotated MS^2^ fragmentation spectra of oglu#56 in WT *C. elegans* endo metabolome (top) compared to a synthetic standard of oglu#56 (bottom). **e,** SOS reversal latency of animals of the indicated genotypes fed on either WT or *cfa* mutant *E. coli* in response to 33% octanol following 20 minutes of food removal. Each dot indicates an independent assay performed on separate days with at least 20 animals each. Error bars are SEM. Top: ***, p<0.001 relative to WT *C. elegans* fed on WT *E. coli*, ANOVA with TUKEY post hoc correction. n.s., not significant.

To test whether the fatty acyl moieties in the *cest-2.1*-dependent metabolites are derived from endogenously produced bemeth#1 or bacterial cyclopropyl lipids, we analyzed the metabolomes of *fcmt-1* mutant animals, which are defective in the biosynthesis of C_11_-β-methyl fatty acids^35^, and of WT animals fed *E. coli* defective in the biosynthesis of cyclopropyl fatty acids due to deletion of the *E. coli* cyclopropane fatty acid synthase (*cfa*)^34,39,40^. We found that *fcmt-1* mutant animals specifically lack *cest-2.1*-dependent metabolites featuring unsaturated and saturated C_11_ fatty acyl moieties (Fig. 2c and Extended Data Fig. 2a-c), whereas animals fed *Δcfa* lack *cest-2.1*-dependent metabolites featuring C_13_-fatty acyl moieties with one degree of unsaturation (Extended Data Fig. 2d). Finally, we employed total synthesis (Fig. 2d and Extended Data Fig. 3a-b) to unambiguously establish the structure of oglu#56, the most abundant *cest-2.1*-dependent metabolite that was also *fcmt-1*-dependent, which revealed a 3’-*O*-phosphorylated octopamine glucoside bearing a β-methyldecanoic fatty acyl moiety in position 2 of the glucose, consistent with the known role of FCMT-1 in the biosynthesis of C_11_-β-methyl fatty acids^35^ (see Fig. 2d and Extended Data Fig. 3a-b). Taken together, these results show that *cest-2.1* is specifically required for the biosynthesis of octopamine-based MOGLs that incorporate bacterially-derived fatty acids and endogenous fatty acid agonists of the nuclear receptor NHR-49/PPAR-α. As NHR-49/PPAR-α is a master regulator of fatty acid biosynthesis and breakdown^36–38^, these observations suggest the possibility that lipidated octopamine derivatives may integrate information about the overall metabolic state of the animal.

### Octopamine-glucosides encode a metabolism-dependent neuromodulatory signal

To better understand the role of the two different types of fatty acyl moieties linked to octopamine glucosides (Fig. 2a-d, Table S1), we tested to what extent loss of C_11_-β-methyl fatty acid production via FCMT-1 and/or loss of cyclopropyl fatty acid production by the *E. coli* diet impacts olfactory behavior. To eliminate cyclopropane fatty acids from the standard *C. elegans* diet, we transduced a deletion of the *cfa* gene into *E. coli* OP50 using phage P1*vir*^41^. We then compared octanol avoidance responses of WT and *fcmt-1* mutant worms fed WT or *cfa* knockout *E. coli*, respectively. We found that growth of WT animals on *cfa* knockout bacteria reduced octanol avoidance response latency (Fig. 2e). Similarly, *fcmt-1* mutants, which lack C_11_-β-methyl fatty acids, exhibited phenotypes similar to animals deficient in octopamine or octopamine glucosides (Fig. 2e) when grown on both WT and *cfa* knockout *E. coli* (Fig. 2e). These results indicate that production of NHR-49/PPARα agonists – endogenously produced β-methyl branched fatty acids and dietary cyclopropane fatty acids – is essential for octopamine glucoside-dependent olfactory behavior modulation. These results demonstrate that olfactory behavior regulated by octopamine glucoside signaling is directly tied to the production of NHR-49 lipid agonists that regulate expression of a key enzyme of fat metabolism, FAT-7/SCD-135.

### Octopamine-glucosides are produced in intestinal lysosomal related organelles

We next explored the cellular basis of octopamine glucoside biosynthesis. Prior work indicates that MOGL production requires intestinal lysosome-related organelles (LRO)^17–19^, suggesting that biosynthesis of the *cest-2.1*-dependent octopamine glucosides may also be LRO-dependent. To test this hypothesis, we analyzed the metabolome of mutant animals lacking the RAB32 family GTPase encoded by *glo-1*^42^, in which LRO biogenesis is abolished (Fig. 3a). We found that all *cest-2.1*-dependent octopamine glucosides were also absent in *glo-1* mutants, suggesting that these MOGLs, like *cest-1.2*-and *cest-4*-dependent MOGLs, are produced in LROs (Fig. 3b, Extended Data Fig. 4a). Next we analyzed aversive olfactory responses in *glo-1* mutants. We found that, like *cest-2.1* mutants, *glo-1* mutants exhibit hypersensitive olfactory driven reversals, consistent with the idea that gut LRO-produced octopamine glucosides are necessary normal behavioral responses (Fig. 3c). Further supporting the hypothesis that *cest-2.1*-dependent octopamine glucosides are produced in the intestine, we found that expression of *cest-2.1* in the intestine, but not in the octopamine-producing cells (RIC neurons and gonad sheath cells), restored production of *cest-2.1*-dependent octopamine glucosides in *cest-2.1* mutants (Fig. 3d, Extended Data Fig. 4b-d) and was sufficient to fully rescue behavioral defects of *cest-2.1* mutants (Fig. 3e). Based on these observations, we conclude that fatty-acid linked octopamine glucosides are likely produced in the intestinal LROs to regulate octopamine-dependent olfactory behavior.

**Fig. 3.**
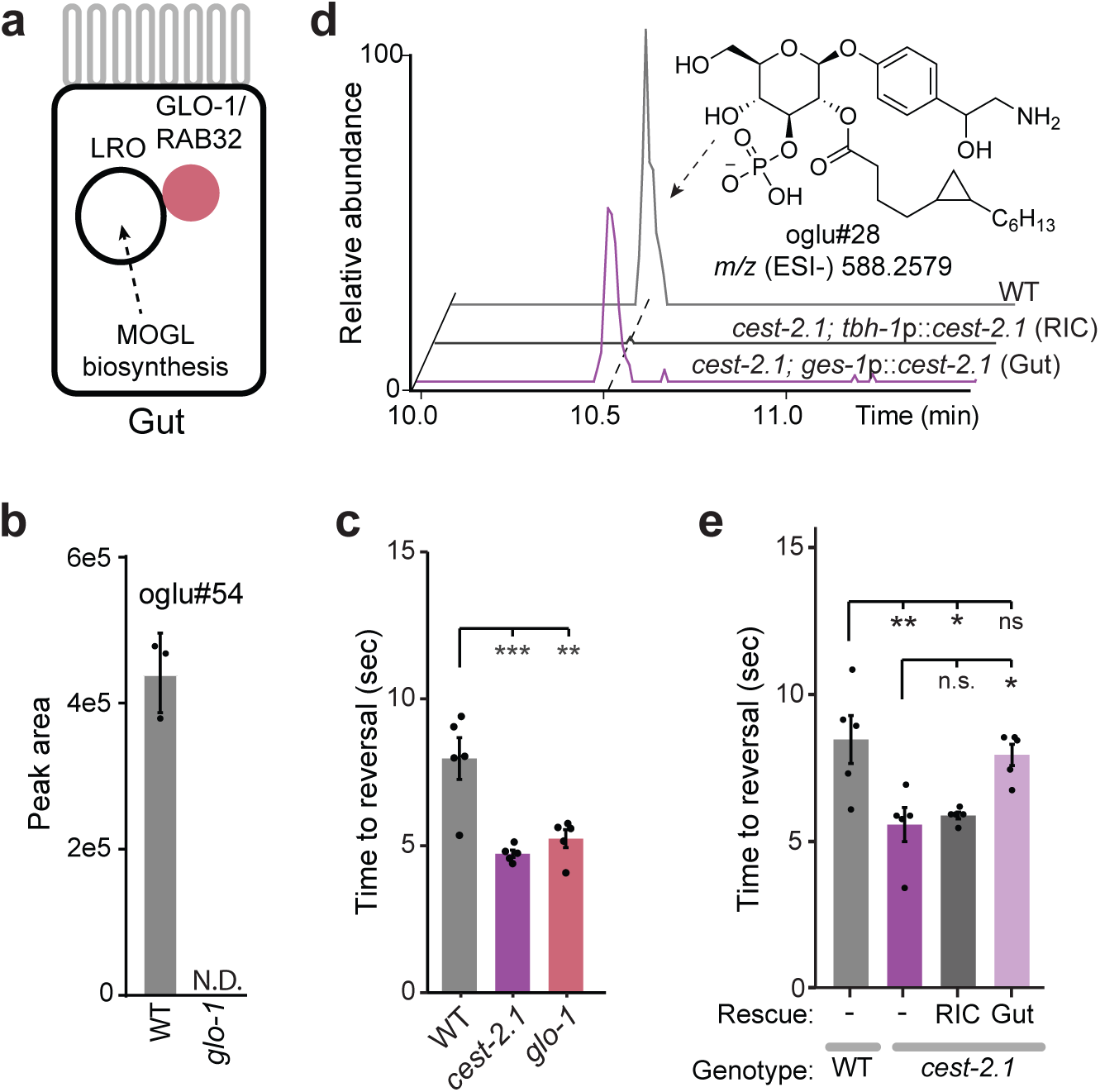
Octopamine-glucosides are produced in intestinal lysosomal related organelles. **a**, Cartoon depicting the role of the RAB-32 GTPase encoded by *glo-1,* in the biogenesis of intestinal lysosomal related organelles (LRO). **b,** Relative abundance of oglu#54 in WT *C. elegans* endo metabolome (gray), and in *glo-1* mutant *C. elegans*. Each dot indicates a biologically independent sample. Error bars are SEM. **c,** SOS reversal latency of animals of the indicated genotype in response to 33% octanol following 20 minutes of food removal. Each dot indicates an independent assay performed on separate days with at least 20 animals each. Error bars are SEM. *** and **, p<0.001 and p<0.01 relative to WT, respectively. ANOVA with TUKEY post hoc correction. **d,** Ion chromatograms of oglu#28 in WT *C. elegans* endo metabolome (gray), in RIC neuron specific reinstatement of *cest-2.1* (*cest-2.1*; *tbh-1*p::cest2.1, black), and in gut specific reinstatement of *cest-2.1* (*cest-2.1*; *ges-1*p::*cest2.1*, purple). **e,** SOS reversal latency of animals of the indicated genotypes bearing transgenic expression of *cest-2.1* cDNA in either the RIC neurons (*tbh-1*p) or the gut (*ges-1*p) or with no rescue transgene (-) in response to 33% octanol following 20 minutes of food removal. Each dot indicates an independent assay performed on separate days with at least 20 animals each. Error bars are SEM. Top ** and *, p<0.01 and p<0.05 relative to WT, respectively. Bottom *, p<0.05 relative to *cest-*mutant. ANOVA with TUKEY post hoc correction. n.s., not significant.

### Glucoside metabolism in the Canal-Associated Neuron is essential for olfactory behavior

The final step of octopamine biosynthesis consists of hydroxylation of its precursor tyramine by TBH-1, which is expressed in only two cell types in *C. elegans*, RIC interneurons and the gonad sheath cells^20,43^. Our observation that *cest-2.1*-dependent octopamine glucosides are produced in the intestine indicates that octopamine released from these cells must transit between tissues, suggesting that octopamine glucosides may serve as an intertissue hormonal signal. To identify additional tissues which may be affected by signaling via octopamine glucosides, we performed RNA-sequencing in *cest-2.1* mutants, with the hypothesis that differentially expressed genes (DEGs) may be enriched among cellular targets of octopamine glucosides (Fig. 4a). An analysis of down-and upregulated genes revealed that nearly half (19 of 44, Table S3) of the DEGs in *cest-2.1* mutants compared to WT worms are expressed in the Canal Associated Neuron^44,45^ (CAN, Fig 4a-b, Table S3). Several of these DEGs are predicted to be involved in glycoside metabolism or transport (Fig. 4b, Table S3), suggesting they may act on octopamine glucosides. We selected a subset of DEGs predicted to regulate glycoside metabolism and/or transport for further characterization. A predicted UDP-galactose/glucuronic acid transporter (encoded by *sqv-7*)^46^, a predicted UDP-glycosyl transferase (encoded by *ugt-64*) and a predicted glucoside hydrolase/glucocerebrosidase (encoded by *gba-4*) were all significantly upregulated in *cest-2.1* mutants relative to WT worms, and are each expressed in CAN (Fig. 4a-b, Table S3). Next, we generated or acquired mutants in these genes and examined their aversive responses to dilute octanol. We found that mutations in each of these genes conferred a quantitatively similar phenotype to *cest-2.1* mutants, suggesting that they may all function in a common process, possibly via CAN (Fig. 4c). In contrast, mutations in a CAN-expressed gene that was downregulated in *cest-2.1* mutants, *ctg-1,* which encodes a SEC14-like lipid binding protein, did not affect olfactory behavior under these conditions (Extended Data Fig. 5). Lastly, we confirmed via cell-specific rescue that *gba-4* functions in the CAN cell to regulate olfactory behavior (Fig. 4d). These results suggest that the CAN cell, possibly via metabolism of octopamine glucosides, regulates olfactory behavior.

**Fig. 4.**
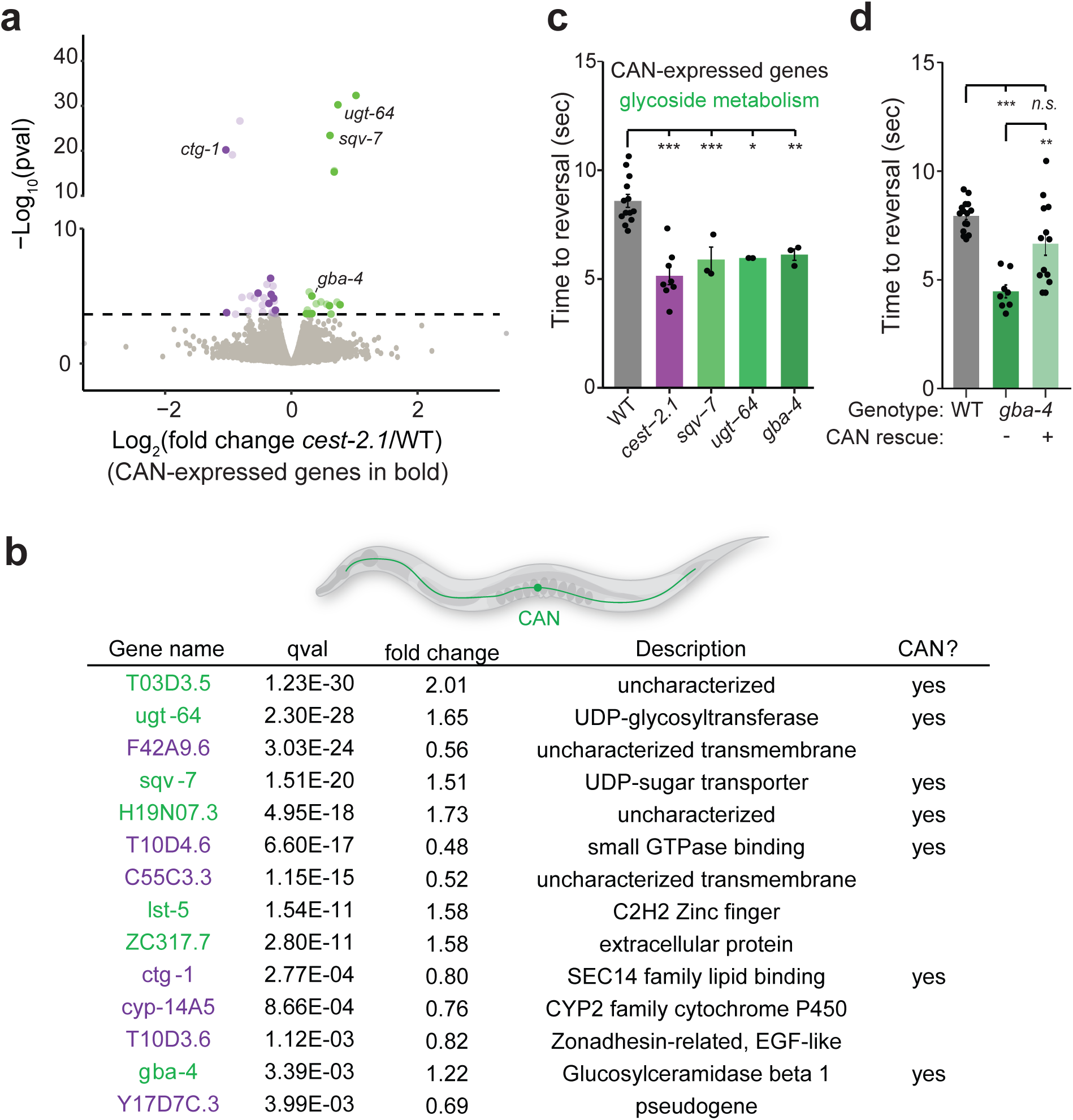
Glucoside metabolism in the CAN cell is essential for olfactory behavior. **a**, Volcano plot showing relative mRNA expression of genes in *cest-2.1* mutants relative to WT worms. Horizontal dashed line indicates the genome-wide significance cutoff (genome-wide q-value < 0.05). Significantly upregulated genes are indicated in green, significantly downregulated genes are indicated in purple. Genes that are known to be expressed in the CAN cell are indicated in bold. Genes of emphasis are indicated with text. **b,** List of top 14 up-or down-regulated genes in *cest-2.1* mutants indicating enrichment for genes expressed CAN and genes putatively involved in glycoside metabolism. **c, d,** SOS reversal latency of animals of the indicated genotype in response to 33% octanol following 20 minutes of food removal. Each dot indicates an independent assay performed on separate days with at least 20 animals each. Error bars are SEM. For (**d**), genotypes bearing transgenic expression of *gba-4* cDNA in the CAN neurons (*ceh-23*p, ‘+’) or with no rescue transgene (-) are indicated. Top, ***, ** and *, p<0.001, p<0.01 and p<0.05 relative to WT, respectively. For (**d**) bottom **, p<0.01 relative to mutant with no rescue transgene. ANOVA with TUKEY post hoc correction.

### Octopamine is transferred to the Canal Associate Neuron via glucoside trafficking

As *gba-4* encodes a putative glucoside hydrolase, this suggests the possibility that *cest-2.1*-dependent octopamine glucosides, which are produced in the intestine, may be imported into and hydrolyzed in the CAN cell, releasing free octopamine. In *C. elegans,* both serotonin and dopamine are produced enzymatically within a subset of neurons, but also can be directly imported into, and function within, non-serotonergic or non-dopaminergic neurons, respectively^47,48^. Our finding that octopamine from *tbh-1*-expressing cells is transferred to other tissues suggests that octopamine, like serotonin and dopamine, may be imported and/or released from non-octopaminergic neurons. A shared feature of neurons with the capacity to release non-natively produced monoamines is the expression of a plasma membrane uptake transporter and a vesicular monoamine transporter (VMAT), the latter of which is encoded by *cat-1* in *C. elegans*^27,43,47–49^. While CAN does not form chemical synapses, it intriguingly expresses *cat-1*, suggesting that under certain circumstances this cell may be capable of releasing monoamines^43,50^ (Fig. 5a). As an initial test of this hypothesis, we reasoned that if true, CAT-1 would be expected to function in both the octopaminergic RIC neurons (to initially synthesize and release free octopamine) and subsequently in the CAN cell (to release octopamine derived from octopamine glucoside hydrolysis, see Fig. 5a). We performed cell-specific rescue experiments in *cat-1* mutants by expressing *cat-1* cDNA in RIC and the gonad sheath (via *tbh-1*p)^20^, CAN (via *ceh-23*p)^51^ or in both cell types, respectively. Consistent with an essential role of *cat-1* in octopamine signaling, we found that *cat-1* mutant animals exhibited octanol responses similar to that of *tbh-1* mutant animals (Fig. 5b, Extended Data Fig. 1a).

**Fig. 5.**
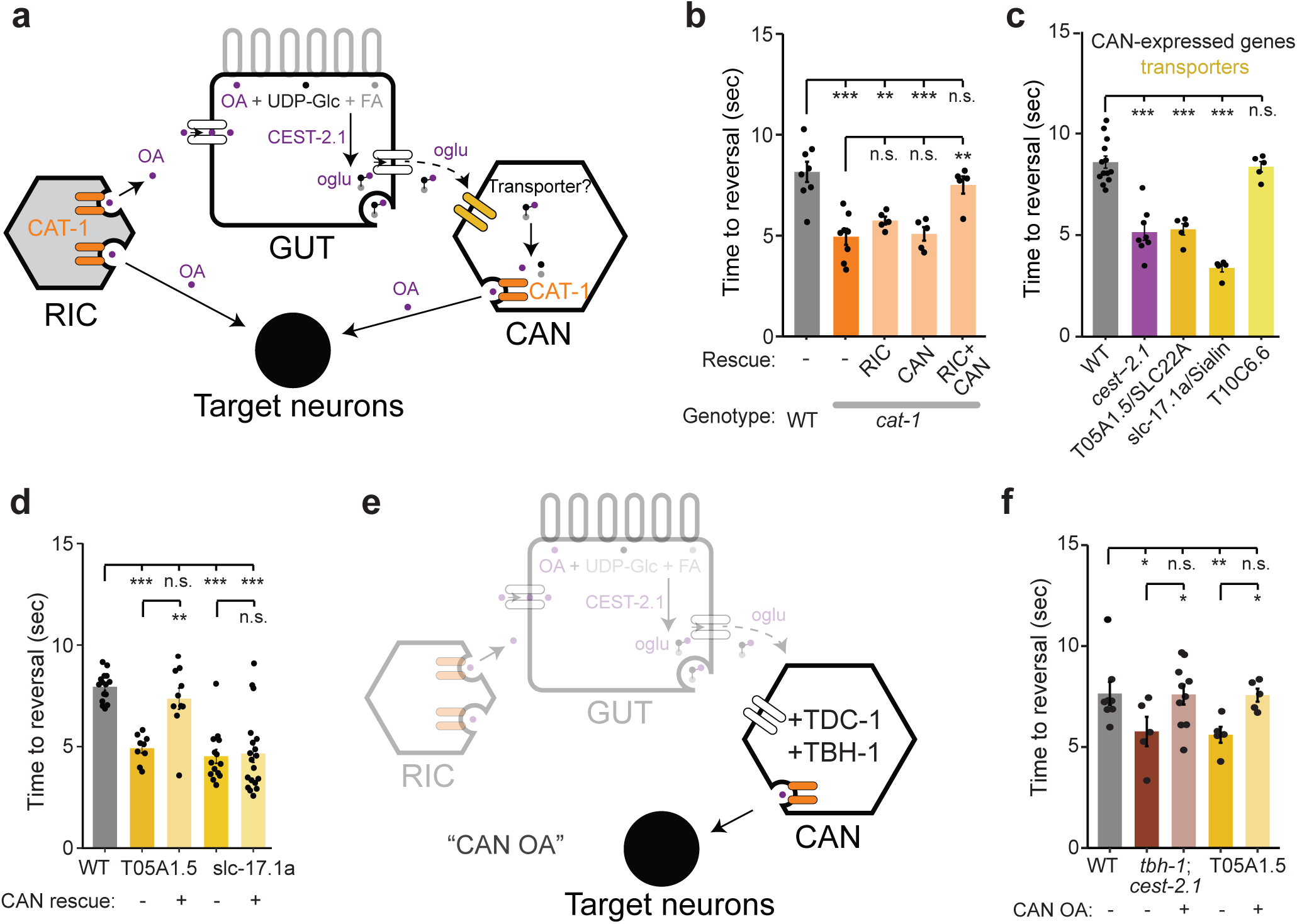
Octopamine is transferred to the CAN cell via glucoside trafficking. **a**, Cartoon depicting hypothetical model of transport of octopamine between tissues. RIC neurons and the gonad sheath cell (not shown) synthesize and release octopamine (purple dots) via the VMAT/CAT-1 transporter (orange). Octopamine glucosides (three-colored ball and stick) are putatively assembled from octopamine, UDP-glucose (black dots) and fatty acids (grey dots) in the intestine prior to release. The CAT-1 expressing CAN neuron may import and subsequently release octopamine glucosides (release not shown) or hydrolyzed octopamine (free purple dots in CAN). Putative transporter(s) for import of octopamine glucosides into CAN are indicated in yellow. **b-d,** SOS reversal latency of animals of the indicated genotypes in response to 33% octanol following 20 minutes of food removal. Each dot indicates an independent assay performed on separate days with at least 20 animals each. Error bars are SEM. Top, *** and **, p<0.001 and p<0.01, relative to WT, respectively. In (**b, d**) bottom **, p<0.01 relative to respective mutant with no rescue transgene. ANOVA with TUKEY post hoc correction. n.s., not significant. **e,** Cartoon depicting experimental approach to bypass RIC and intestinal production of octopamine glucosides, referred to as “CAN OA”. Transgenic expression of biosynthetic enzymes encoded by *tdc-1* and *tbh-1* in CAN should result in exclusive production of octopamine in CAN when introduced in a *tbh-1; cest-2.1* mutant, and CAN intracellular production of octopamine in the putative *T05A1.5* octopamine glucoside transporter mutant. **f,** SOS reversal latency of animals of the indicated genotypes bearing ectopic transgenic expression of *tdc-1* cDNA and *tbh-1* cDNA in the CAN neurons (“CAN OA”, via *ceh-23*p) indicated with (+) or with no transgene (-) in response to 33% octanol following 20 minutes of food removal. Each dot indicates an independent assay performed on separate days with at least 20 animals each. Error bars are SEM. Top, ** and *, p<0.01 and p<0.05 relative to WT, respectively. Bottom, * indicates p<0.05 relative to respective mutants lacking transgene. ANOVA with TUKEY post hoc correction. n.s., not significant.

We found that *cat-1* mutant animals in which *cat-1* is re-expressed in both RIC and CAN, but not in either cell type individually, displayed wild type octanol responses (Fig. 5b). Importantly, because both serotonin and dopamine also require CAT-1 for vesicular loading and release, and neither serotonin or dopamine are produced in RIC or CAN, it is unlikely that these effects are mediated by serotonin or dopamine^49^. These results indicate that CAT-1/VMAT is required in both CAN and RIC for octopamine-dependent behaviors, consistent with a model in which octopamine or octopamine glucosides are imported into and function within the CAN cell.

Our observation that CAN-associated genes are involved in the regulation of *cest-2.1*-dependent phenotypes and metabolism prompted us to hypothesize that intertissue transport of octopamine, via *cest-2.1*-dependent octopamine glucosides, might underlie the requirement for CEST-2.1 in olfactory behavior. As no octopamine reuptake transporters have yet been identified in *C. elegans,* we searched for putative transmembrane transporters that may enable trafficking of octopamine glucosides from the gut to the CAN cell. In animals, ATP-independent transport of small-molecule metabolites across cellular membranes typically occurs via the activity of Major Facilitator Superfamily (MFS) transporters^52^, which includes both plasma membrane and vesicular transporters (such as CAT-1 in the latter category)^49^. Therefore, we also looked for genes encoding MFS transporters that are expressed in the CAN cell. Of 70 genes encoding putative MFS proteins we identified in the *C. elegans* genome, a small subset are likely expressed in CAN^45^ (Table S4). We acquired loss-of-function mutations in 3 of the top 5 most highly CAN-expressed MFS-encoding genes and examined olfactory responses in these mutants to dilute octanol. We found that two of these three mutant strains – bearing mutations in *T05A1.5* and *slc-17.1a*, respectively – exhibited response latency phenotypes similar to *cest-2.1* mutants (Fig. 5c). Olfactory behavioral phenotypes of *T05A1.5* mutants, but not *slc-17.1* mutants, were rescued upon CAN-specific expression (Fig. 5d). *T05A1.5* encodes a member of the Solute carrier 22 A (SLC22A) family, which are typically plasma-membrane localized proteins implicated in intra-and inter-organism metabolite trafficking and communication, and is specifically expressed in the CAN cell^45,52–54^. *slc-17.1* encodes a *C. elegans* orthologue of vertebrate SLC-17 family channels, which are typically vesicular-localized transporters capable of transporting metabolites such as amino acids, nucleotides and sugars and includes the lysosomal sialic acid transporter (Sialin) as well as the vesicular glutamate transporter, and is broadly expressed in *C. elegans*^45,52,55^. To initially ascertain whether T05A1.5 may plausibly transport octopamine or octopamine glucosides into CAN, we performed simulated ligand docking experiments via the recently developed Protenix implementation of Alphafold3^56^. We found that while CAT-1 and the *C. elegans* dopamine transporter, DAT-1, are predicted to accommodate only small monoamines such as octopamine, but not octopamine glucosides, T05A1.5/SLC22 appears to be potentially capable of binding to diverse ligands, including octopamine glucosides (Extended Data Fig. 6a). With the caveat that these are simulations and we are not able to directly visualize octopamine glucoside import into CAN *in vivo*, these results suggest that the CAN cell plays a critical role in octopamine glucoside-dependent olfactory behaviors, likely through intertissue octopamine transport via a CAN-expressed transporter and octopamine release via CAT-1/VMAT.

Our behavioral results suggest that *cat-1*, and by extension octopamine release, is required in both the *tbh-*1-expressing RIC/gonad sheath and CAN. In principle, this could reflect a requirement of octopamine release from both RIC and CAN during this behavior, or alternatively, RIC may supply the necessary octopamine for eventual CAN-mediated release of octopamine or octopamine glucosides. To distinguish between these possibilities, we introduced the biosynthetic enzymes required for octopamine production (TDC-1 and TBH-1) directly into the CAN cells via transgenic expression in *tbh-1; cest-2.1* biosynthesis mutant and T05A1.5/SLC-22A transporter mutant worms (Fig. 5e). We found that ectopic production of octopamine in CAN rescued olfactory phenotypes of both *tbh-1; cest-2.1* and *T05A1.5/SLC-22* mutants, suggesting that CAN-specific free octopamine release, likely via octopamine glucoside import and subsequent hydrolysis, underlies this olfactory behavioral modulation off food (Fig. 5f).

## Discussion

In this work we identified a neurotransmitter trafficking pathway that integrates biochemical input from the regulation of fat metabolism into a neuromodulatory behavioral circuit. Our genetic, behavioral and biochemical evidence suggests that production of agonists of the central regulator of lipid signaling, NHR-49/PPARα, results in two distinct physiological outcomes: alteration of fat metabolism via NHR-49/PPARα^35^ and the modulation of behavior via octopamine signaling (Extended Data Fig. 6b). In the latter case, NHR-49/PPARα agonists are covalently linked to the neurotransmitter octopamine via glucosylation and acylation in the intestine, resulting in the production of lipidated neurohormones. We show that these biochemical transformations are required for the regulation of the effects of octopamine on olfactory behavior and occur via recruitment of additional cells with the capacity to release octopamine upon food removal. These results suggest that internal metabolic state – specifically fat metabolism – is coupled to neurotransmitter-regulated and feeding-state-dependent behaviors via covalent attachment of an NHR-49/PPARα agonist to a neurotransmitter-derived glucoside.

We have shown that the neurotransmitter octopamine is glucosylated and further acylated in the intestine via the carboxyl esterase *cest-2.1*, revealing a multi-tissue octopamine trafficking loop that terminates in the release of free octopamine from CAN cells. This process is necessary for full octopamine-dependent attenuation of olfactory avoidance responses after long-term food removal. Octopamine is synthesized only in two cell types in *C. elegans*: the RIC interneurons and the gonad sheath cells, the latter of which are a layer of epithelial cells that envelope the hermaphrodite germline^20,57,58^. This suggests that octopamine must be transferred between RIC or gonad sheath and the gut. While we do not yet know how octopamine is imported into intestinal cells, we speculate that this likely occurs via a major facilitator superfamily transporter similar to the type we have implicated in the CAN cell. This process of neurotransmitter inactivation is reminiscent of the role of a glia-expressed *Drosophila* non-ribosomal peptide synthetase, encoded by *ebony*, which catalyzes the inactivating linkage of beta-alanine to both dopamine and histamine, the latter of which is hydrolyzed via beta-alanyl-amine hydrolase, encoded by *tan,* resulting in active neurotransmitter^59,60^.

CAN are a peculiar cell type that has no apparent chemical synapses, but nonetheless exhibits some characteristics of neurons, such as long axon-like processes, which form electrical synapses with other neurons. Moreover, while the CAN cell does not appear to express biosynthetic enzymes required for the production of neurotransmitters or biogenic amines, it does, curiously, express a vesicular monoamine transporter^43,50^. Via RNA-sequencing, behavioral analysis and cell-specific rescue, we identified plausible components of the proposed multi-tissue octopamine trafficking loop terminating with octopamine release by the CAN cell. *T05A1.5* encodes an orthologue of human Solute Carrier 22 A transporters, which are implicated in trafficking of organic cations, anions and zwitterions as well as sugars across the plasma membrane^61^. The closest orthologue of T05A1.5 by sequence homology, Organic Anion Transporter 3 (OAT3) has broad substrate specificity, can bind to glucuronide conjugates and is able to transport various endogenous metabolites, as well as drug and gut microbial derived products^62–64^. Members of this transporter family have also been implicated in the uptake of monoamine neurotransmitters^53^. While we cannot directly visualize spatial localization of octopamine and its derivatives *in vivo*, our data suggests that this transporter mediates import of octopamine glucosides into the CAN cell.

How are the *cest-2.1*-dependent octopamine glucosides then processed in the CAN cell? The requirement of a CAN-expressed putative glucoside hydrolase encoded by *gba-4,* coupled with the cell-specific requirement for CAT-1/VMAT in CAN suggests that intracellular hydrolysis of octopamine glucosides may release free octopamine for uptake into CAT-1-positive vesicles prior to exocytosis. Consistent with this model, biosynthetic production of octopamine in CAN is sufficient to bypass the RIC and intestinal components of this pathway. While this model predicts that hydrolysis of inert octopamine glucosides to release their free, active octopamine form occurs intracellularly, this does not preclude the possibility of extracellular hydrolysis of MOGLs.

Somewhat paradoxically, CAN does not appear to have many dense-core vesicles and lacks clear synaptic vesicles, which can be sites of neuronal monoamine release^65,66^. However, VMAT-dependent release of monoamines such as dopamine from somatodendritic compartments via other vesicular pathways has also been described^67^. We speculate that CAN uses a calcium-and VMAT-dependent mechanism to drive octopamine-containing vesicle release, consistent with known release mechanisms of other monoamines^65^. Upon release, the positioning of the CAN cell process may enable octopamine to access different target cells than those accessible to RIC-released octopamine, expanding the spatial and physiological response properties of this neuromodulator.

The nuclear receptor NHR-49/PPARα plays a central role in *C. elegans* lipid metabolism, for example, by regulating the expression of the desaturase FAT-7/SCD1, which catalyzes the biosynthesis of oleic acid which then serves as a precursor for most polyunsaturated fatty acids^33^. We show that octopamine-dependent behaviors depend on biosynthesis of both the endogenous agonist of NHR-49 via the methyl transferase *fcmt-1,* and of dietary cyclopropane fatty acids, followed by attachment of these fatty acyl moieties to octopamine glucosides via CEST-2.1. This direct coupling of lipid homeostasis and neurotransmitter signaling may enable *C. elegans* to flexibly alter behavior concurrent with metabolic changes. For example, expanding the duration and extent of octopamine-modulated olfactory responses via octopamine glucoside signaling may reflect an adaptive behavioral response to extended-nutrient limitation.

Why do *C. elegans* engage with a complex, multitissue pathway as we have described here? Notably, glycosylation of neurotransmitters occurs in many species. For example, glucuronide conjugates of dopamine and serotonin are detectable in the brain of mammals, but they are generally assumed to play a role in detoxification and/or clearance^68–71^. We speculate that neurotransmitter glycosylation and further biochemical modification in the way we describe here might confer two specific advantages. Firstly, inactivating a neurotransmitter via glucosylation likely alters the types of transporters capable of intertissue trafficking while avoiding the risk of inappropriately activating neurotransmitter receptors during transit, and secondly, as we have shown here, glucosylation may enable further modification to integrate signals from other metabolic pathways. Adaptive behavioral responses to feeding state and nutrient status often involve complex multisensory integration events^72,73^. Our results suggest that covalent incorporation of chemical signals that depend on lipid homeostasis may enable balancing foraging costs and benefits, by temporally coordinating both lipid physiological and behavioral responses.

## Methods

### Strains

*C. elegans:* All worms were maintained on Nematode Growth Media (NGM) at 20 °C. For all behavioral assays, animals were maintained on these plates for a single generation by transferring 6-10 adult worms or 40-50 eggs or 30-40 L4 worms at 20°C. A list of *C. elegans* and bacteria strains used in this work is provided in Table S5. Multiple independent transgenic lines were used in all experiments.

*Bacteria:* All bacterial strains were streaked from glycerol stocks onto LB media, containing an antibiotic when appropriate, within 4 weeks of culture. P1*vir* phage transduction was performed as described^74^. Briefly, 100µL of P1 phage supernatant was used to infect 5mL culture of the donor strain in LB media + 0.2% glucose and 5mM CaCl_2_ and grown at 37°C with shaking for approximately 3hr until visible lysis was apparent. Phage supernatant from donor strain was extracted by adding 200µL CHCl_3_ followed by centrifugation for 5 mins at ∼9200 x g, 4°C to recover bacteria-free supernatant, which was stored at 4°C prior to transduction after confirming the absence of viable bacterial cells via plating. 2µL of this supernatant was used to infect a resuspended pellet from a 1.5mL culture of overnight-grown OP50 cells, resuspended in P1 salt solution (10mM MgCl_2_, 5mM CaCl_2_) for 30 min at 37°C. Calcium was chelated via addition of 1ml of LB + 200 µl of 1M sodium citrate followed by incubation at 37°C for 1 hr. Cells were then pelleted at maximum speed, followed by resuspension in 100 µL LB prior to plating on LB + 50mg/mL Kanamycin + 5mM sodium citrate. Transduced deletions were confirmed by PCR. For seeding of bacteria onto NGM media, 200 µL of a saturated overnight culture grown overnight at 37°C and derived from a single colony grown in LB broth was added to 6 cm petri dishes and allowed to grow for 2-5 days prior to worm culture.

### Behavior

*Smell on a stick (SOS):* SOS assays in response to 1-octanol were performed as previously described^22,26^. NGM plates were pre-dried overnight before assays. Age-matched young adult worms were picked from food to a clean transfer plate and allowed to briefly crawl away from food for approximately 1 min.

For experiment involving worms grown on *E. coli* cfa mutant bacteria, animals were maintained on *cfa* mutant *E. coli* for at least three generations before behavioral assays. With the exception of Extended Data Fig. 1a, worms were transferred to a clean NGM plate for a total of 20 min from the time the first worm was transferred from food before assaying responses to 33% octanol. Thirty-three per cent octanol was prepared immediately before the assay by dilution in 200-proof ethanol (Acros Organics 61509-0010).

### Molecular biology

*Cloning:* Plasmids used in all experiments are listed in Table S6. All vector maps are available on Github. *Microinjection:* All microinjections were performed using standard procedures^75^.

Extrachromosomal arrays were established by injection of 100ng/µl total DNA containing transgene plasmids, co-injection marker, and DNA ladder (Invitrogen 10787018, see Table S5). Targeted multicopy transgene insertion was performed using the FLInt method^76,77^

### RNA sequencing

Total RNA was isolated from 3 biologically independent, well-fed mixed-stage cultures of WT and *cest-1.2* mutant worms using QIAzol lysis reagent (QIAGEN sciences). mRNA was isolated using poly A selection and sequenced with paired-end Illumina by the Yale Center for Genome Analysis (YCGA). Trimmed reads were quantified using Kallisto v.0.48^78^. Differential expression analysis was performed using Sleuth^79^.

1. *E. coli suspension for nematode cultures*.
2. *E. coli* was streaked on LB agar plates and grown at 37 °C for 20 hours. Plates were then sealed and stored at 4 °C for up to 4 weeks. Liquid cultures were generated by inoculating a single colony into 125 mL Erlenmeyer flasks containing 30 mL LB. Cultures were grown at 37 °C shaking at 200 RPM for 20 hours. 10 mL of this culture was diluted in 2 L LB in a 4 L Erlenmeyer flask, and the resultant culture was grown at 37 °C shaking at 180 RPM for 20 hours. Bacteria were collected by centrifugation (2500xg, 4 °C, 20 min), supernatant was removed, and the bacterial pellet was resuspended in M9 solution to yield a 1 g/mL suspension of bacteria in M9.
3. *C. elegans liquid cultures*.

6 cm maintenance plates of *C. elegans* were transferred to 10 cm diameter petri dish plates seeded with *E. coli* OP50 and grown to confluence. Confluent plates were washed with 5 mL S-complete medium into a 125 mL Erlenmeyer flask with 20 mL S-complete medium and fed with 2 mL of 1 g/mL *E. coli* suspension in M9. Mixed stage cultures were grown for 72 hours at 20 °C shaking at 180 RPM. After 72 hours, mixed stage cultures were pelleted, washed with 25 mL water, and treated with alkaline bleach to yield a suspension of eggs, which were washed two times with 25 mL M9 buffer and rocked overnight in 5 mL M9 solution to yield synchronized, starved L1 larvae. Cultures of synchronized adults were prepared by adding 70,000 synchronized L1 larvae, to 25 mL of S-complete medium in a 125 mL Erlenmeyer flask to which was added 2 mL of 1 g/mL *E. coli* suspension in M9 solution. Cultures were incubated at 20 °C shaking at 180 RPM for 60-63 hours until animals were gravid.

### C. elegans liquid culture harvest and sample preparation for HPLC-HRMS

Cultures of 70,000 synchronized gravid adult worms were transferred to 50 mL falcon tubes and settled for ten minutes. 15 mL of supernatant was transferred to a separate 50 mL falcon tube, snap frozen in liquid nitrogen, and lyophilized to dryness (exo-metabolome). The remaining worm pellet was suspended in 25 mL M9 solution and allowed to settle. This washing step was repeated two more times with M9 solution and the resultant pellet of gravid adult worms was snap frozen in liquid nitrogen. Frozen samples were lyophilized to dryness using a VirTis BenchTop 4 K Freeze Dryer. The resultant powder was suspended in 10 mL methanol and samples were allowed to rock at room temperature for 18 hours. After extraction, samples were centrifuged (2500Xg, 10 min, 20 °C), the clarified extract was transferred to a 20 mL scintillation vial, and samples were concentrated to dryness in an SC250EXP Speedvac Concentrator coupled to an RVT5105 Refrigerated Vapor Trap (Thermo Scientific). The resulting powder was suspended in 1 mL of methanol by vigorous vortex and 10 minutes of sonication (Mettler Electronics Cavitator ultrasonic cleaner). The resulting suspension was transferred to a 1 mL Eppendorf tube, centrifuged (10,000Xg, 5 min, 20 °C) in an Eppendorf 5417 R centrifuge and the clarified extract was transferred to an HPLC vial (Thermo Scientific cat. no. 6PSV9-1P). Samples were concentrated to dryness as above, suspended in 100 µL of methanol with vigorous vortex, transferred to a 1 mL Eppendorf tube, centrifuged (18,000Xg, 10 min, 20 °C) in an Eppendorf 5417 R centrifuge, and 70 µL was transferred to HPLC vial inserts (SureSTART 6PME03C1SSP). Samples were stored at -20 °C until analysis.

### Mass spectrometric analysis

HPLC−HRMS analysis was performed on a ThermoFisher Scientific Vanquish Horizon UHPLC System controlled by Chromelion software (Thermo Fisher Scientific) coupled with a Thermo Q Exactive HF hybrid quadrupole-orbitrap high-resolution mass spectrometer, controlled by the same software, and equipped with a HESI ion source. Metabolites were separated using a water-acetonitrile gradient on an Agilent Zorbax Eclipse XDB-C18 column (150 mm × 2.1 mm, particle size 1.8 μm) maintained at 40 °C. Solvent A: 0.1% formic acid in water; Solvent B: 0.1% formic acid in acetonitrile. A/B gradient started at 1% B for 5 min after injection and increased linearly to 99% B at 20 min, using a flow rate 0.5 mL/min. Mass spectrometer parameters: spray voltage 3.0 kV, capillary temperature 380 °C, probe heater temperature 300°C; sheath, auxiliary, and spare gas 60, 20, and 2, respectively; S-lens RF level 50, resolution 240,000 at *m/z* 200, AGC target 3×10^6^. The instrument was calibrated with positive and negative ion calibration solutions (ThermoFisher). Each sample was analyzed in positive and negative modes using an *m/z* range 70 to 1000. Tandem mass spectrometry parameters: Resolution 45,000, AGC target 5×10^4^, Maximum IT 80 ms, loop count 10, TopN 10, isolation window 1.0 *m/z*, stepped NCE 10 to 30, minimum AGC target 8×10^3^, intensity threshold 1×10^5^, no apex trigger, no charge exclusion, no peptide match, exclude isotopes on, dynamic exclusion 1.0 s, if idle, pick others. HPLC-HRMS RAW data were converted into mzXML file format using MSConvert (v.3.0, ProteoWizard) and were analyzed using Metaboseek v.0.9.9^30^.

### Statistical analysis

All statistical analyses were performed in R (https://www.R-project.org/) and RStudio (http://www.rstudio.com). For behavioral experiments, P-values and n indicated in figures are derived from comparison of sample means calculated for each day of behavior, which typically included measurements of ∼ 20 animals per sample. Statistical comparisons, raw data and code to replicate these analyses are available on Github. Sample sizes were chosen according to conventional estimates of power for all assays. Samples were blinded prior to behavior analysis.

## Acknowledgements

We thank Piali Sengupta, Inna Nechipurenko and members of the O’Donnell and Schroeder labs for critical comments on the manuscript; Cori Bargmann and David Breslow for helpful suggestions; the Caenorhabditis Genetics Center for *C. elegans* strains; This work was partly supported by the NIH (DP2 GM154014 to M.P.O; R35 GM131877 to F.C.S).

## Author contributions

M.P.O. and F.C.S. supervised the study. P.M. and T.B. carried out behavioral experiments and analyzed all behavioral data. A.F.S. carried out metabolomics and chemical synthesis and analyzed most of the metabolomics data. P.M., T.B., N.M. and M.D. generated molecular reagents and strains. C.J.W. assisted with metabolomic analyses. T.B. and M.D. performed RNA sequencing experiments. M.P.O., A.F.S., T.B., P.M. and F.C.S. analyzed data and wrote the manuscript with input from all co-authors.

## Data availability

Raw sequencing reads will be uploaded to the Sequence Read Archive (SRA), and MS data for all metabolome samples analyzed in this study will be uploaded to the GNPS Web site (massive.ucsd.edu) and will be made publicly available upon publication.

---

**Extended Data Fig. 1.**
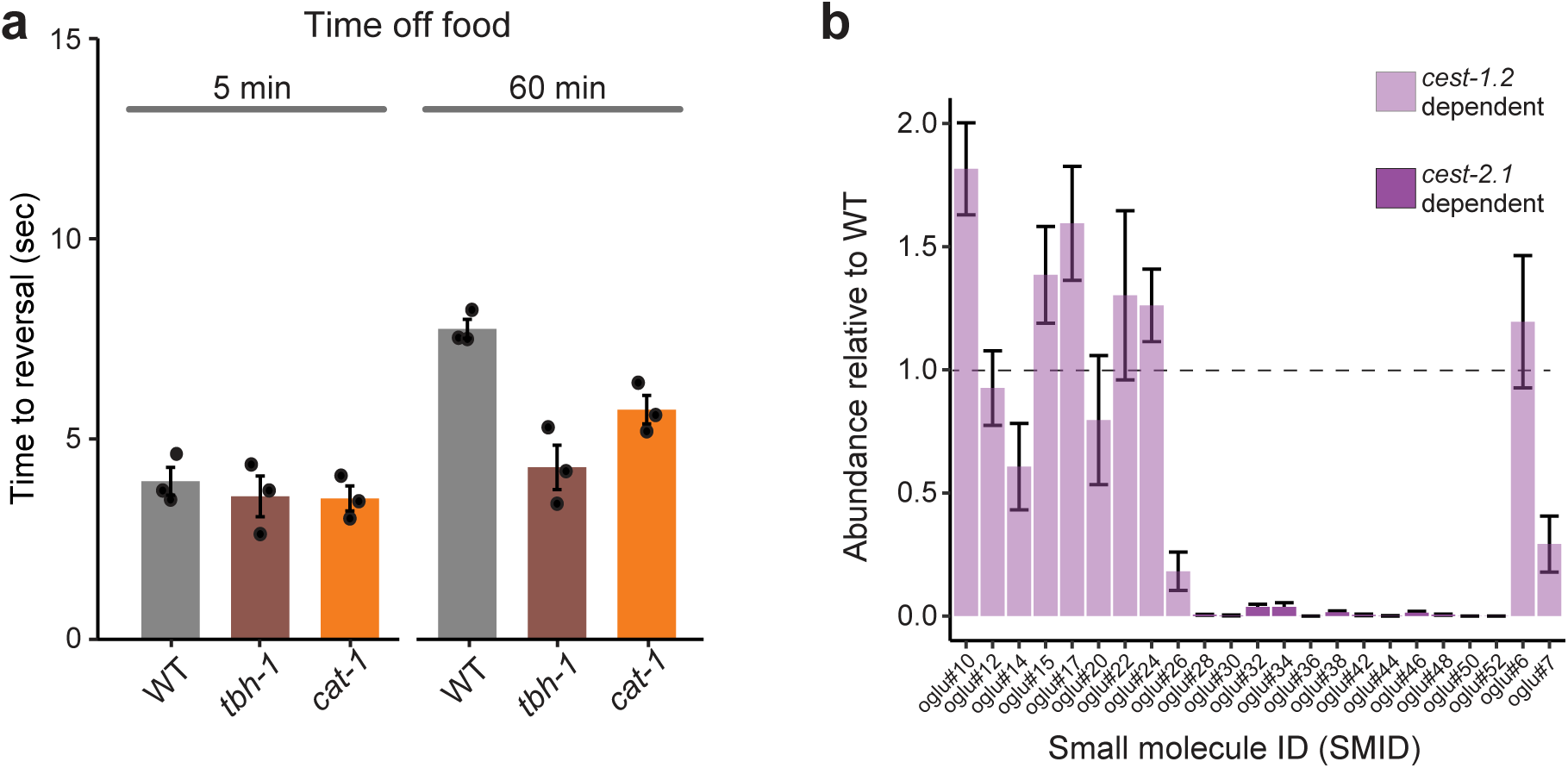
SOS reversal latency and octopamine glucoside profiles of the indicated genotypes. **a,** SOS reversal latency of animals of the indicated genotypes in response to 33% octanol following removal from food for the indicated durations. Each dot indicates an independent assay performed on separate days with at least 20 animals each. Error bars are SEM. **b,** Profile of octopamine glucoside abundance in *cest-2.1* mutant animals relative to WT animals. *Cest-1.2*-dependent octopamine glucosides are shown in light purple, *cest-2.1* dependent octopamine glucosides are shown in dark purple. Error bars are SEM of 4 independent replicates.

**Extended Data Fig. 2.**
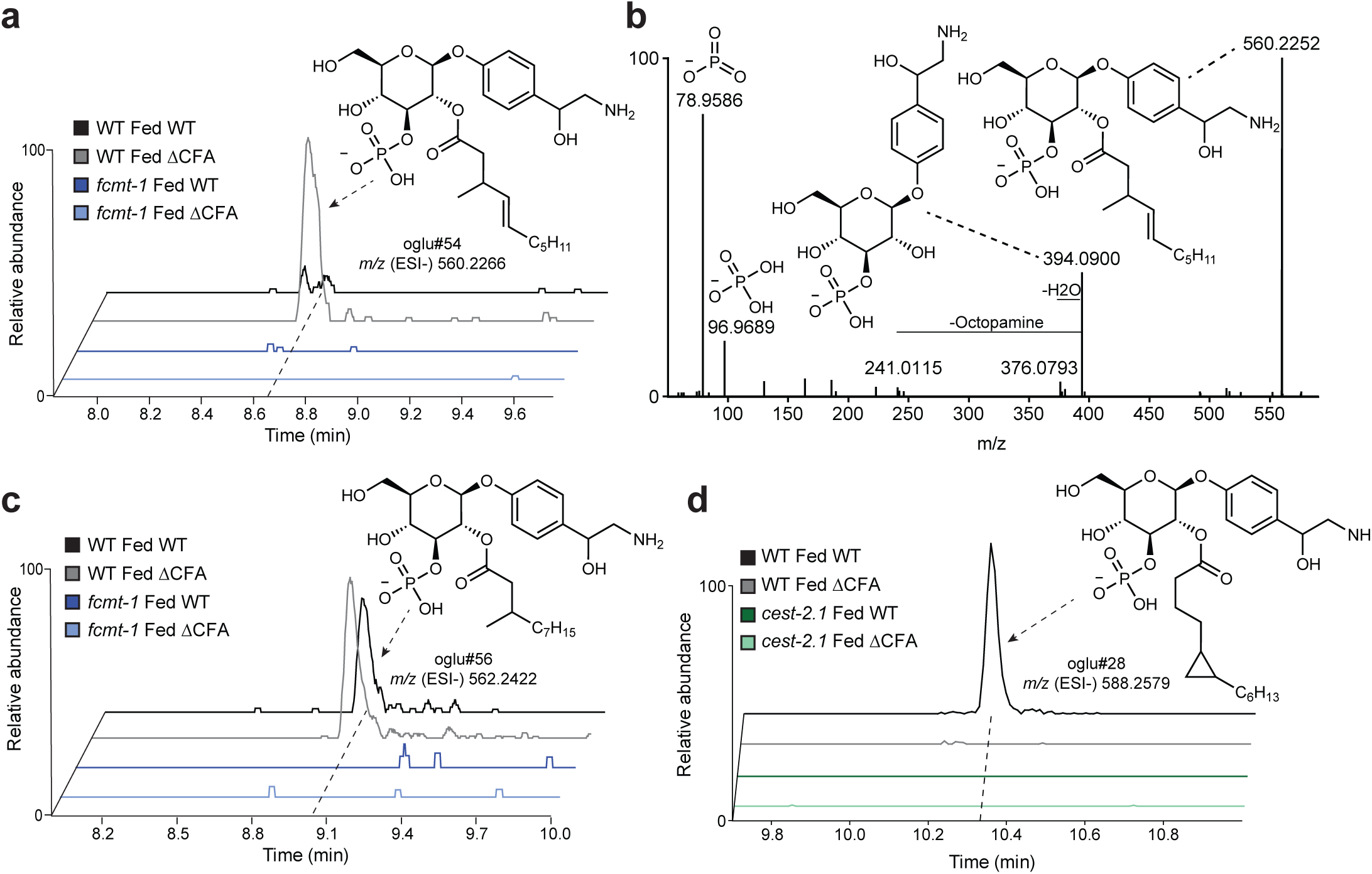
Dependence of octopamine glucoside production on different bacterial diets. **a, c,** Ion chromatograms of oglu#54 (**a**) or oglu#56 (**c**) in WT *C. elegans* fed WT *E. coli* (black), WT *C. elegans* fed *Δcfa E. coli* (gray), *fcmt-1* mutant *C. elegans* fed WT *E. coli* (blue) and *fcmt-1* mutant C. elegans fed *Δcfa E. coli* (light blue). **b,** Annotated MS^2^ fragmentation spectrum of oglu#54. **d,** Ion chromatograms of oglu#28 in WT *C. elegans* fed WT *E. coli* (black), WT *C. elegans* fed *Δcfa E. coli* (gray), *cest-2.1* mutant *C. elegans* fed WT *E. coli* (green) and *cest-2.1* mutant *C. elegans* fed *Δcfa E. coli* (light green).

**Extended Data Fig. 3.**
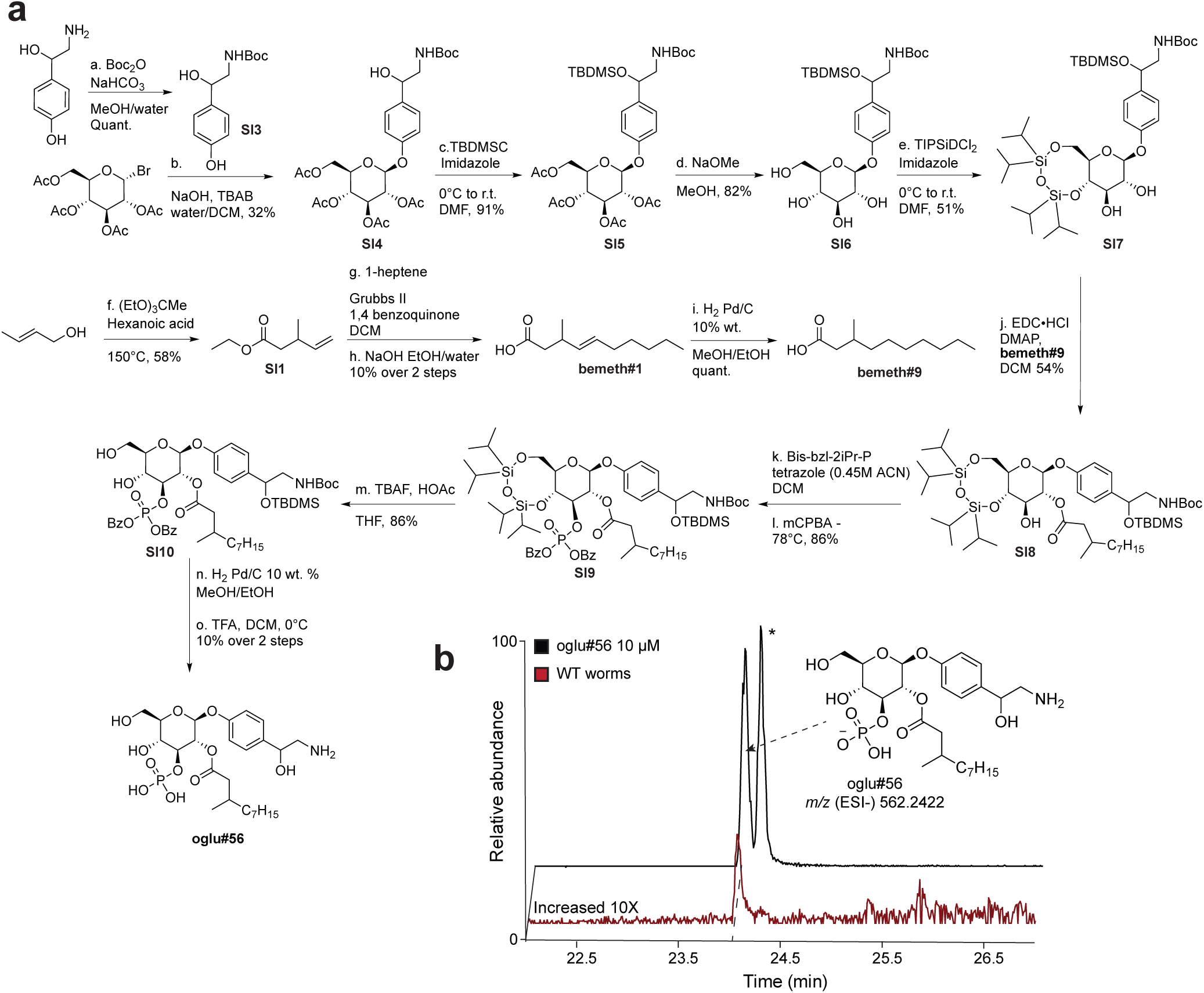
Confirmation of the structure of oglu#56 via total synthesis. **a,** Synthetic route to oglu#56**. b,** Ion chromatograms of a synthetic standard of oglu#56 (black), and oglu#56 in WT *C. elegans* endo metabolome (maroon). * indicates non-biologically relevant diastereomer.

**Extended Data Fig. 4.**
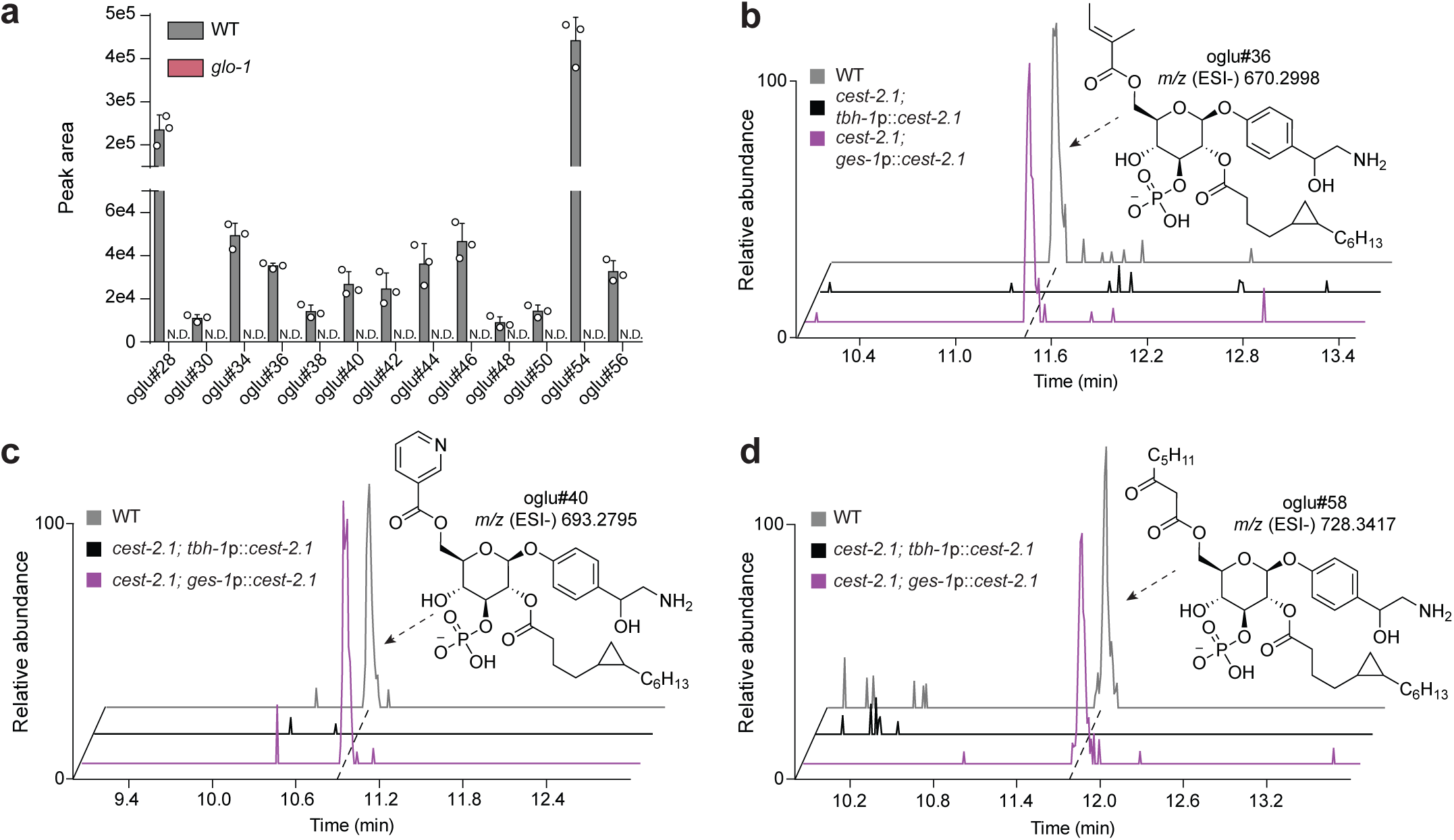
Octopamine glucoside production in different mutants. **a,** Relative abundance of the indicated octopamine glucosides in WT animals (grey bars) or *glo-1* mutants identified via HPLC-MS. Each dot indicates the peak area of an individual metabolite in independent samples. Error bars are SEM. **b-d,** Ion chromatograms of oglu#36 (**b**), oglu#40 (**c**), or oglu#58 (**d**) in WT C. elegans (grey), *cest-2.1* mutant *C. elegans* with *cest-2.1* expressed in RIC and gonad sheath tissue (*tbh-1*p::*cest-2.1*, black), and *cest-2.1* mutant *C. elegans* with *cest-2.1* expressed in intestinal tissue (*ges-1*p::*cest-2.1*, purple).

**Extended Data Fig. 5.**
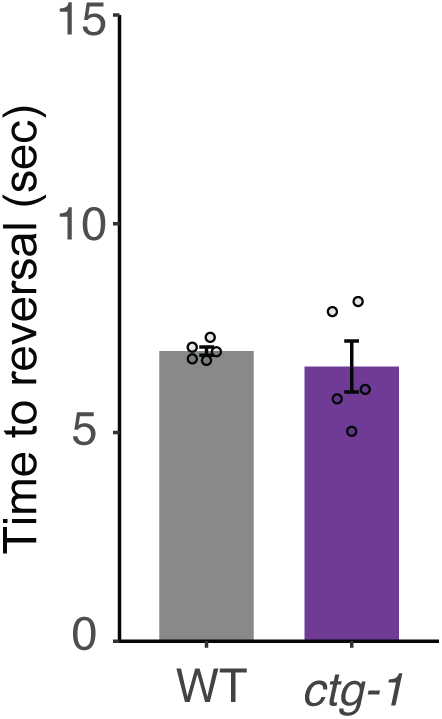
SOS reversal latency is not affected by loss of *ctg-1*. SOS reversal latency of animals of the indicated genotypes in response to 33% octanol following removal from food for 20 minutes. Each dot indicates an independent assay performed on separate days with at least 20 animals each. Error bars are SEM.

**Extended Data Fig. 6.**
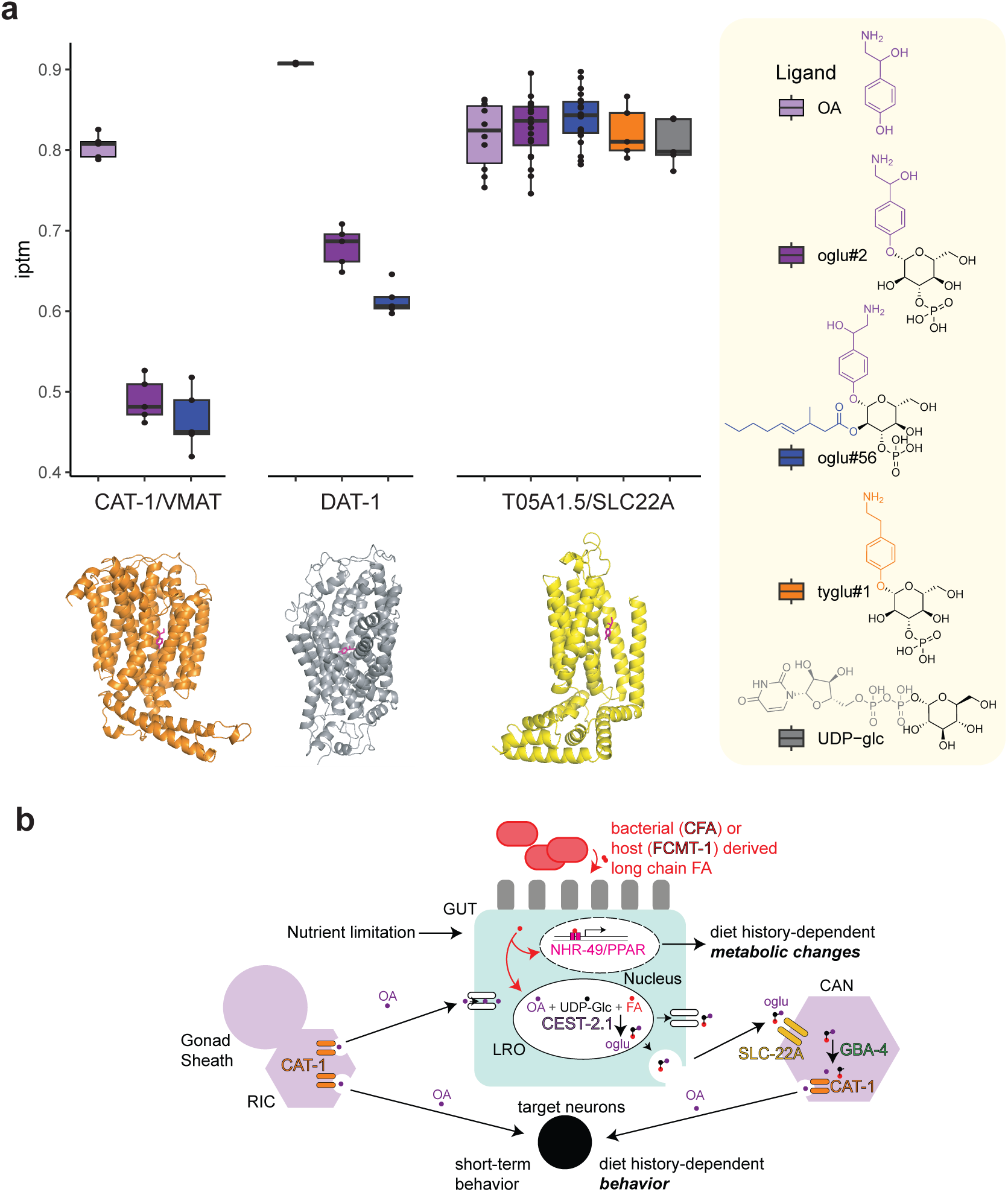
Docking of octopamine derivatives and model for octopamine intertissue transport. **a,** Simulated ligand docking of the indicated ligands to the indicated transporters via Protenix Alphafold3 reproduction. Dots indicate iPTM values for individual ligand docking replicate simulations, which are summarized in colored boxplots. **b,** Cartoon depicting hypothetical model of transport of octopamine between tissues. Proteins encoded by genes identified/implicated in this study are colored with black outline. Putative pathway proceeds from left. RIC neurons and the gonad sheath cell (purple) synthesize and release octopamine (purple dots) via the VMAT/CAT-1 transporter (orange ovals). Octopamine is imported into the intestine via unidentified transporter(s). Nutrient limitation leads to accumulation of β-methyl and cfa-containing medium chain fatty acids (red dots), which are incorporated into octopamine glucosides (three-colored ball and stick) via the activity of CEST-2.1 in intestinal LRO. Octopamine glucosides are released via unknown mechanisms and are imported into the CAN cell putatively via the SLC22A/T05A1.5 transporter (yellow ovals). Once in CAN, octopamine glucosides may be hydrolyzed by GBA-4 resulting in free octopamine which is imported into secretory organelles via VMAT/CAT-1. Release of octopamine from CAN leads to increased reversal latency in animals removed from food.

