## supplemental information for "Nuclear receptor-neurotransmitter coupling links behavior to metabolic state"

### 1. Synthetic procedures

#### 1.1 General synthetic methods

All oxygen and moisture-sensitive reactions were carried out under argon atmosphere in flame-dried glassware. Solutions and solvents sensitive to moisture and oxygen were transferred via standard syringe and cannula techniques. Reactions were cooled with ice water to 0 °C, with dry ice and acetone to -78 °C, or heated with mineral oil baths depending on reaction temperature. Mixtures (reaction or from chromatography) were concentrated using a Buchi rotary evaporator. Di-*tert*-Butyl decarbonate ($Boc_2O$), sodium hydroxide (NaOH), *tert*-Butyldimethylsilyl chloride (TBDMSCl), imidazole, 1,3-dichloro-1,1,3,3-tetraisopropyldisiloxane ($TIPSiDCl_2$), triethyl orthoacetate ($(EtO)_3CMe$), hexanoic acid, Grubbs II, 1,4 benzoquinone, palladium on carbon 10 wt. % (Pd/C), dibenzyl *N*,*N*-diisopropylphosphoramidite (bis-bzl-2iPr-P), and 3-chloroperbenzoic acid (mCPBA) were purchased from Sigma Aldrich. 4-(*N,N*-dimethyl)aminopyridine (DMAP), *N*-(3-dimethylaminopropyl)-*N'*-ethylcarbodiimide (EDC), Tetrabutylammonium bromide (TBAB), and 1-heptene were purchased from TCI. Sodium methoxide (NaOME) was purchased from Acros Organics. Tetrazole solution 0.45 M in acetonitrile was purchased from Honeywell Fluka. Trifluoroacetic acid (TFA) was purchased from VWR. Tetrabutylammonium fluoride (TBAF), glacial acetic acid (HOAc), acetonitrile (ACN), water ($H_2O$), dichloromethane (DCM), ethyl acetate (EtOAc), hexanes, methanol (MeOH), dimethylformamide (DMF), were purchased form Fisher Scientific. Thin layer chromatography (TLC) was performed using J. T. Baker Silica Gel IB2F plated with analysis via UV and *p*-anisaldehyde, phosphomolybdic acid, and potassium permanganate stains. Flash chromatography was performed using Teledyne Isco CombiFlash systems and Teledyne Isco RediSep Rf silica and C18 columns. All deuterated solvents were purchased from Cambridge Isotopes. Nuclear Magnetic Resonance (NMR) spectra were recorded on Varian INOVA 600 (600 MHz) or Bruker (500 MHz) AVIII with BBO Prodigy cryoprobe spectrometer at Cornell University's NMR facility. $^{1}H$ NMR chemical shifts are reported in ppm (δ) relative to residual solvent peaks (3.31 ppm for methanol-$d_4$, 4.79 ppm for $D_2O$, and 7.26 ppm for chloroform-*d*). $^{13}C$ NMR chemical shifts are reported in ppm (δ) relative to residual solvent peaks (49.00 ppm for methanol-$d_4$, 49.00 ppm for methanol signal for $D_2O$, and 77.16 ppm for chloroform-*d*). All NMR data processing was done using MNOVA 15.0.0 (https://mestrelab.com/).

### 1.2 Synthesis of bemeth#9

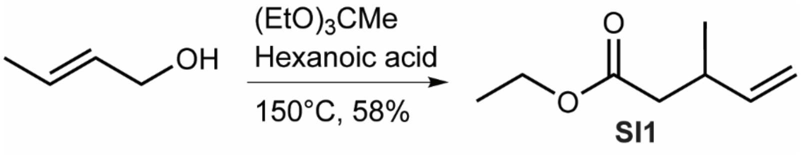

**Ethyl 3-methylpent-4-enoate (SI1).** Prepared as described previously[1]. Hexanoic acid (111 µL, 0.885 mmol, 0.04 eq), triethyl orthoacetate (8.6 mL, 46.91 mmol, 2.28 eq) and crotyl alcohol (1.75 mL, 20.5 mmol, 1eq) were stirred in a round bottom flask fit with a distillation arm. The reaction mixture was heated to 150 °C and was monitored for ethanol distillation. After 2.5 hours, another 50 µLof hexanoic acid (0.4 mmol, 0.02 eq) was added and the reaction was left at 150 °C for 18 hours. The reaction was cooled to room temperature, diluted in hexanes and extracted with water three times. The collected organic phase was washed with saturated aqueous sodium bicarbonate, dried over magnesium sulfate, and concentrated *in vacuo*. Flash column chromatography on silica using a gradient of 0-30% EtOAc/Hexanes afforded **SI1** (**SI1**, 1.951 g, 13.729 mmol, 66.9%). Product was used without further purification.

**$^1$H NMR (500 MHz, chloroform-*d*):** δ (ppm) 5.79 (m, 1H), 5.05 (dt, 1H), 4.98 (dt, 1H), 4.15 (q, 2H), 2.70 (m, 1H), 2.38 – 2.25 (m, 2H), 1.27 (t, 3H), 1.08 (d, 3H).

**$^{13}$C NMR (126 MHz, chloroform-*d*):** δ (ppm) 172.73, 142.69, 113.47, 60.41, 41.54, 34.62, 19.87, 14.46.

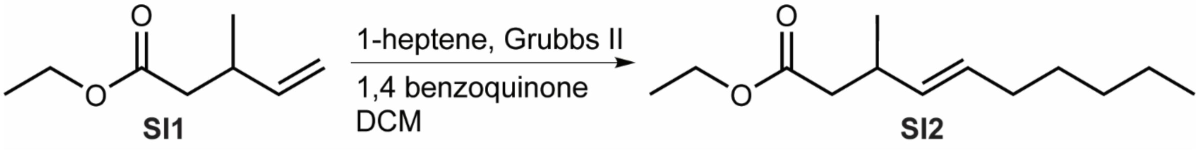

**Ethyl (*E*)-3-methyldec-4-enoate (SI2).** To a stirred solution of **SI1** (400 mg, 2.81 mmol, 1 eq) and 1-heptene (396 µL, 2.81 mmol, 1eq) in DCM (17.58 mL) was added Grubbs catalyst second generation (238 mg, 0.281 mmol, 0.1 eq) and 1,4 benzoquinone (15 mg, 0.05 mmol, 0.05 eq). The reaction was stirred at room temperature for three hours then concentrated *in vacuo*. Flash column chromatography on silica using a gradient of 0-20% EtOAc/Hexanes afforded **SI2** (**SI2,** 277 mg, 1.30 mmol, 46.4%).

**$^1$H NMR (500 MHz, chloroform-*d*):** δ (ppm) 5.42 (m, 1H), 5.31 (m, 1H), 4.11 (q, 2H), 2.62 (p, 1H), 2.31 – 2.20 (m, 2H), 1.95 (q, 2H), 1.35 – 1.21 (m, 9H), 1.02 (d, 3H), 0.88 (t, 3H).

**$^{13}$C NMR (126 MHz, chloroform-*d*):** δ (ppm) 172.94, 134.15, 129.80, 60.30, 42.28, 33.92, 32.62, 31.50, 29.36, 22.71, 20.65, 14.48, 14.25.

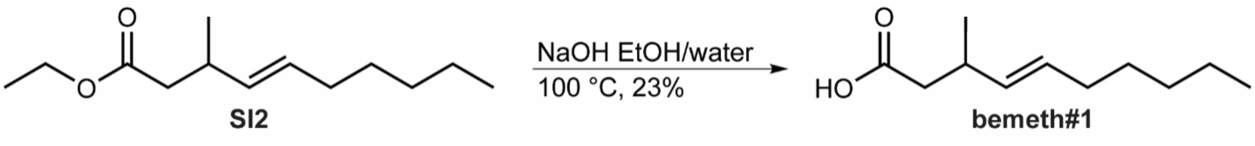

**bemeth#1, (1).** To a stirred solution of **SI2** (270 mg, 1.27 mmol, 1 eq) in a 2:1 mixture of EtOH:$H_2O$ (2.54 mL) was added NaOH (70 mg, 1.52 mmol, 1.2 eq). The reaction was refluxed for one hour. After one hour, the EtOH was removed *in vacuo.* and the remaining aqueous mixture was acidified to pH = 3 with 1M HCl. The aqueous phase was extracted three times with EtOAc, the combined organic phase was washed with brine, dried over magnesium sulfate, and concentrated *in vacuo*. Flash column chromatography on silica using a gradient of 0-30% EtOAc/Hexanes afforded **bemeth#1** (**bemeth#1,** 54 mg, 0.293 mmol, 23%).

**$^{1}$H NMR (500 MHz, chloroform-*d*):** δ (ppm) 5.45 (m, 1H), 5.34 (m, 1H), 2.63 (m, 1H), 2.37 – 2.26 (m, 2H), 1.96 (q, 2H), 1.37 – 1.20 (m, 8H), 1.06 (d, 3H), 0.88 (t, 3H).

**$^{13}$C NMR (126 MHz, chloroform-*d*):** δ (ppm) 177.65, 133.79, 130.14, 41.69, 33.60, 32.61, 31.49, 29.31, 22.70, 20.59, 14.24.

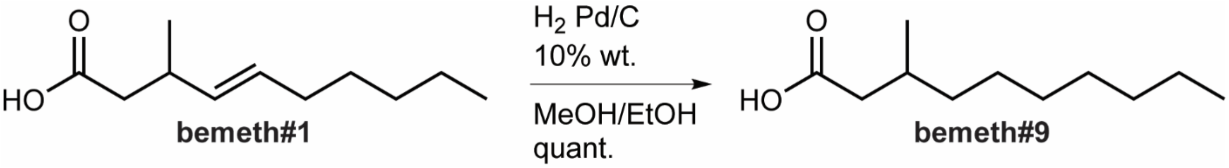

**bemeth#9, (2).** To a stirred solution of bemeth#1 **(1)** (50 mg, 0.271 mmol, 1 eq) in a 1:1 mixture of MeOH:EtOH (16 mL) was added Pd/C 10 wt. % (25 mg). The reaction was flushed with argon for five minutes followed by $H_2$ for two hours. The reaction mixture was then filtered through a pad of celite, and concentrated *in vacuo*. Flash column chromatography on silica using a gradient of 0-20% EtOAc/Hexanes afforded **bemeth#9** (**bemeth#9**, 50 mg, 0.268 mmol, 99%).

**$^{1}$H NMR (500 MHz, chloroform-*d*):** δ (ppm) 2.35 (m, 1H), 2.14 (m, 1H), 1.95 (m, 1H), 1.36 – 1.17 (m, 13H), 0.96 (d, 3H), 0.88 (t, 3H).

**$^{13}$C NMR (126 MHz, chloroform-*d*):** δ (ppm) 178.76, 41.58, 36.85, 32.04, 30.36, 29.87, 29.47, 27.08, 22.86, 19.88, 14.29.

#### 1.3 Synthesis of oglu#56

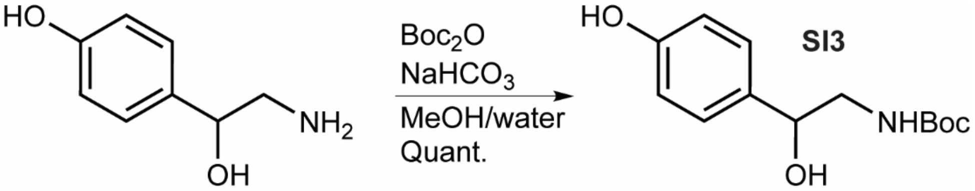

***tert*-Butyl (2-hydroxy-2-(4-hydroxyphenyl)ethyl)carbamate, (SI3).** To a stirred solution of octopamine (189 mg, 1 mmol, 1 eq) in a mixture of 2:1 MeOH:$H_2O$ (6 mL) was added di-*tert*-Butyl decarbonate (327 mg, 1.5 mmol, 1.5 eq) and sodium bicarbonate (252 mg, 3 mmol, 3 eq) at room temperature. After two hours and forty minutes the reaction was diluted in $H_2O$ and extracted with EtOAc three times. The combined organic phase was dried over sodium sulfate and concentrated *in vacuo*. Flash column chromatography on silica using a gradient of 20-40% EtOAc/Hexanes afforded **SI3** (**SI3**, 209 mg, 0.825 mmol, 82.5%).

**$^1$H NMR (500 MHz, methanol-*d4*):** δ (ppm) 7.18 (m, 2H), 6.75 (m, 2H), 4.61 (dd, 1H), 3.21 (m, 2H), 1.42 (s, 9H).

**$^{13}$C NMR (126 MHz, methanol-*d4*):** δ (ppm) 158.56, 158.04, 134.68, 128.46, 116.03, 80.14, 73.75, 49.28, 28.73.

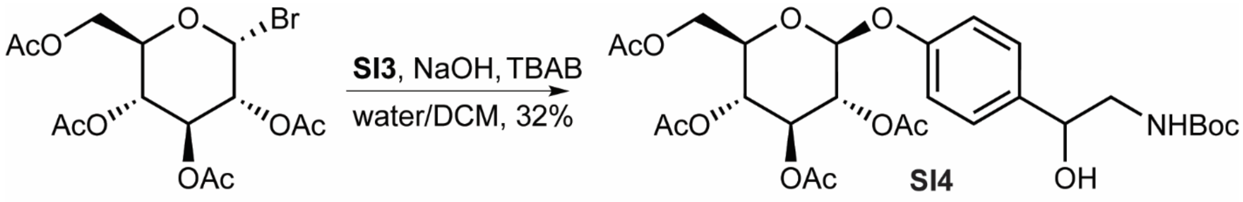

**(2*R*,3*R*,4*S*,5*R*,6*S*)-2-(acetoxymethyl)-6-(4-(2-((*tert*-butoxycarbonyl)amino)-1-hydroxyethyl)-phenoxy)tetrahydro-2H-pyran-3,4,5-triyl triacetate, (SI4).** NaOH (206 mg, 5.17 mmol, 1.2 eq) and tetrabutylammonium bromide (69 mg, 0.215 mmol, 0.05 eq) were solubilized in $H_2O$ (8.9 mL), **SI3** (1.115 g, 4.73 mmol, 1.1 eq) was added and the reaction was left to stir at room temperature for 15 minutes. Bromo-α-D-glucose tetraacetate (1.75 g, 4.3 mmol, 1 eq) was added dropwise as a solution in DCM (17.9 mL). The reaction was stirred vigorously for 24 hours at room temperature then diluted in water and extracted into DCM three times. The combined organic layer was washed with brine, dried over magnesium sulfate, and concentrated *in vacuo*. Flash column chromatography on silica using a gradient of 20-70% EtOAc/Hexanes afforded **SI4** (**SI4**, 734 mg, 1.25 mmol, 29.2%).

**$^1$H NMR (500 MHz, chloroform-*d*):** δ (ppm) 7.3 (m, 2H), 6.97 (m, 2H), 5.28 (m, 2H), 5.16 (t, 1H), 5.07 (m, 1H), 4.91 (s broad, 1H), 4.80 (m, 1H), 4.28 (m, 1H), 4.17 (m, 1H), 3.86 (m, 1H), 3.44 (m, 1H), 3.22 (m, 1H), 2.08 – 2.03 (m, 14H), 1.45 (s, 9H).

**$^{13}$C NMR (126 MHz, chloroform-*d*): δ (ppm)** 170.77, 170.44, 169.60, 169.49, 156.61, 137.03, 127.37, 117.17, 117.11, 99.31, 77.40, 73.72, 72.88, 72.25, 71.33, 68.45, 62.15, 48.64, 28.54, 21.24, 20.91, 20.83, 20.81, 20.79.

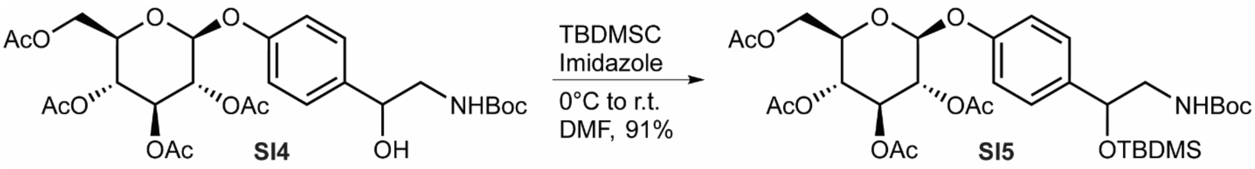

**(2*R*,3*R*,4*S*,5*R*,6*S*)-2-(acetoxymethyl)-6-(4-(2,2,3,3,10,10-hexamethyl-8-oxo-4,9-dioxa-7-aza-3-silaundecan-5-yl)phenoxy)tetrahydro-2H-pyran-3,4,5-triyl triacetate (SI5).** To a stirred solution of **SI4** (730 mg, 1.25 mmol, 1 eq) in DMF (2.5 mL) at 0 °C was added imidazole (170 mg 2.5 mmol, 2.5 eq). TBDMSCl (226 mg, 1.5 mmol, 1.2 eq) was added slowly and the reaction was left to warm to room temperature. After 18 hours, the reaction was concentrated *in vacuo* and flash column chromatography on silica using a gradient of 30-70% EtOAc/Hexanes afforded **SI5** (**SI5**, 794 mg, 1.13 mmol, 91%).

**$^1$H NMR (500 MHz, chloroform-*d*):** δ (ppm) 7.24 (m, 2H), 6.93 (m, 2H), 5.27 (m, 2H), 5.16 (m, 1H), 5.07 (m, 1H), 4.81 – 4.70 (m, 2H), 4.28 (m, 1H), 4.17 (m, 1H), 3.86 (m, 1H), 3.35 (s, 1H), 3.0 (m, 1H), 2.07 – 2.03 (m, 15H), 1.43 (s, 9H), 0.89 (s, 9H), 0.06 (m, 6H).

**$^{13}$C NMR (126 MHz, chloroform-*d*):** δ (ppm) 171.35, 170.78, 170.76, 170.44, 169.60, 169.50, 156.35, 156.03, 137.66, 127.35, 127.34, 116.80, 99.25, 77.40, 73.46, 72.93, 72.21, 71.34, 68.53, 68.49, 62.22, 62.17, 60.59, 49.32, 28.60, 25.99, 21.25, 20.90, 20.88, 20.85, 20.82, 20.79, 18.38, 14.39, -4.53, -4.85.

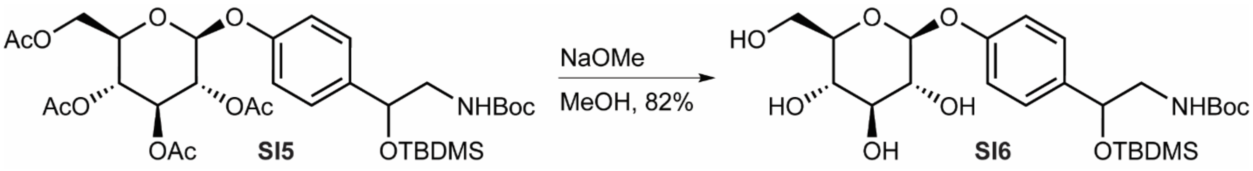

***tert*-Butyl (2-((*tert*-butyldimethylsilyl)oxy)-2-(4-(((2*S*,3*R*,4*S*,5*S*,6*R*)-3,4,5-trihydroxy-6-(hydroxymethyl)tetrahydro-2H-pyran-2-yl)oxy)phenyl)ethyl)carbamate (SI6).** To a stirred solution of **SI5** (975 mg, 1.39 mmol, 1 eq) in methanol (6.95 mL) under an inert atmosphere of argon was added sodium methoxide (7.5 mg, 0.139 mmol, 0.1 eq) in methanol (1.35 mL). The reaction was stirred for two hours, cooled to 0 °C, acidified to pH=7 with 1M HCl, and concentrated *in vacuo.* Flash column chromatography on silica using a gradient of 0-30% MeOH/DCM afforded **SI6** (**SI6**, 605 mg, 0.866 mmol, 62%).

**$^1$H NMR (500 MHz, methanol-*d$_4$*):** δ (ppm) 7.27 (m, 2H), 7.07 (m, 2H), 4.89 (m, 1H), 4.76 (m, 1H), 3.89 (m, 1H), 3.70 (m, 1H), 3.48 – 3.36 (m, 4H), 3.11 (m, 2H), 1.43 (s, 9H), 0.88 (s, 9H), 0.06 (s, 3H), -0.10 (s, 3H).

**$^{13}$C NMR (126 MHz, methanol-*d$_4$*):** δ (ppm) 158.62, 158.33, 138.11, 128.31, 128.29, 117.52, 117.42, 102.44, 80.07, 78.12, 77.98, 74.93, 74.77, 71.37, 71.36, 62.50, 50.29, 28.83, 28.72, 26.37, 19.10, -4.58, -4.70.

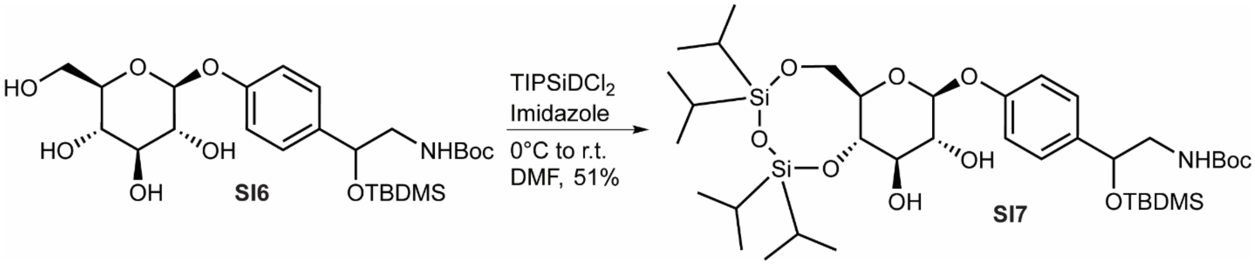

***tert*-Butyl (2-((*tert*-butyldimethylsilyl)oxy)-2-(4-(((6a*R*,8*S*,9*R*,10*R*,10a*S*)-9,10-dihydroxy-2,2,4,4-tetraisopropylhexahydropyrano[3,2-f][1,3,5,2,4]trioxadisilocin-8-yl)oxy)phenyl)-ethyl)carbamate (SI7).** To a stirred solution of **SI6** (605 mg, 1.14 mmol, 1 eq) in DMF (2.28 mL) at 0 °C under an inert atmosphere of argon was added imidazole (341 mg, 5.01 mmol, 4.4 eq) then $TIPSiDCl_2$ (445 mg, 1.37 mmol, 1.2 eq) dropwise. The reaction was left to warm to room temperature over three hours. The reaction was then diluted in $H_2O$ and extracted with DCM three times. The combined organic phase was washed with brine, dried over magnesium sulfate, and concentrated *in vacuo.* Flash column chromatography on silica using a gradient of 0-40% EtOAc/Hexanes afforded **SI7** (**SI7**, 405 mg, 0.52 mmol, 46%).

**$^1$H NMR (500 MHz, methanol-*d$_4$*):** δ (ppm) 7.22 (m, 2H), 7.06 (m, 2H), 4.92 (d, 1H), 4.76 (m, 1H), 4.18 (m, 1H), 3.94 (m, 1H), 3.82 (t, 1H), 3.53 (t, 1H), 3.42 (m, 2H), 3.11 (m, 2H), 1.43 (s, 9H), 1.18 – 1.06 (m, 25H), 1.02 (m, 3H), 0.88 (s, 9H), 0.06 (s, 3H), -0.09 (s, 3H).

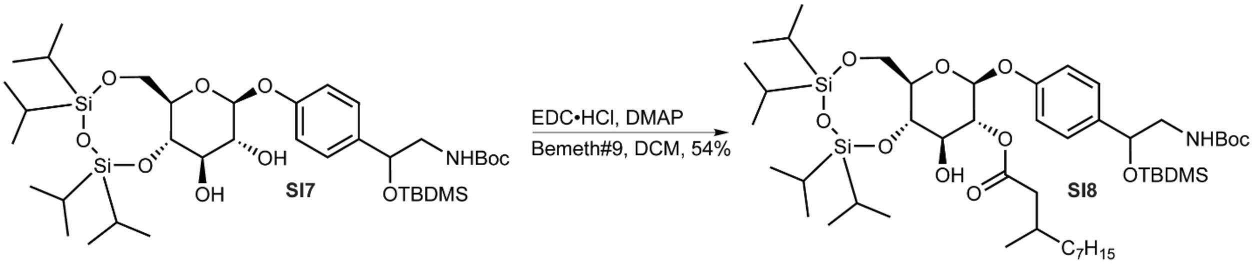

**(6a*R*,8*S*,9*R*,10*R*,10a*S*)-8-(4-(2,2,3,3,10,10-hexamethyl-8-oxo-4,9-dioxa-7-aza-3-silaundecan-5-yl)phenoxy)-10-hydroxy-2,2,4,4-tetraisopropylhexahydropyrano[3,2-f]-[1,3,5,2,4]trioxadisilocin-9-yl 3-methyldecanoate (SI8).** To a stirred solution of bemeth#9 (**2**) (44 mg, 0.236 mmol, 1 eq) in DCM (2.36 mL) was added EDC (90 mg, 0.472 mmol, 2 eq) and the reaction was stirred for ten minutes at room temperature. DMAP (72 mg, 0.59 mmol, 2.5 eq) was added and the reaction was stirred for another ten minutes at room temperature. **SI7** (218 mg, 0.283 mmol, 1.2 eq) was added and the reaction was stirred for 18 hours. After 18 hours the reaction mixture was concentrated *in vacuo.* and flash column chromatography on silica using a gradient of 0-20% EtOAc/Hexanes afforded **SI8** (**SI8**, 120 mg, 0.127 mmol, 45%).

**$^1$H NMR (500 MHz, chloroform-*d*):** δ (ppm) 7.26 (m, 2H), 6.96 (m, 2H), 5.13 (m, 1H), 4.95 (m, 1H), 4.76 (m, 1H), 4.19 (m, 1H), 3.97 (m, 1H), 3.89 (t, 1H), 3.71 (t, 1H), 3.49 (d, 1H), 3.10 (m, 2H), 2.35 (m, 1H), 2.18 (m, 1H), 1.92 (m, 1H), 1.43 (s, 9H), 1.38 – 0.97 (m, 42H), 0.93 – 0.85 (m, 15H), 0.06 (s, 3H), -0.10 (s, 3H).

**$^{13}$C NMR (126 MHz, chloroform-*d*):** δ (ppm) 173.88, 158.32, 157.91, 138.38, 128.40, 117.23, 100.36, 80.06, 77.88, 75.48, 75.24, 74.79, 74.72, 70.67, 62.07, 50.28, 42.86, 37.68, 37.60, 33.07, 31.60, 30.80, 30.45, 28.84, 28.01, 26.39, 23.76, 20.09, 20.05, 19.12, 18.00, 17.85, 17.82, 17.80, 17.75, 17.72, 17.65, 17.53, 14.82, 14.51, 14.48, 13.91, 13.76, -4.54, -4.69.

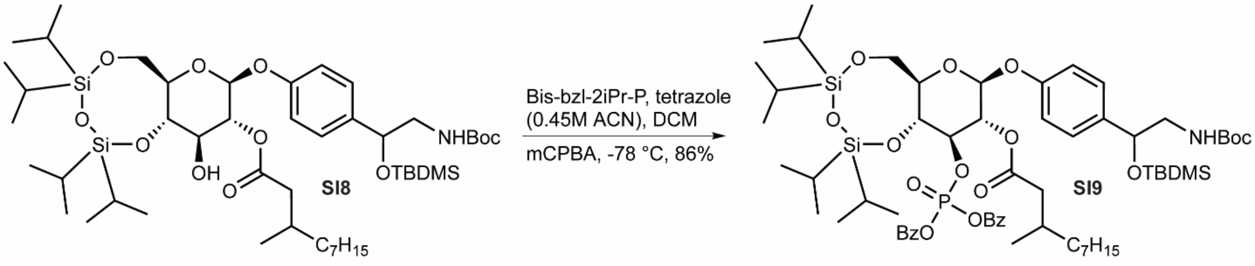

**(6a*R*,8*S*,9*R*,10*R*,10a*R*)-10-((bis(benzyloxy)phosphoryl)oxy)-8-(4-(2,2,3,3,10,10-hexamethyl-8-oxo-4,9-dioxa-7-aza-3-silaundecan-5-yl)phenoxy)-2,2,4,4-tetraisopropylhexahydro-pyrano[3,2-f][1,3,5,2,4]trioxadisilocin-9-yl 3-methyldecanoate (SI9).** To a stirred solution of

**SI8** (115 mg, 0.122 mmol, 1 eq) in DCM (1 mL) was added dibenzyl *N,N*-diisopropylphosphoramidite (143 µL, 0.427 mmol, 3.5 eq) and tetrazole as a 0.45 M solution in acetonitrile (948 µL, 0.427 mmol, 3.5 eq). When the starting material was consumed by TLC, the reaction was flushed with argon, cooled to -78 °C, and mCPBA (55 mg, 0.317 mmol, 2.6 eq) was added dropwise as a solution in DCM (450 µL). The reaction was left at -78 °C for 20 minutes and then taken out of the -78 °C bath for ten minutes. After ten minutes the reaction was returned to -78 °C and a second portion of mCPBA was added (21 mg, 0.122 mmol, 1 eq) in DCM (200 µL). The reaction was then taken out of the -78 °C bath for ten minutes. After ten minutes the reaction was returned to -78 °C and a third portion of mCPBA was added (21 mg, 0.122 mmol, 1 eq) in DCM (200 µL). The reaction was then taken out of the -78 °C bath for ten minutes. After ten minutes the reaction was complete by TLC and cooled to -78 °C. The reaction was then diluted in DCM, warmed to room temperature, and water was added. The mixture was extracted into DCM three times and then the combined organic phase was washed with sat. aq. sodium sulfate two times, washed with sat. aq. sodium bicarbonate, washed with brine, and dried over magnesium sulfate. The crude extract was concentrated *in vacuo.* and flash column chromatography on silica using a gradient of 0-50% EtOAc/Hexanes afforded **SI9** (**SI9**, 124 mg, 0.103 mmol, 84.6%).

**$^{1}$H NMR (500 MHz, chloroform-*d*):** δ (ppm) 7.36 – 7.27 (m, 10H), 7.19 (m, 2H), 6.87 (m, 2H), 5.28 (m, 1H), 5.03 – 4.90 (m, 5H), 4.76 (m, 1H), 4.71 (m, 1H), 4.51 (m, 1H), 4.18 – 4.09 (m, 3H), 4.04 (m, 1H), 3.33 (m, 2H), 3.0 (m, 1H), 2.24 (m, 1H), 2.02 (m, 1H), 1.83 (m, 1H), 1.43 (s, 9H), 1.32 – 0.79 (m, 63H), 0.03 (s, 3H), -0.12 (d, 3H).

**$^{13}$C NMR (126 MHz, chloroform-*d*):** δ (ppm) 172.56, 172.36, 171.35, 156.62, 156.04, 135.96, 135.91, 128.69, 128.66, 128.65, 128.31, 128.30, 128.26, 127.19, 127.16, 116.39, 116.32, 99.09, 99.00, 80.66, 79.38, 77.40, 76.68, 73.52, 71.64, 69.70, 69.65, 69.60, 68.75, 60.86, 60.59, 53.61, 49.29, 42.02, 41.89, 36.79, 36.77, 32.07, 31.78, 30.39, 30.21, 29.89, 29.50, 28.61, 27.02, 26.99, 26.00, 22.87, 22.84, 21.25, 19.67, 19.59, 19.57, 18.38, 17.55, 17.49, 17.45, 17.39, 17.34, 17.17, 14.39, 14.31, 13.40, 13.22, 13.05, 12.95, -4.50, -4.52, -4.87.

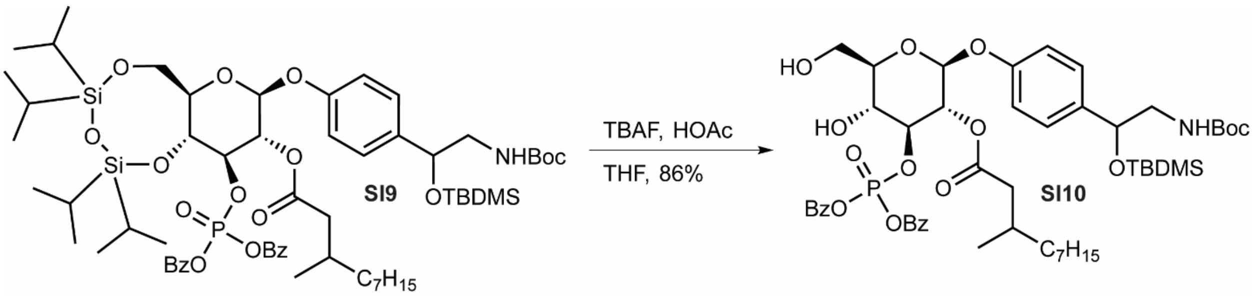

**(2*S*,3*R*,4*S*,5*R*,6*R*)-2-(4-(2-amino-1-((*tert*-butyldimethylsilyl)oxy)ethyl)phenoxy)-4-((bis(benzyloxy)phosphoryl)oxy)-5-hydroxy-6-(hydroxymethyl)tetrahydro-2H-pyran-3-yl 3-methyldecanoate (SI10).** To a stirred solution of **SI9** (115 mg, 0.097 mmol, 1 eq.) in THF (970 µL) at 0 °C under an inert atmosphere of argon was added glacial acetic acid (16.5 µL, 0.291 mmol, 3 eq.) dropwise followed by TBAF 1M in THF (291 µL, 0.291 mmol, 3 eq.) dropwise. The reaction was left to warm to room temperature over 70 minutes. Upon completion, the reaction was diluted in DCM, concentrated *in vacuo.* to a paste, and flash column chromatography on silica using a gradient of 50-80% EtOAc/Hexanes afforded **SI10** (**SI10**, 69 mg, 0.0817 mmol, 84.3%).

**$^1$H NMR (500 MHz, methanol-*d$_4$*):** δ (ppm) 7.40 – 7.31 (m, 10H), 7.27 (m, 2H), 6.97 (m, 2H), 5.24 – 5.17 (m, 2H), 5.13 (dd, 2H), 5.07 – 4.96 (m, 2H), 4.76 (m, 1H), 4.63 – 4.55 (m, 1H), 3.94 (dd, 1H), 3.81 – 3.73 (m, 2H), 3.62 – 3.56 (m, 1H), 3.14 (m, 1H), 3.08 – 2.94 (m, 2H), 2.20 – 2.14 (m, 1H), 2.09 (m, 1H), 2.01 (m, 1H), 1.92 – 1.86 (m, 1H), 1.78 (q, 1H), 1.63 (m, 1H), 1.42 (s, 9H), 1.39 (m, 1H), 1.35 – 1.08 (m, 15H), 1.01 (m, 3H), 0.90 (m, 3H), 0.82 (dd, 1H), 0.77 (d, 2H), 0.05 (s, 3H), -0.09 (s, 3H).

**$^{13}$C NMR (126 MHz, methanol-*d$_4$*):** δ (ppm) 173.65, 173.58, 172.98, 158.31, 157.84, 138.57, 137.34, 137.28, 137.12, 137.06, 129.71, 129.66, 129.64, 129.17, 129.15, 129.13, 128.41, 117.17, 82.92, 80.06, 77.79, 74.76, 74.69, 73.04, 70.97, 70.92, 70.05, 61.94, 61.53, 54.80, 54.21, 50.29, 42.77, 42.59, 37.61, 37.51, 33.04, 32.76, 31.34, 31.29, 30.78, 30.76, 30.67, 30.45, 30.42, 28.84, 27.97, 27.94, 27.88, 27.41, 26.39, 23.76, 23.70, 21.09, 20.86, 20.04, 19.99, 19.11, 14.49, 14.46, 14.43, 14.00, -4.56, -4.68.

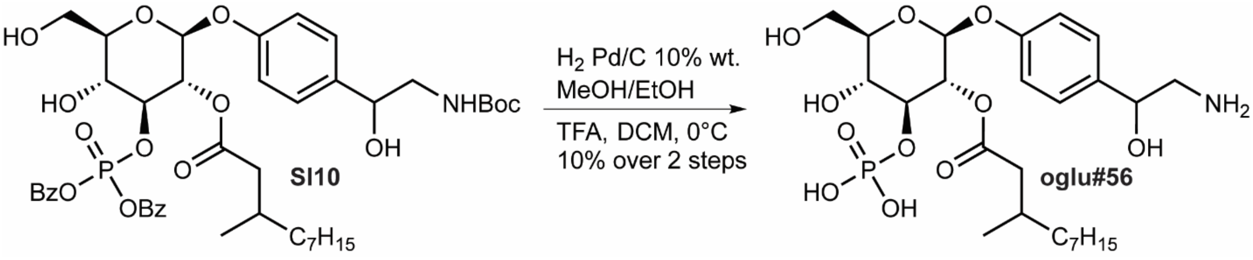

**(2*S*,3*R*,4*S*,5*R*,6*R*)-2-(4-(2-amino-1-hydroxyethyl)phenoxy)-5-hydroxy-6-(hydroxymethyl)-4-(phosphonooxy)tetrahydro-2H-pyran-3-yl 3-methyldecanoate (oglu#56).** To a stirred solution of **SI10** (69 mg, 0.0817 mmol, 1 eq.) in a mixture of methanol and ethanol (1:1, 1 mL) was added palladium on carbon 10 wt.% loading (60 mg). The reaction was maintained at 0 °C and flushed with argon for five minutes followed by hydrogen gas for ten minutes. Upon completion the reaction was flushed for argon for ten minutes at 0 °C, warmed to room temperature, filtered through a pad of Celite, and concentrated *in vacuo.* The crude filtrate was then diluted in DCM (830 μL) and stirred at 0 °C. A solution of TFA (55 μL) in 200 μL of DCM was added dropwise and the reaction was maintained at 0 °C for two hours. Upon completion, the reaction was diluted in DCM, concentrated *in vacuo.* to a paste, diluted again in DCM, and flash column chromatography on silica using a gradient of 20-100% MeOH/DCM afforded **oglu#56** (**oglu#56**, 5 mg, 0.008 mmol, 11% over two steps).

**$^1$H NMR (600 MHz, methanol-*d$_4$*):** δ (ppm) 7.33 (m, 2H), 7.00 (m, 2H), 5.11 (m, 1H), 5.04 (m, 1H), 4.82 – 4.77 (m, 1H), 4.25 (m, 1H), 3.87 (dt, 1H), 3.68 (m, 1H), 3.60 (m, 1H), 3.49 (m, 1H), 3.05 (dt, 1H), 2.96 – 2.88 (m, 1H), 2.44 – 2.35 (m, 1H), 2.16 (m, 1H), 1.87 (m, 1H), 1.35 – 1.05 (m, 15H), 0.87 (m, 6H).

**HRMS (ESI -) *m/z*:** [M]$^-$ calcd. for $C_{25}H_{41}NO_{11}P^-$ 562.24227, found 562.24420.

For additional characterization see section **4.1: 2D characterization of oglu#56**.

### 2. NMR spectra appendix

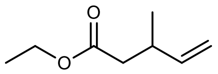

**SI1**

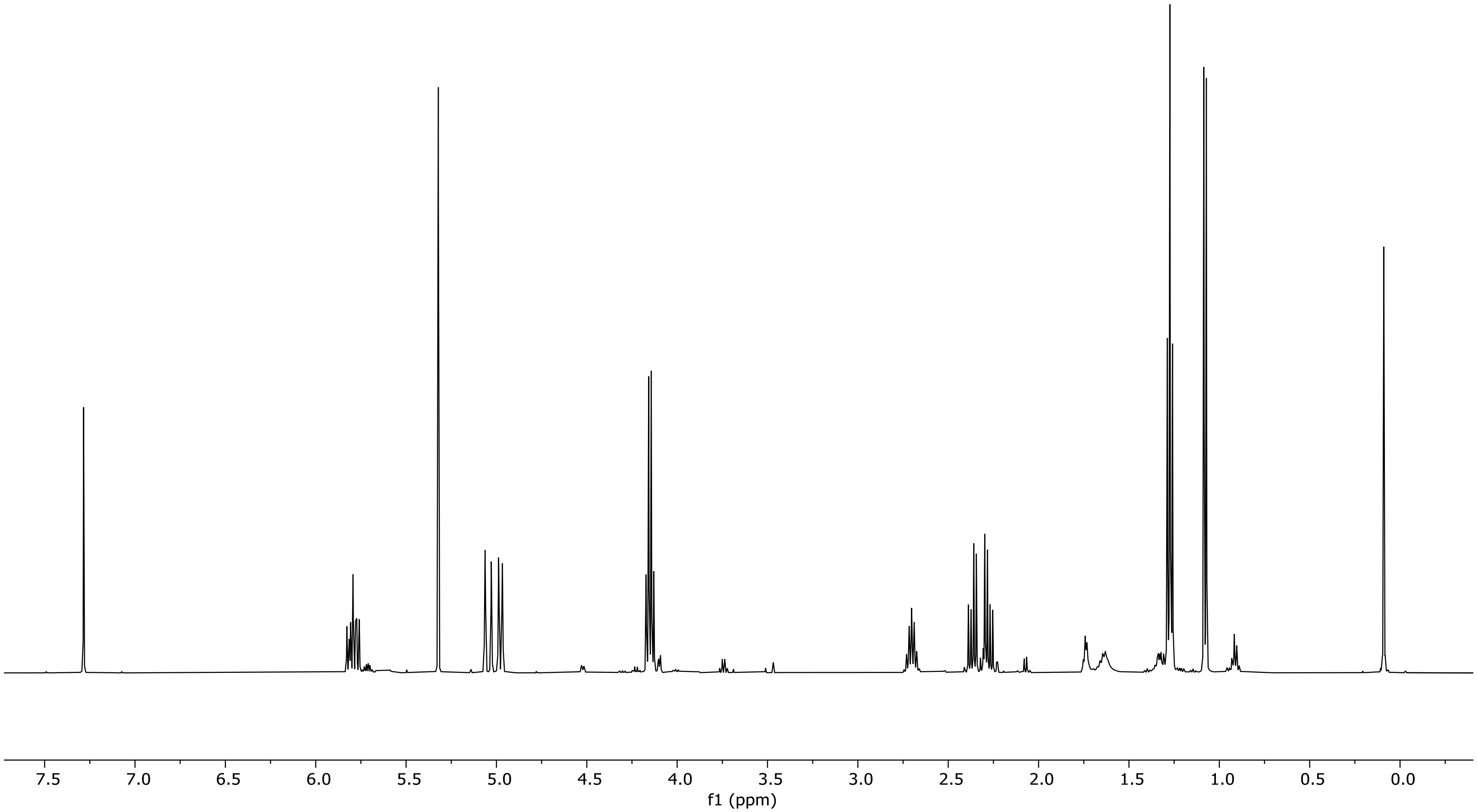

[1]H NMR of ethyl 3-methylpent-4-enoate (**SI1**) in chloroform-*d* (500 MHz).

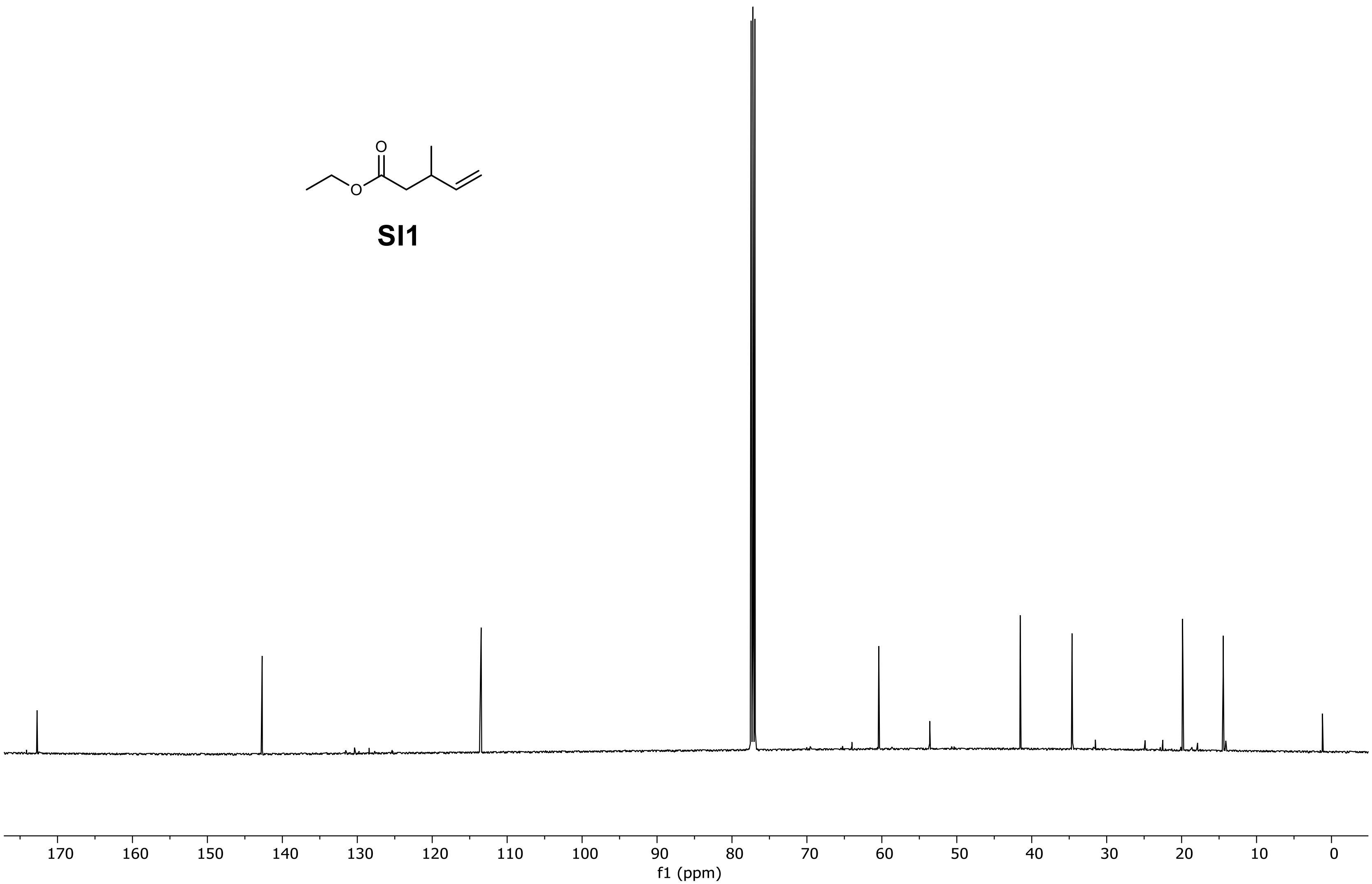

$^{13}$C NMR of ethyl 3-methylpent-4-enoate (**SI1**) in chloroform-*d* (126 MHz).

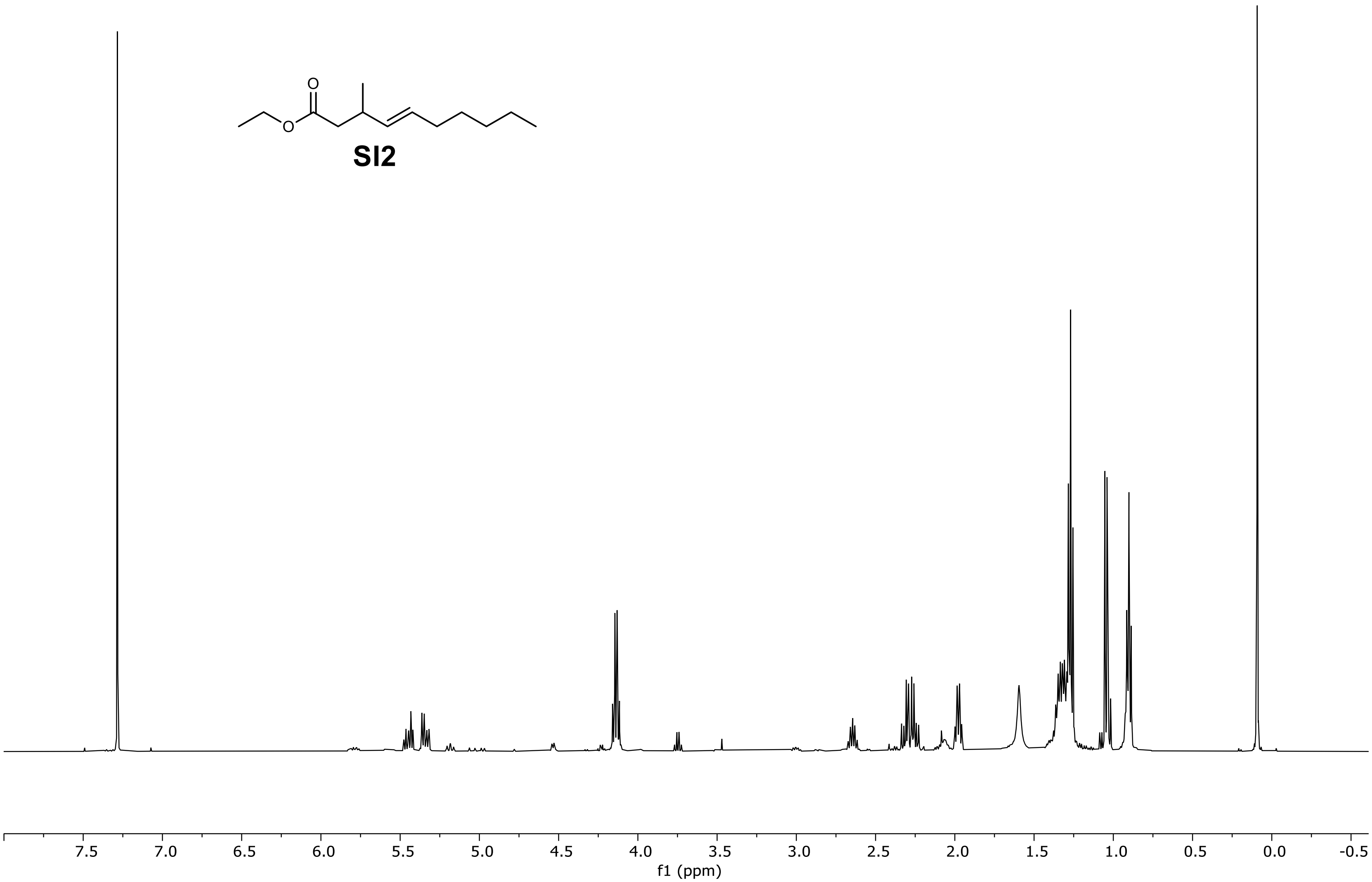

$^{1}$H NMR of ethyl (*E*)-3-methyldec-4-enoate (**SI2**) in chloroform-*d* (500 MHz).

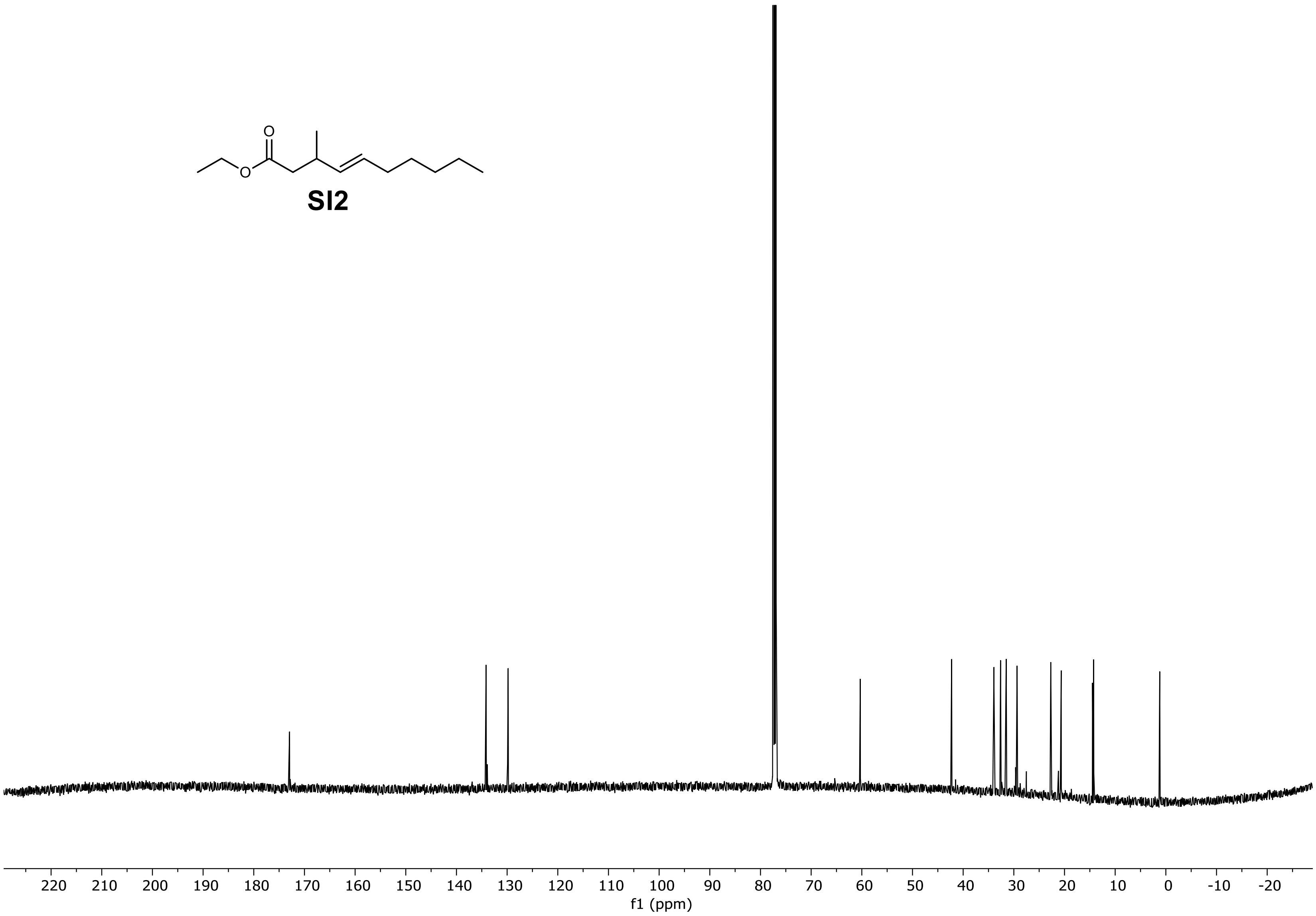

$^{13}$C NMR of ethyl (*E*)-3-methyldec-4-enoate (**SI2**) in chloroform-*d* (126 MHz).

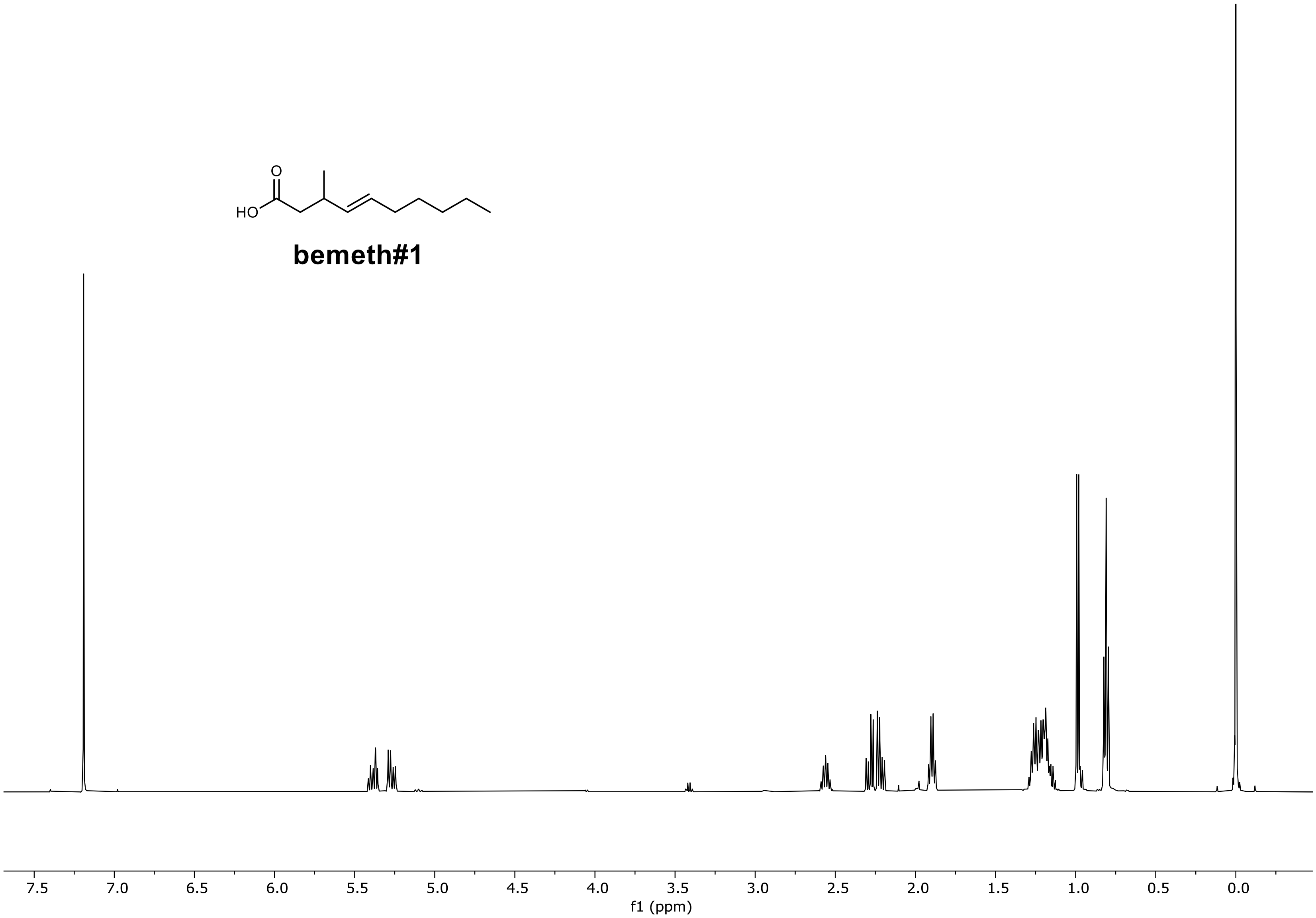

[1]H NMR of **bemeth#1** in chloroform-*d* (500 MHz).

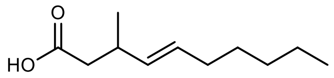

**bemeth#1**

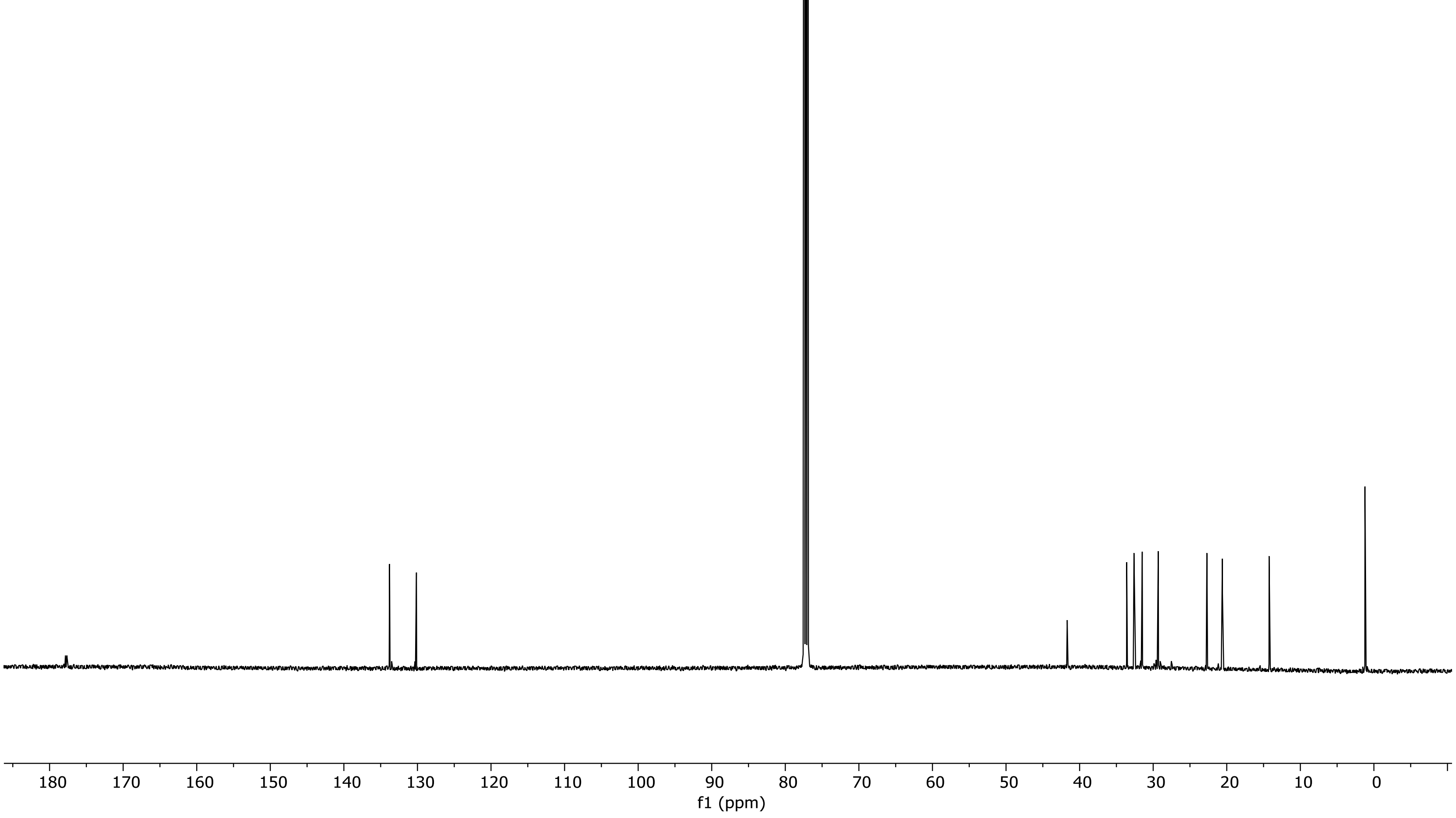

$^{13}C$ NMR of **bemeth#1** in chloroform-*d* (126 MHz).

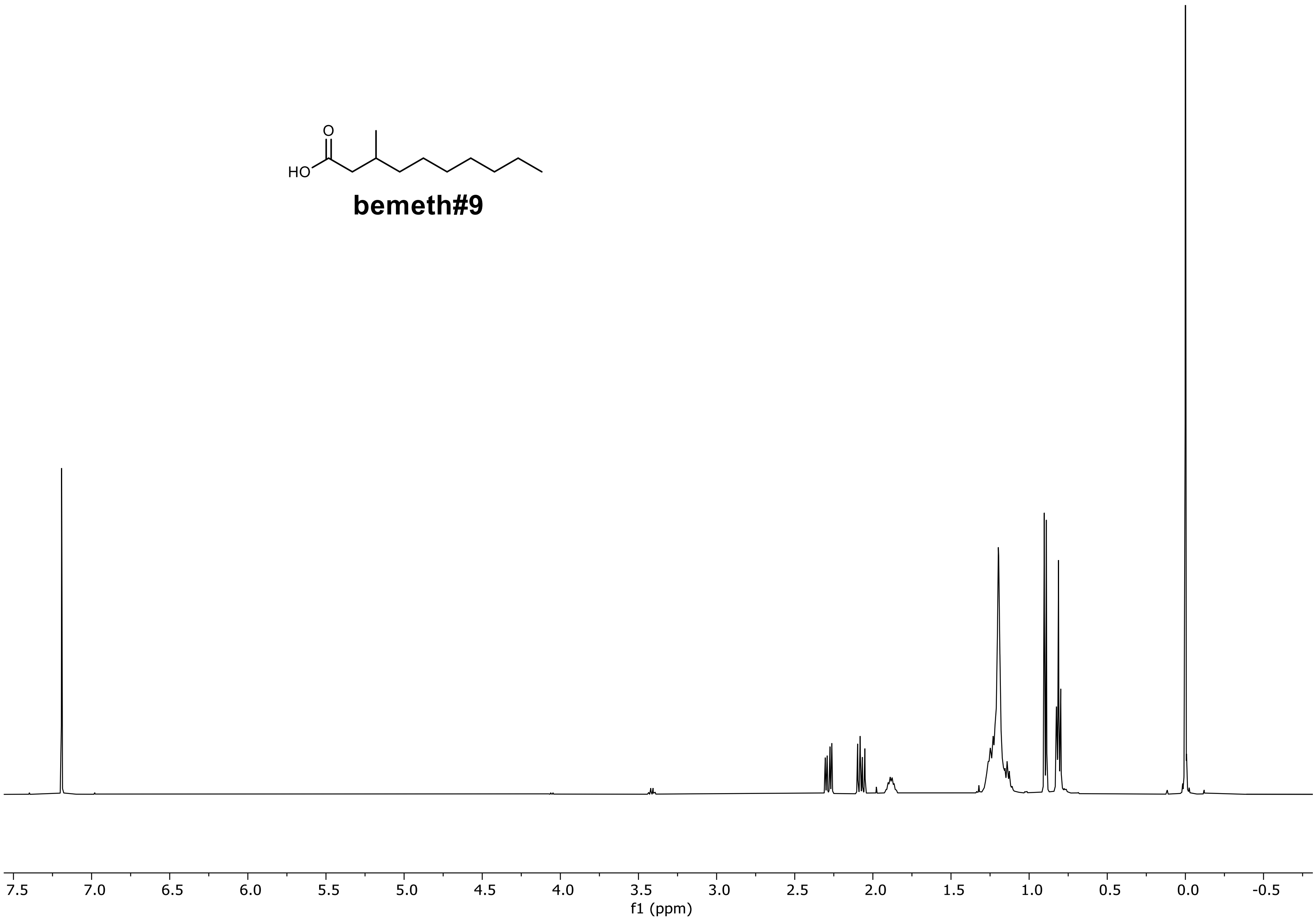

$^{1}$H NMR of **bemeth#9** in chloroform-*d* (500 MHz).

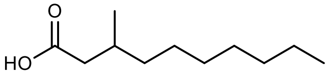

**bemeth#9**

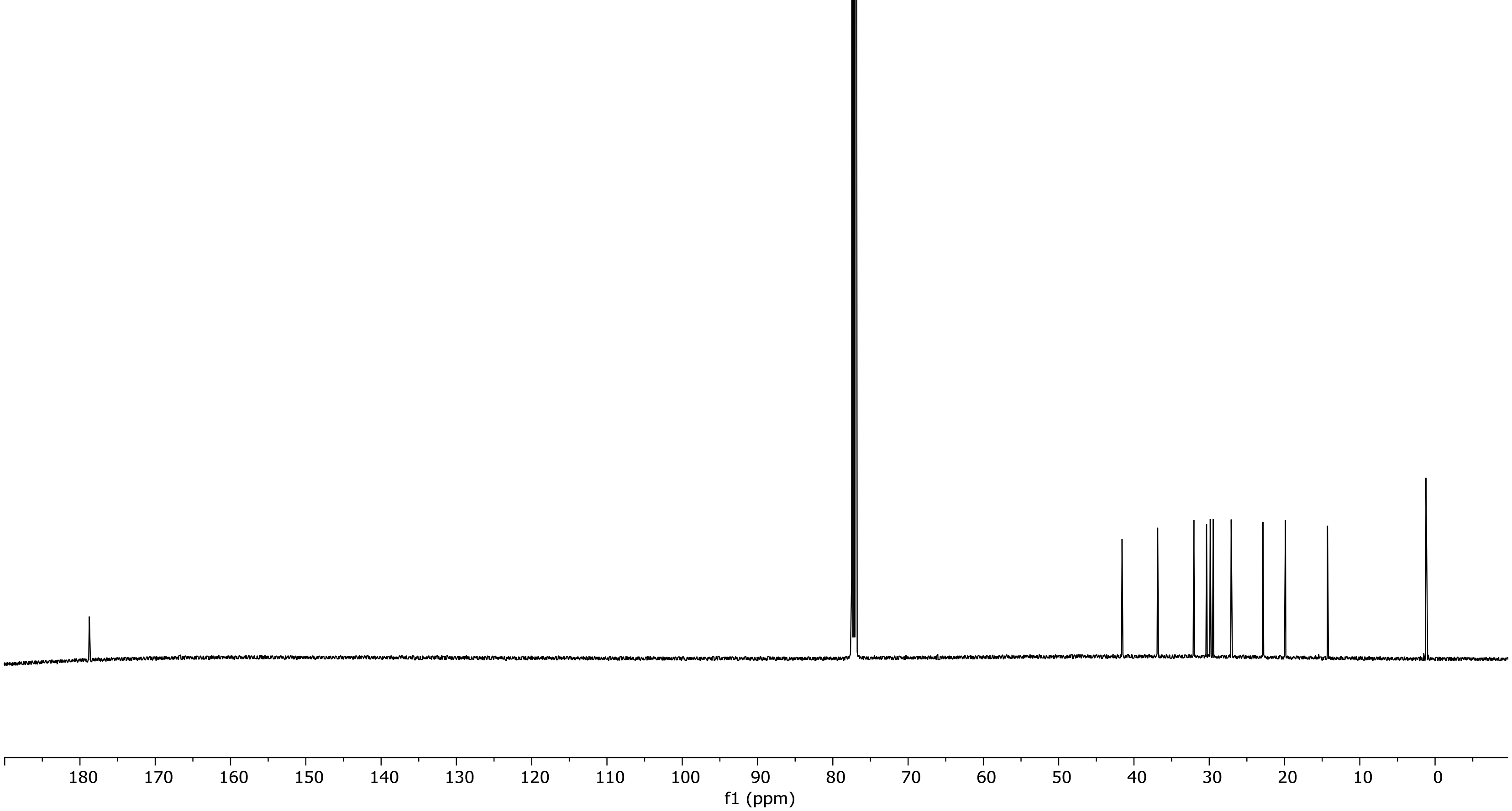

$^{13}C$ NMR of **bemeth#9** in chloroform-*d* (126 MHz).

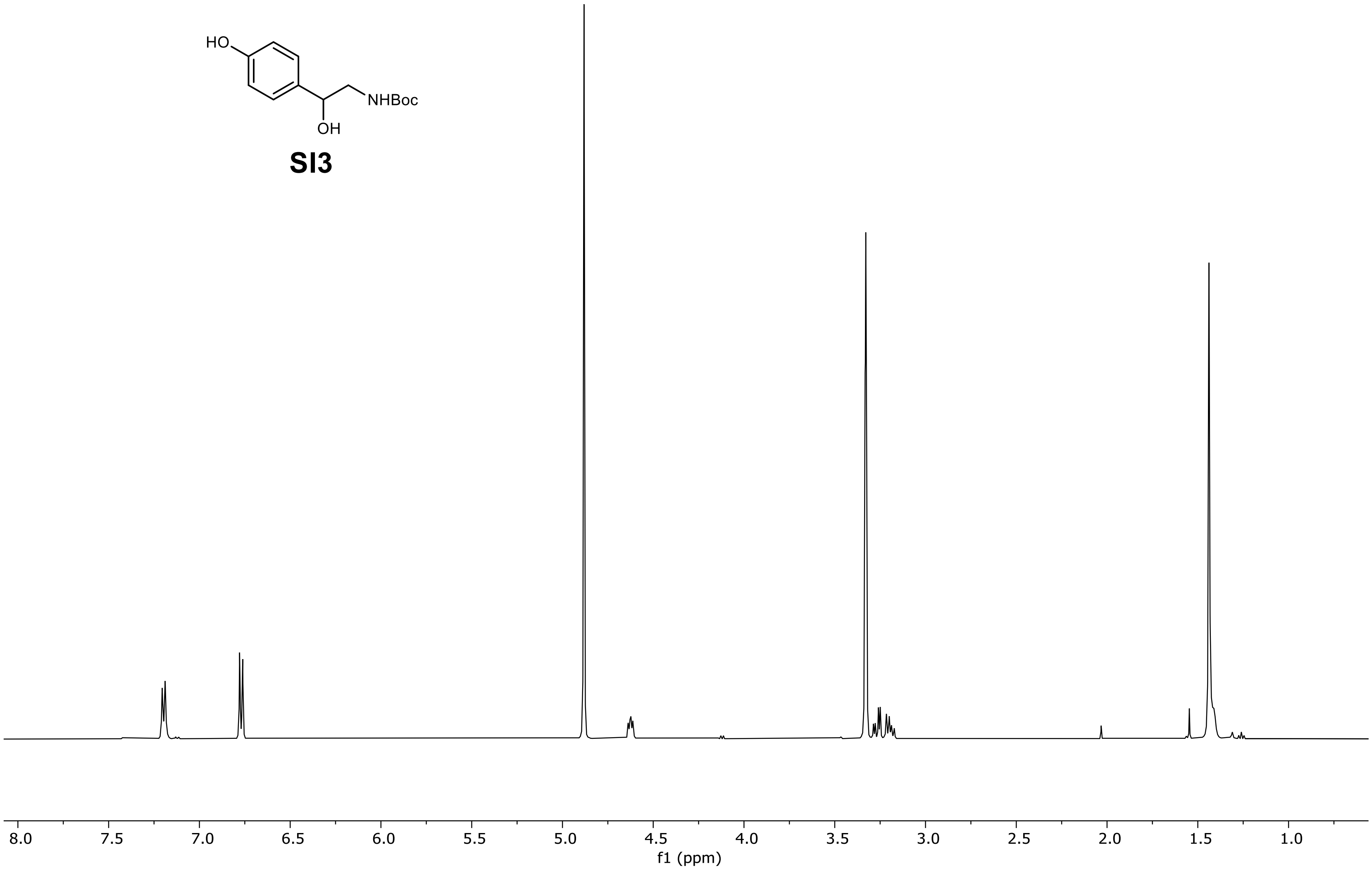

$^1$H NMR of *tert*-butyl (2-hydroxy-2-(4-hydroxyphenyl)ethyl)carbamate (**SI3**) in methanol-$d_4$ (500 MHz).

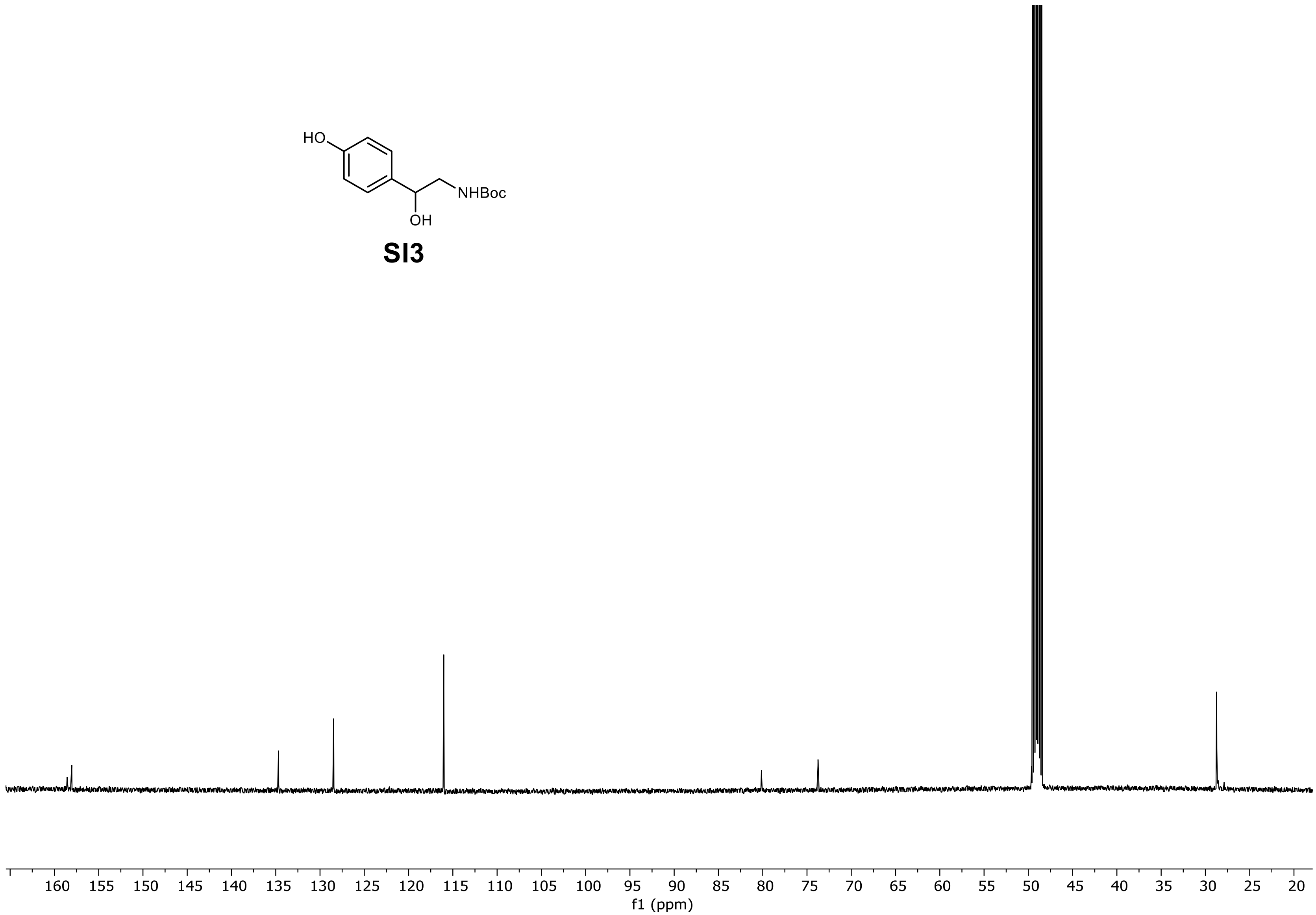

$^{13}$C NMR of *tert*-butyl (2-hydroxy-2-(4-hydroxyphenyl)ethyl)carbamate (**SI3**) in methanol-$d_4$ (126 MHz).

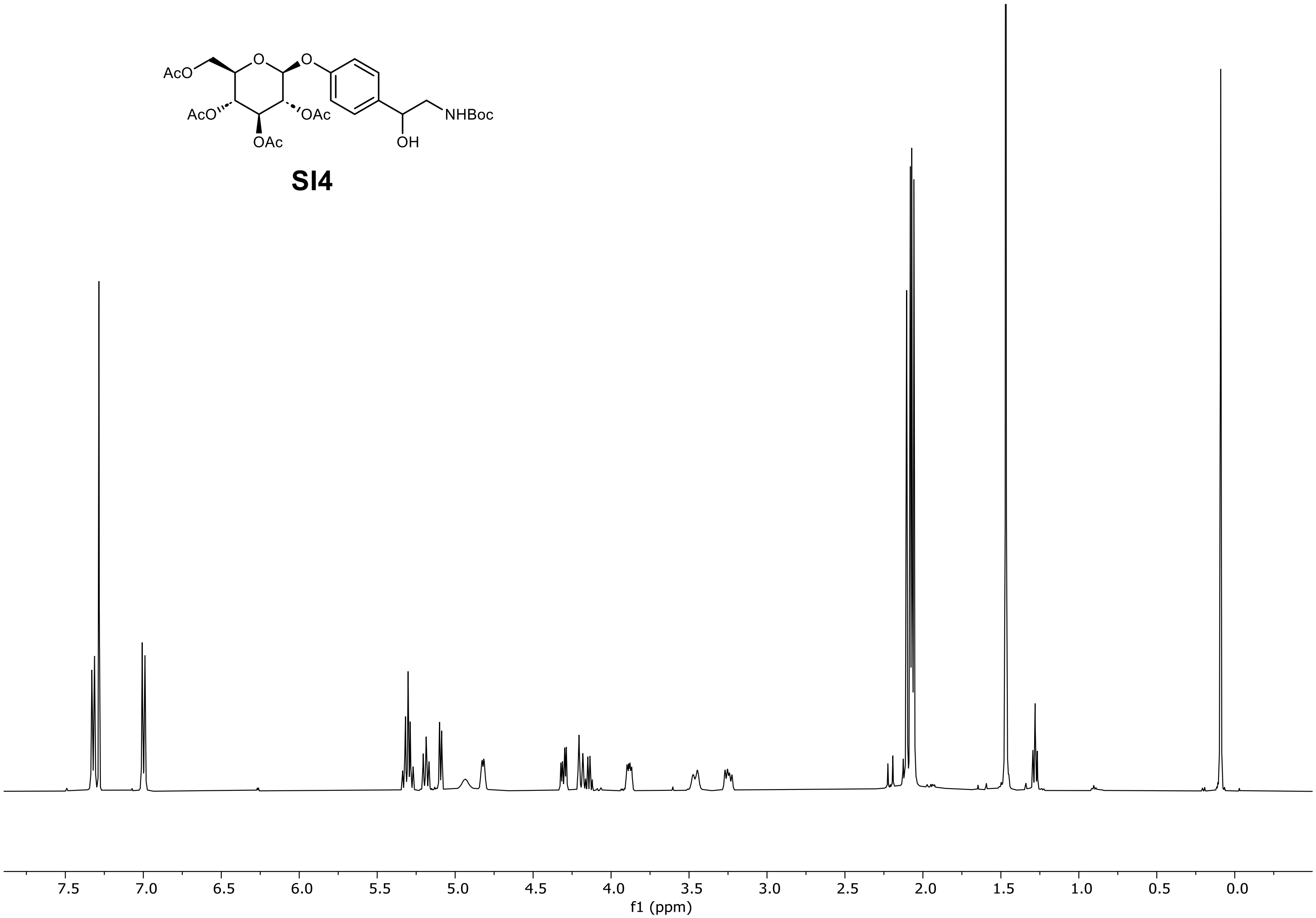

[1]H NMR of (2*R*,3*R*,4*S*,5*R*,6*S*)-2-(acetoxymethyl)-6-(4-(2-((*tert*-butoxycarbonyl)amino)-1-hydroxyethyl)phenoxy)tetrahydro-2H-pyran-3,4,5-triyl triacetate (**SI4**) in chloroform-*d* (500 MHz).

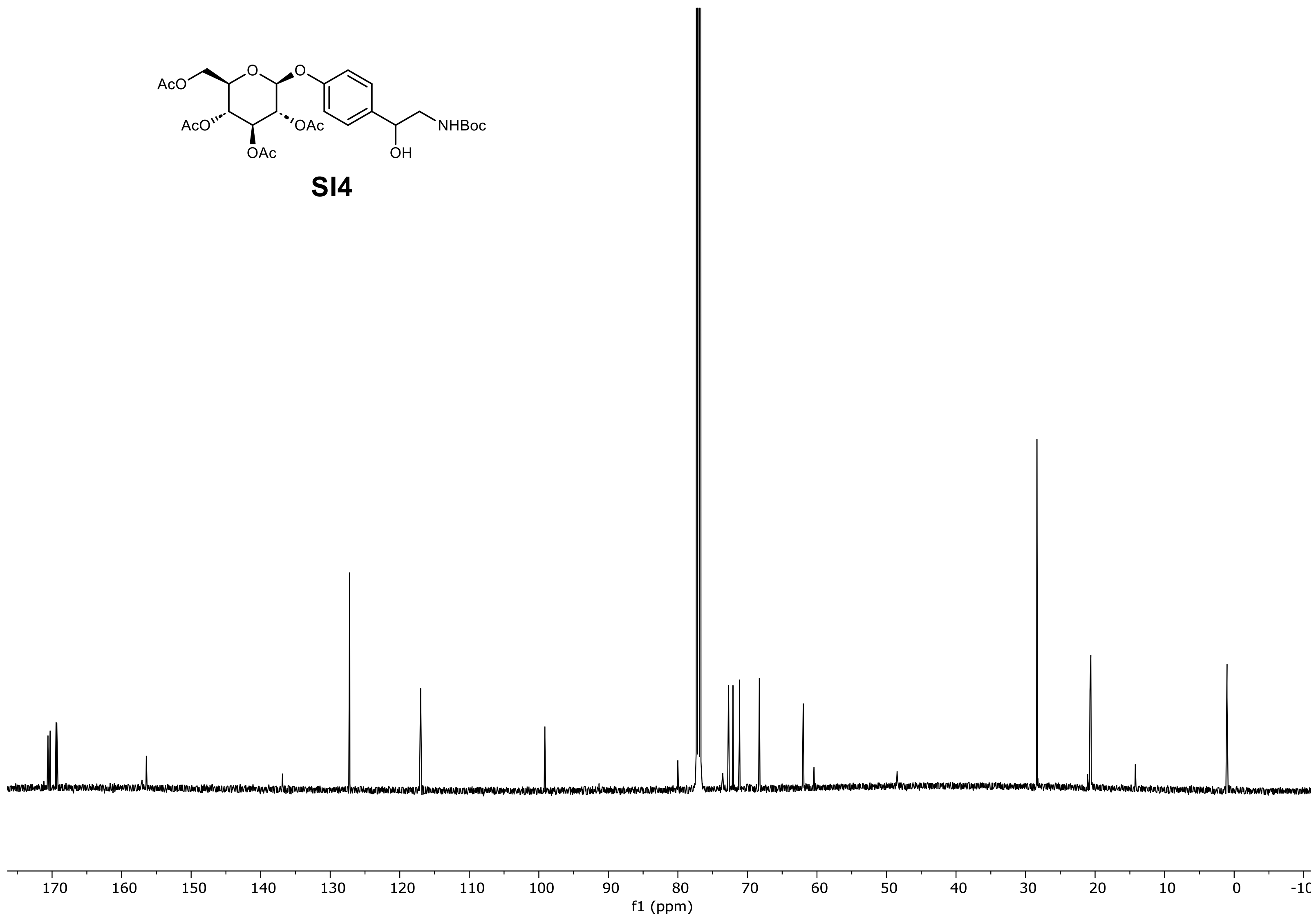

$^{13}C$ NMR of (2*R*,3*R*,4*S*,5*R*,6*S*)-2-(acetoxymethyl)-6-(4-(2-((*tert*-butoxycarbonyl)amino)-1-hydroxyethyl)phenoxy)tetrahydro-2H-pyran-3,4,5-triyl triacetate (**SI4**) in chloroform-*d* (126 MHz).

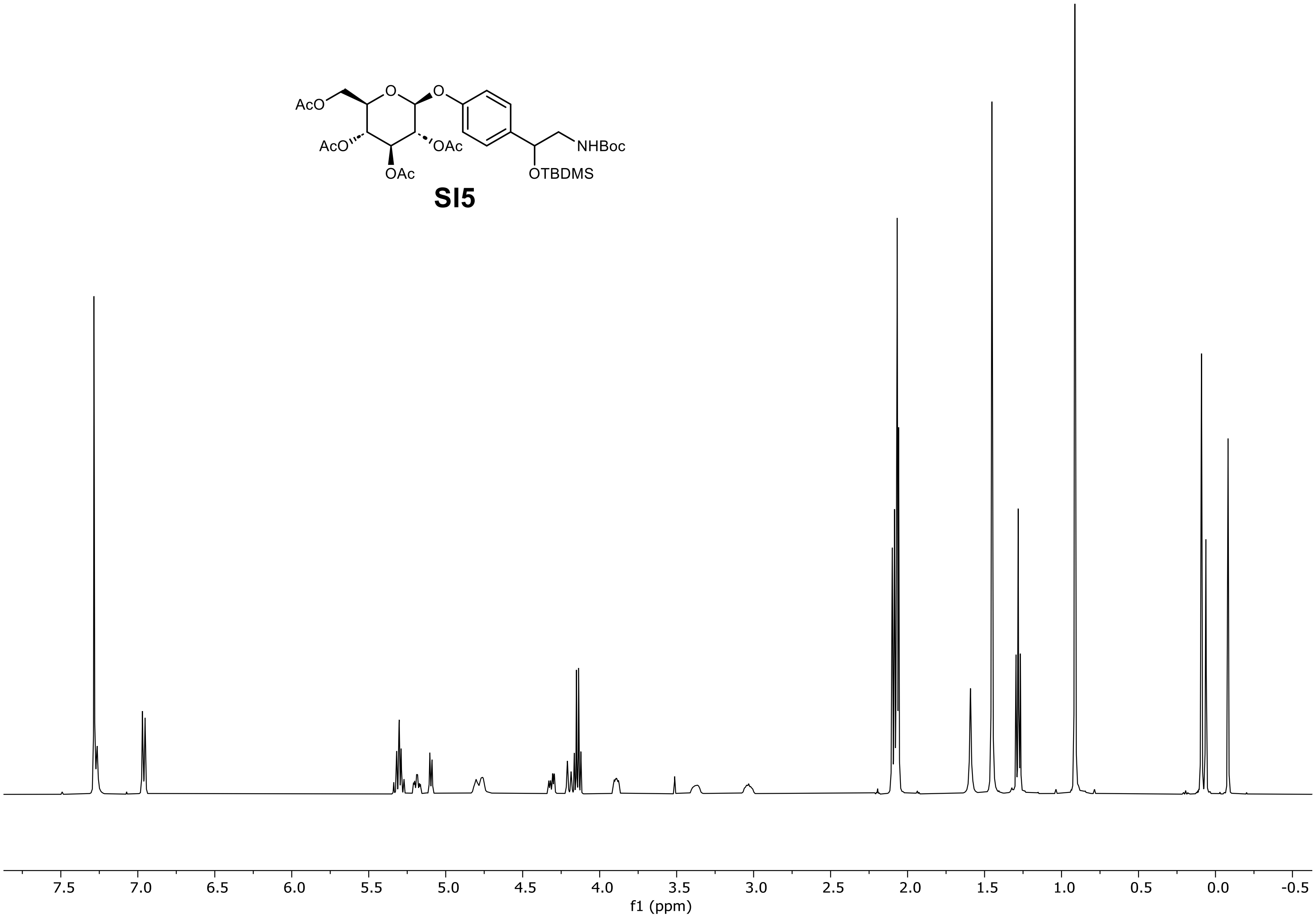

[1]H NMR of (2*R*,3*R*,4*S*,5*R*,6*S*)-2-(acetoxymethyl)-6-(4-(2,2,3,3,10,10-hexamethyl-8-oxo-4,9-dioxa-7-aza-3-silaundecan-5-yl)phenoxy)-tetrahydro-2H-pyran-3,4,5-triyl triacetate (**SI5**) in chloroform-*d* (500 MHz).

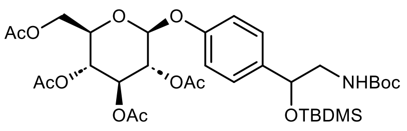

**SI5**

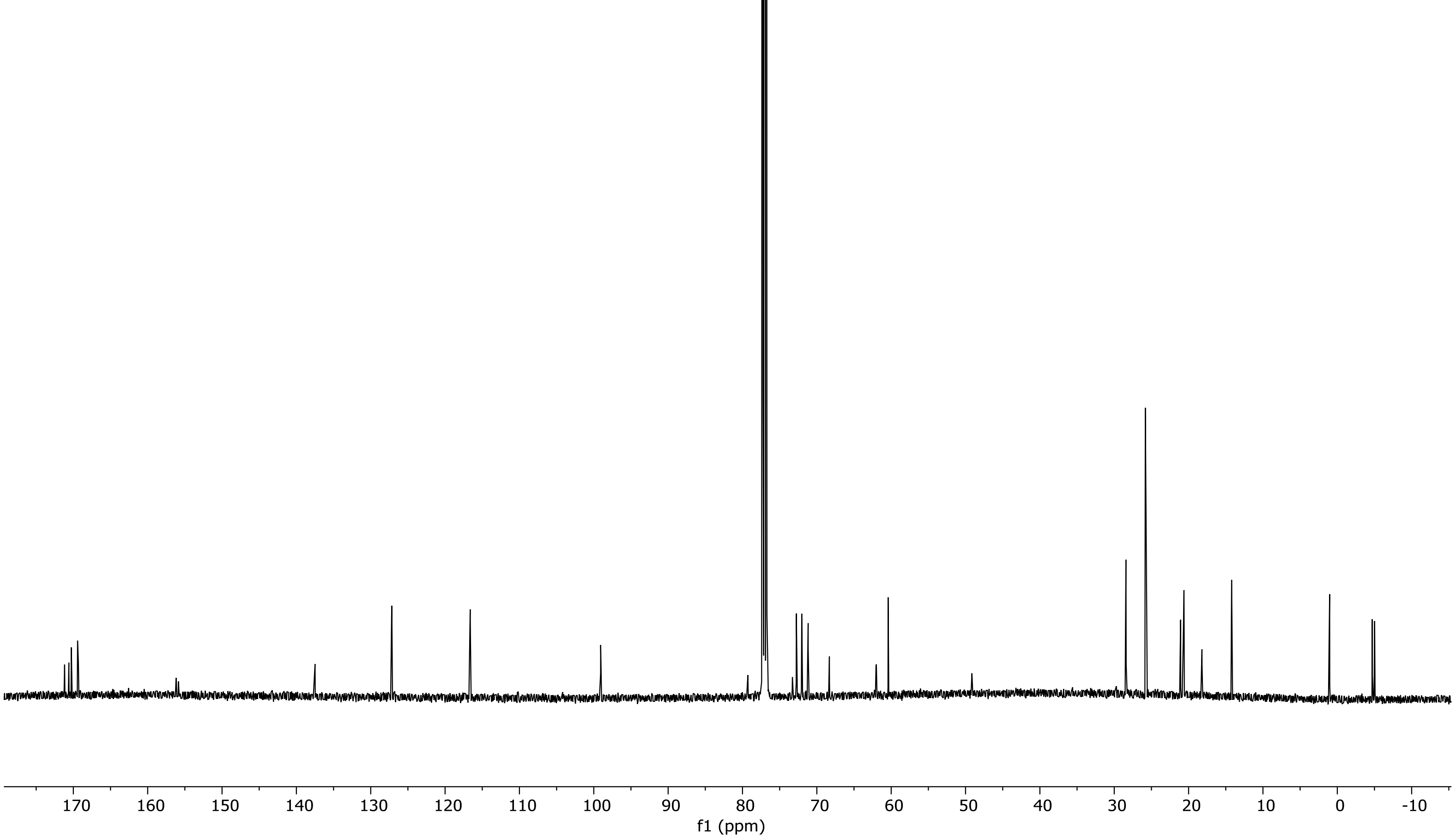

$^{13}C$ NMR of (2*R*,3*R*,4*S*,5*R*,6*S*)-2-(acetoxymethyl)-6-(4-(2,2,3,3,10,10-hexamethyl-8-oxo-4,9-dioxa-7-aza-3-silaundecan-5-yl)-phenoxy)tetrahydro-2H-pyran-3,4,5-triyl triacetate (**SI5**) in chloroform-*d* (126 MHz).

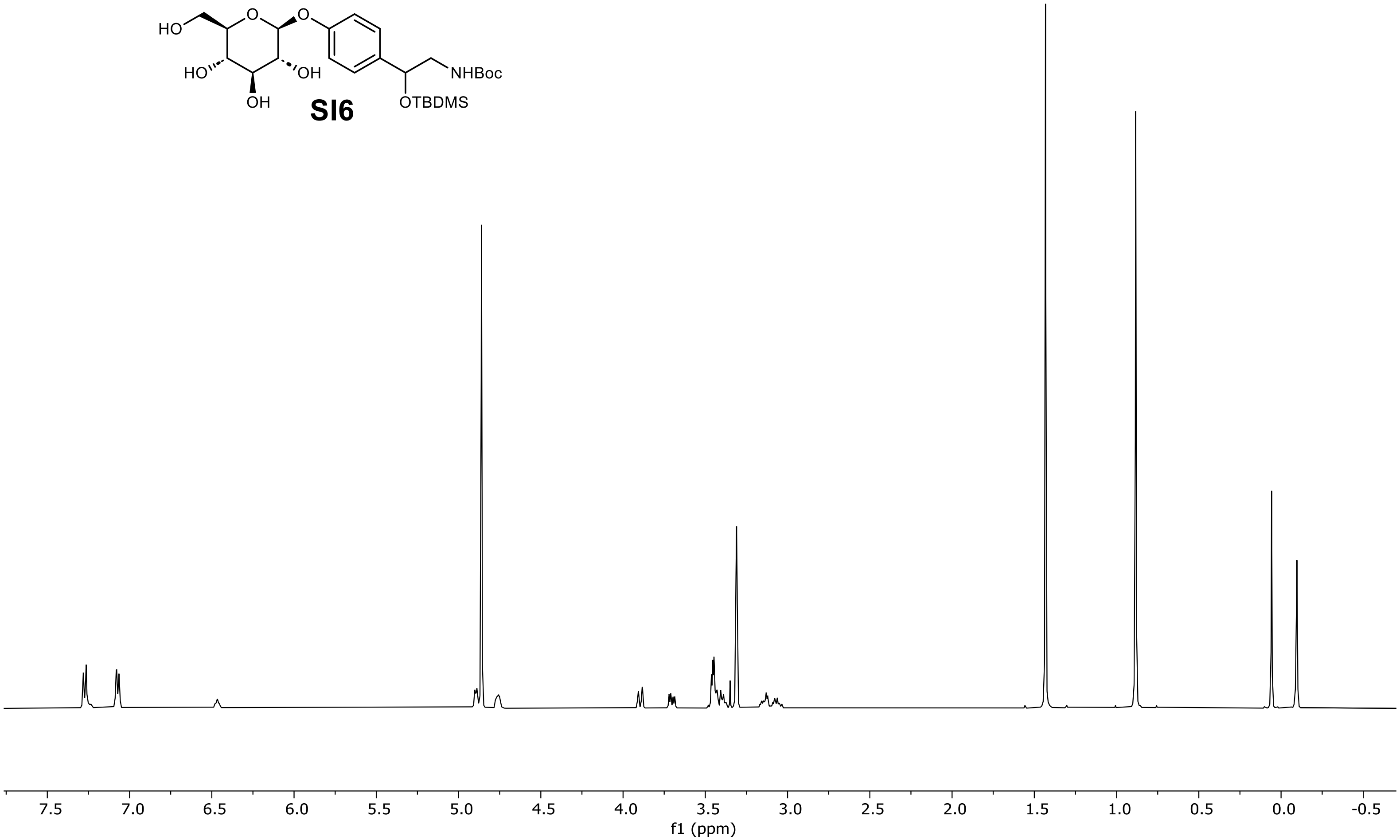

[1]H NMR of *tert*-butyl (2-((*tert*-butyldimethylsilyl)oxy)-2-(4-(((2*S*,3*R*,4*S*,5*S*,6*R*)-3,4,5-trihydroxy-6-(hydroxymethyl)tetrahydro-2H-pyran-2-yl)oxy)phenyl)ethyl)carbamate (**SI6**) in methanol-$d_4$ (500 MHz).

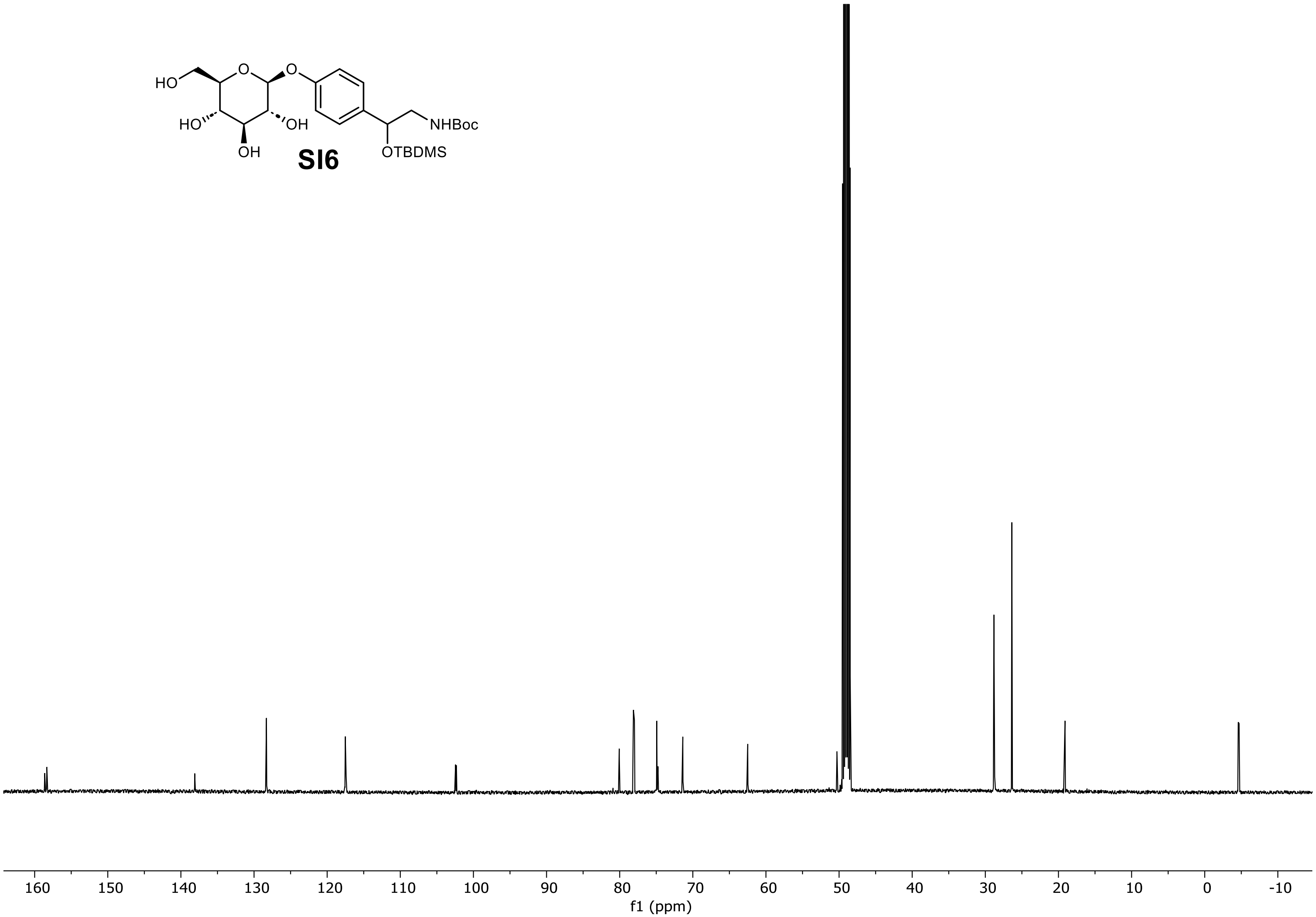

[1]H NMR of *tert*-butyl (2-((*tert*-butyldimethylsilyl)oxy)-2-(4-(((2*S*,3*R*,4*S*,5*S*,6*R*)-3,4,5-trihydroxy-6-(hydroxymethyl)tetrahydro-2H-pyran-2-yl)oxy)phenyl)ethyl)carbamate (**SI6**) in methanol-$d_4$ (126 MHz).

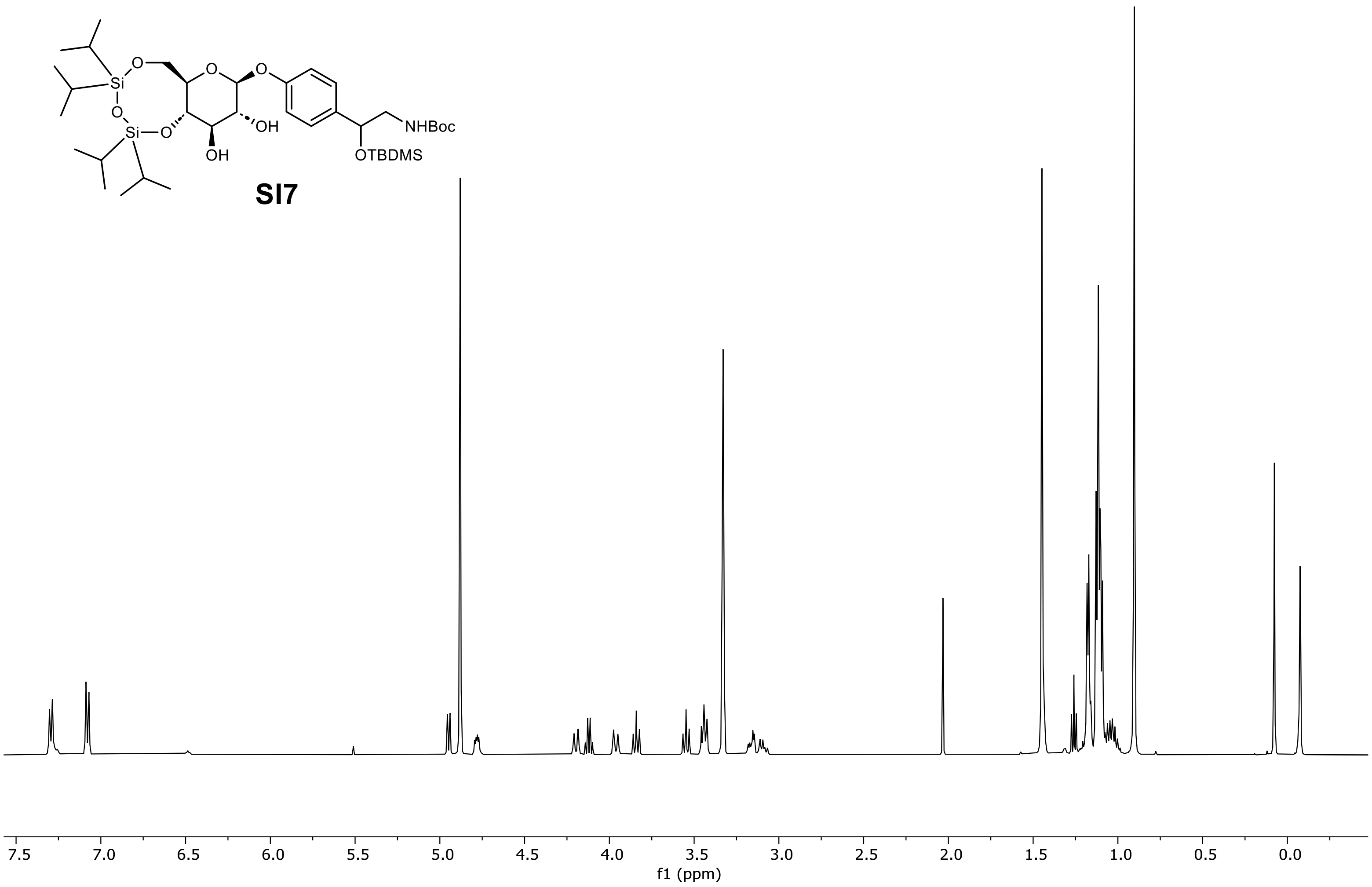

$^{1}H$ NMR of *tert*-butyl (2-((*tert*-butyldimethylsilyl)oxy)-2-(4-(((6a*R*,8*S*,9*R*,10*R*,10a*S*)-9,10-dihydroxy-2,2,4,4-tetraisopropylhexahydro-pyrano[3,2-f][1,3,5,2,4]trioxadisilocin-8-yl)oxy)phenyl)ethyl)carbamate (**SI7**) in methanol-$d_4$ (500 MHz).

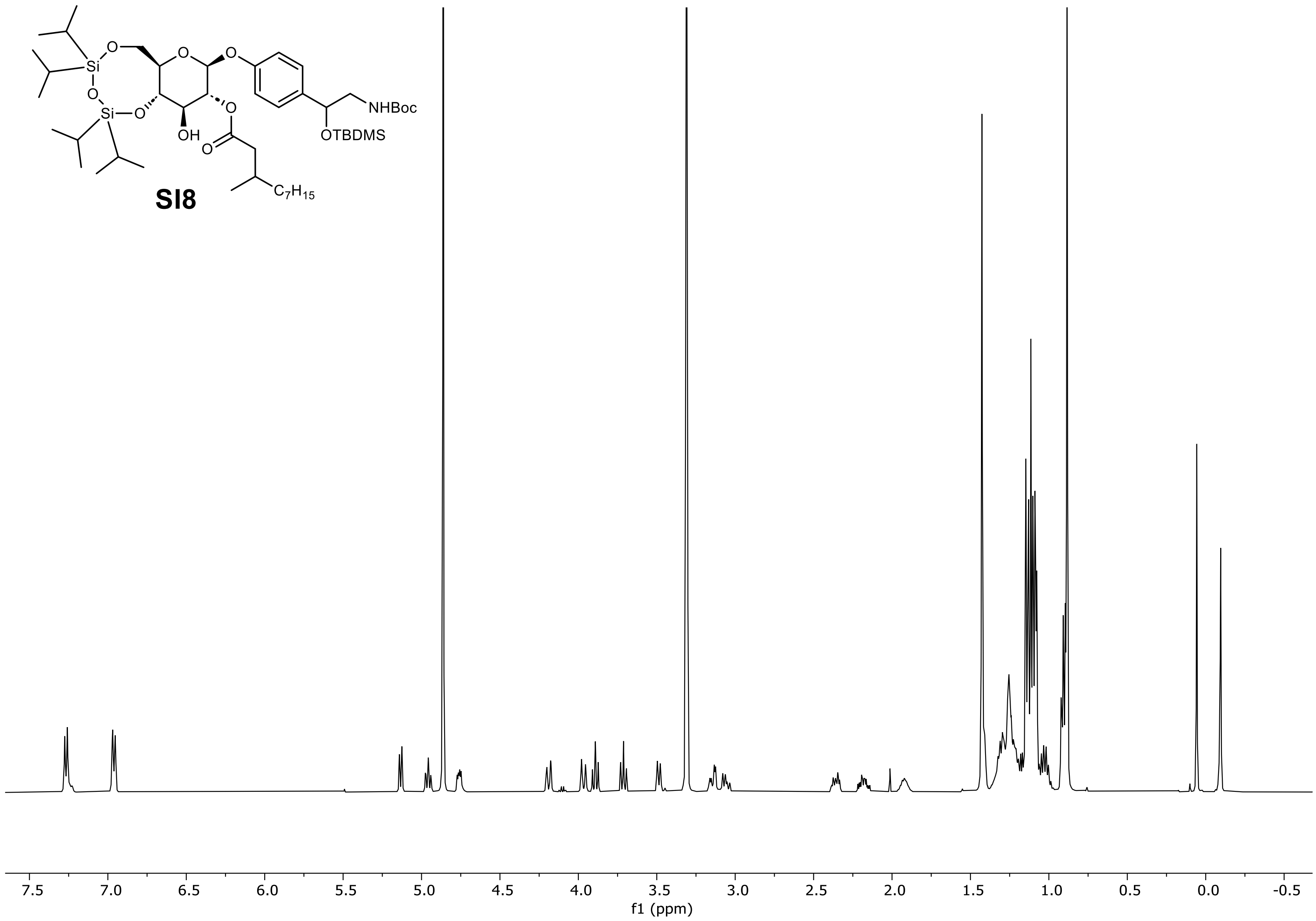

$^1$H NMR of (6a*R*,8*S*,9*R*,10*R*,10a*S*)-8-(4-(2,2,3,3,10,10-hexamethyl-8-oxo-4,9-dioxa-7-aza-3-silaundecan-5-yl)phenoxy)-10-hydroxy-2,2,4,4-tetraisopropylhexahydropyrano[3,2-f][1,3,5,2,4]trioxadisilocin-9-yl 3-methyldecanoate (**SI8**) in methanol-$d_4$ (500 MHz).

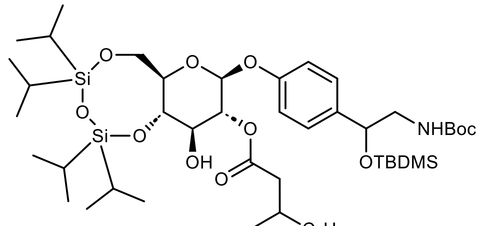

**SI8**

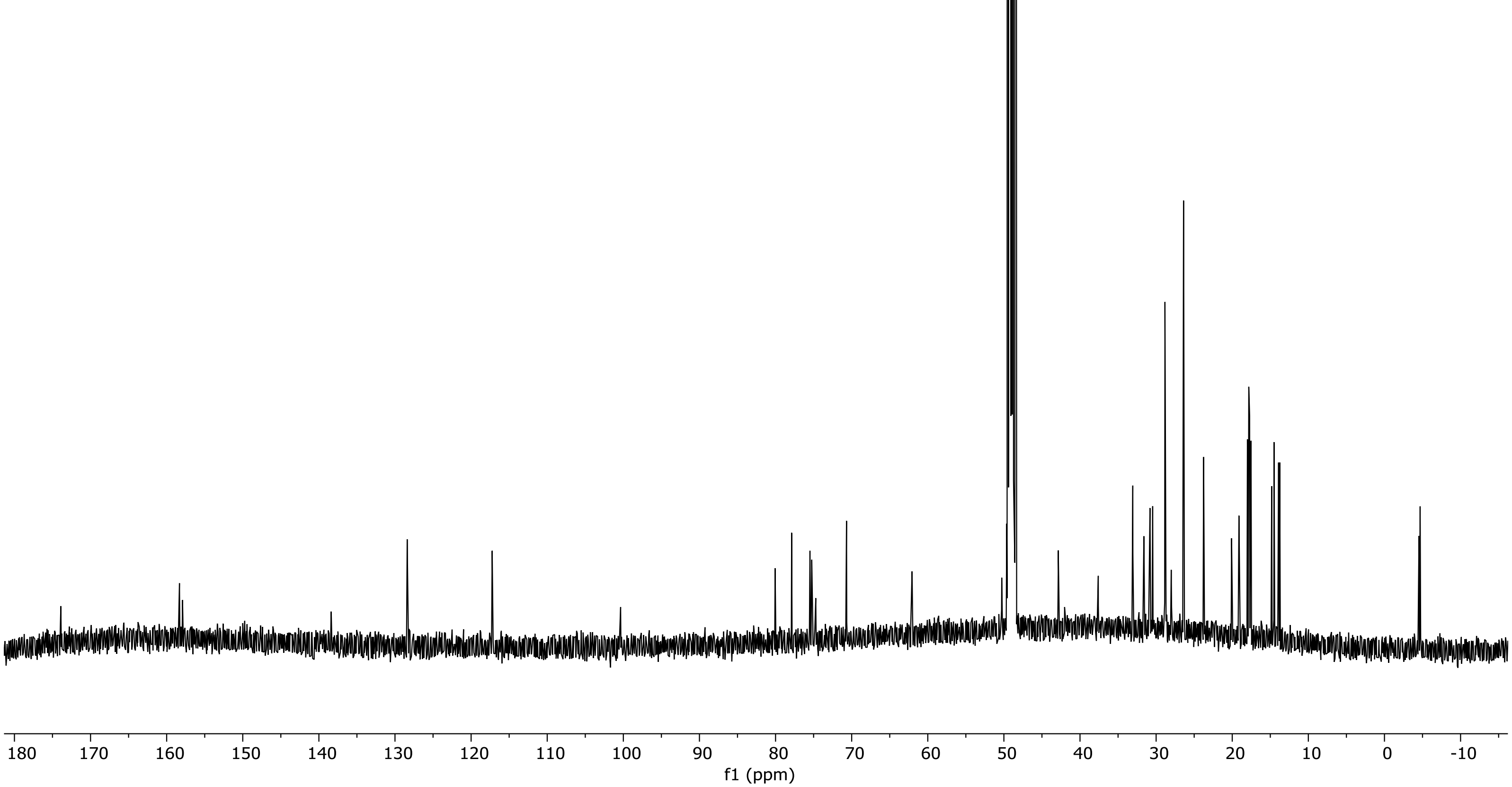

$^{13}$C NMR of (6a*R*,8*S*,9*R*,10*R*,10a*S*)-8-(4-(2,2,3,3,10,10-hexamethyl-8-oxo-4,9-dioxa-7-aza-3-silaundecan-5-yl)phenoxy)-10-hydroxy-2,2,4,4-tetraisopropylhexahydropyrano[3,2-f][1,3,5,2,4]trioxadisilocin-9-yl 3-methyldecanoate (**SI8**) in methanol-$d_4$ (126 MHz).

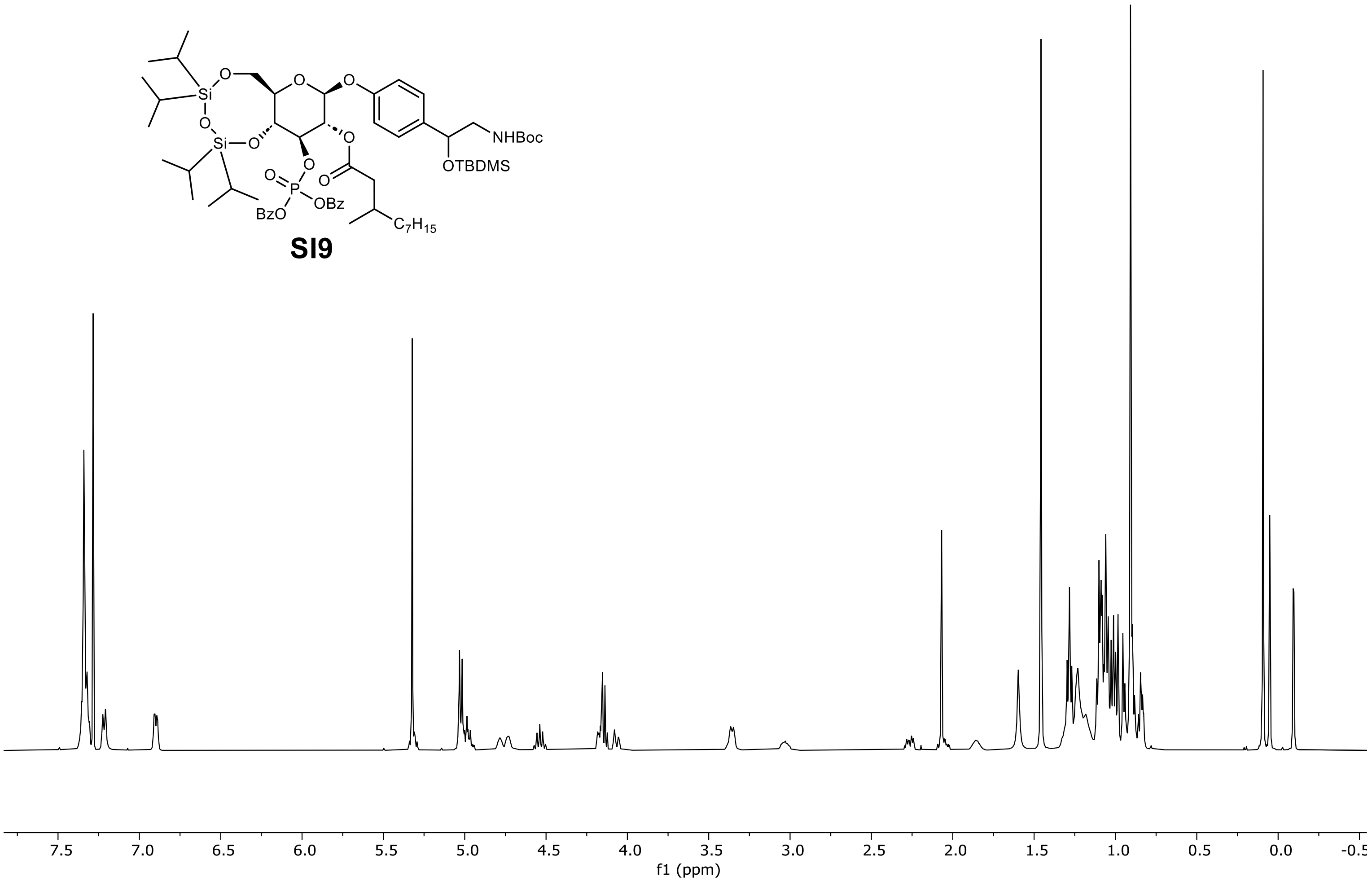

**SI9**

$^1$H NMR of (6a*R*,8*S*,9*R*,10*R*,10a*R*)-10-((bis(benzyloxy)phosphoryl)oxy)-8-(4-(2,2,3,3,10,10-hexamethyl-8-oxo-4,9-dioxa-7-aza-3-silaundecan-5-yl)phenoxy)-2,2,4,4-tetraisopropylhexahydropyrano[3,2-f][1,3,5,2,4]trioxadisilocin-9-yl 3-methyldecanoate (**SI9**) in chloroform-*d* (500 MHz).

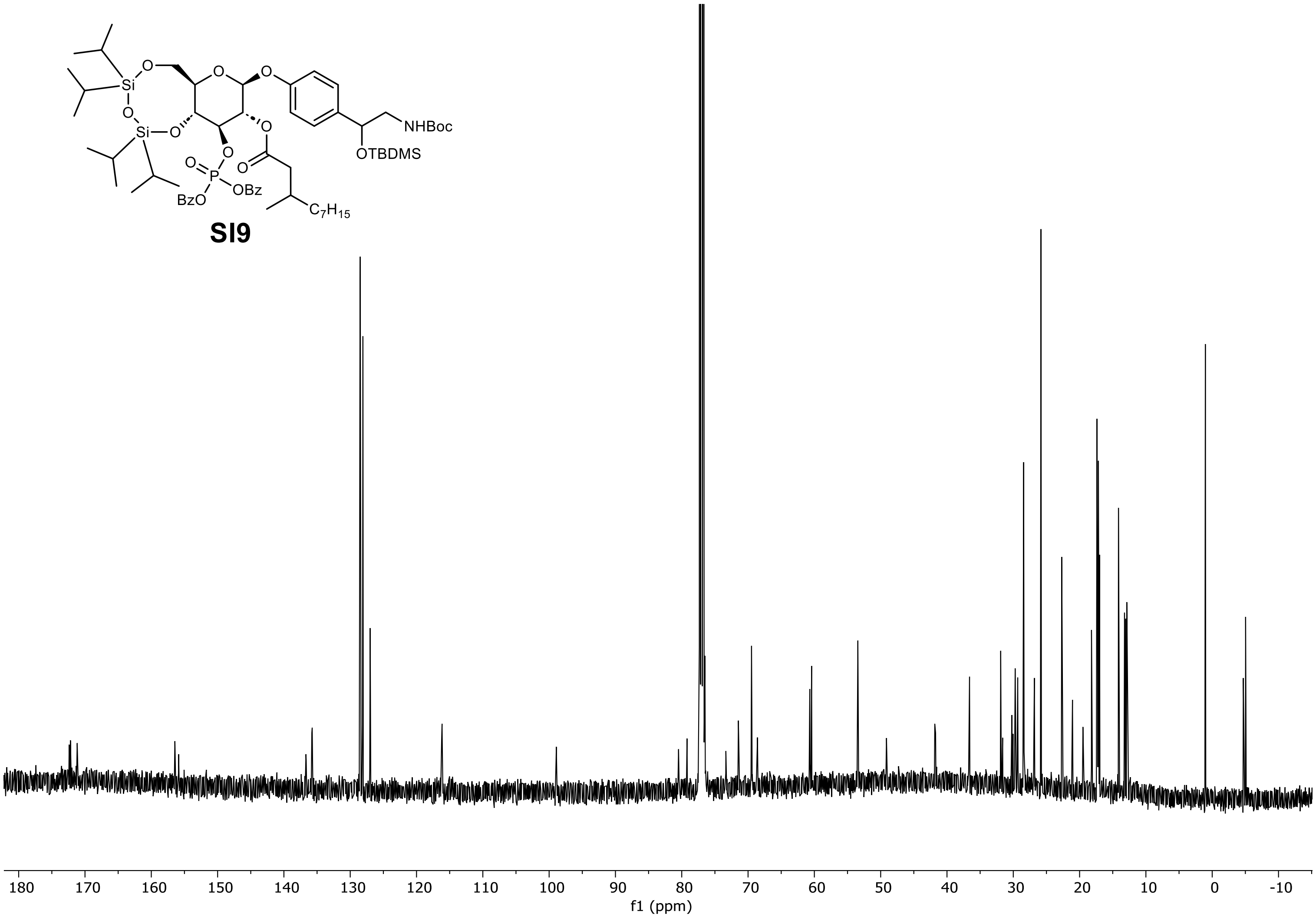

$^{13}C$ NMR of (6a*R*,8*S*,9*R*,10*R*,10a*R*)-10-((bis(benzyloxy)phosphoryl)oxy)-8-(4-(2,2,3,3,10,10-hexamethyl-8-oxo-4,9-dioxa-7-aza-3-silaundecan-5-yl)phenoxy)-2,2,4,4-tetraisopropylhexahydropyrano[3,2-f][1,3,5,2,4]trioxadisilocin-9-yl 3-methyldecanoate (**SI9**) in chloroform-*d* (126 MHz).

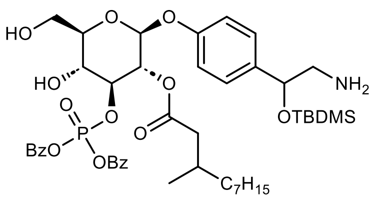

**SI10**

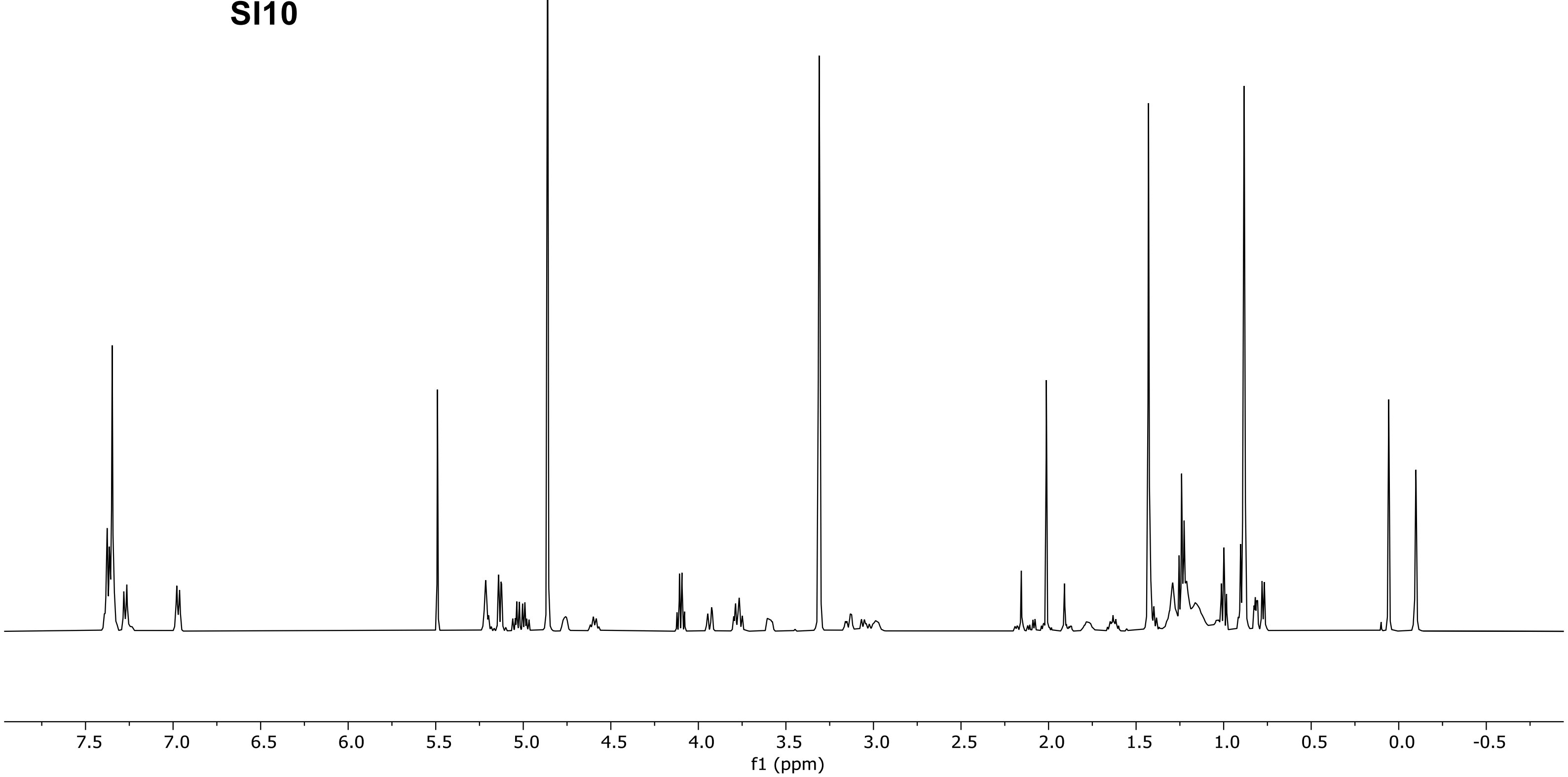

$^{1}$H NMR of (2*S*,3*R*,4*S*,5*R*,6*R*)-2-(4-(2-amino-1-((*tert*-butyldimethylsilyl)oxy)ethyl)phenoxy)-4-((bis(benzyloxy)phosphoryl)oxy)-5-hydroxy-6-(hydroxymethyl)tetrahydro-2H-pyran-3-yl 3-methyldecanoate (**SI10**) in methanol-$d_4$ (500 MHz).

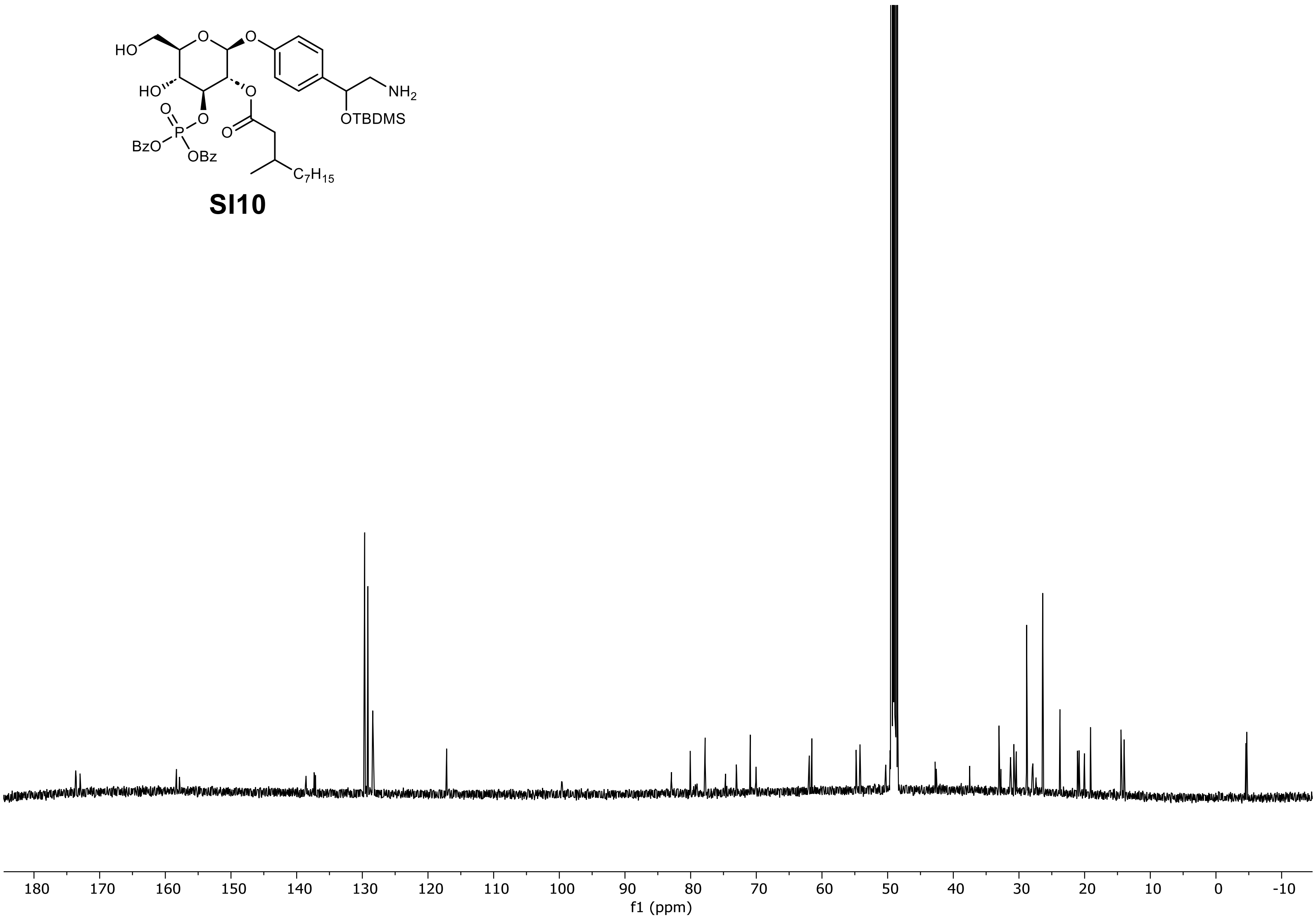

$^{13}$C NMR of (2*S*,3*R*,4*S*,5*R*,6*R*)-2-(4-(2-amino-1-((*tert*-butyldimethylsilyl)oxy)ethyl)phenoxy)-4-((bis(benzyloxy)phosphoryl)oxy)-5-hydroxy-6-(hydroxymethyl)tetrahydro-2H-pyran-3-yl 3-methyldecanoate (**SI10**) in methanol-*$d_4$* (126 MHz).

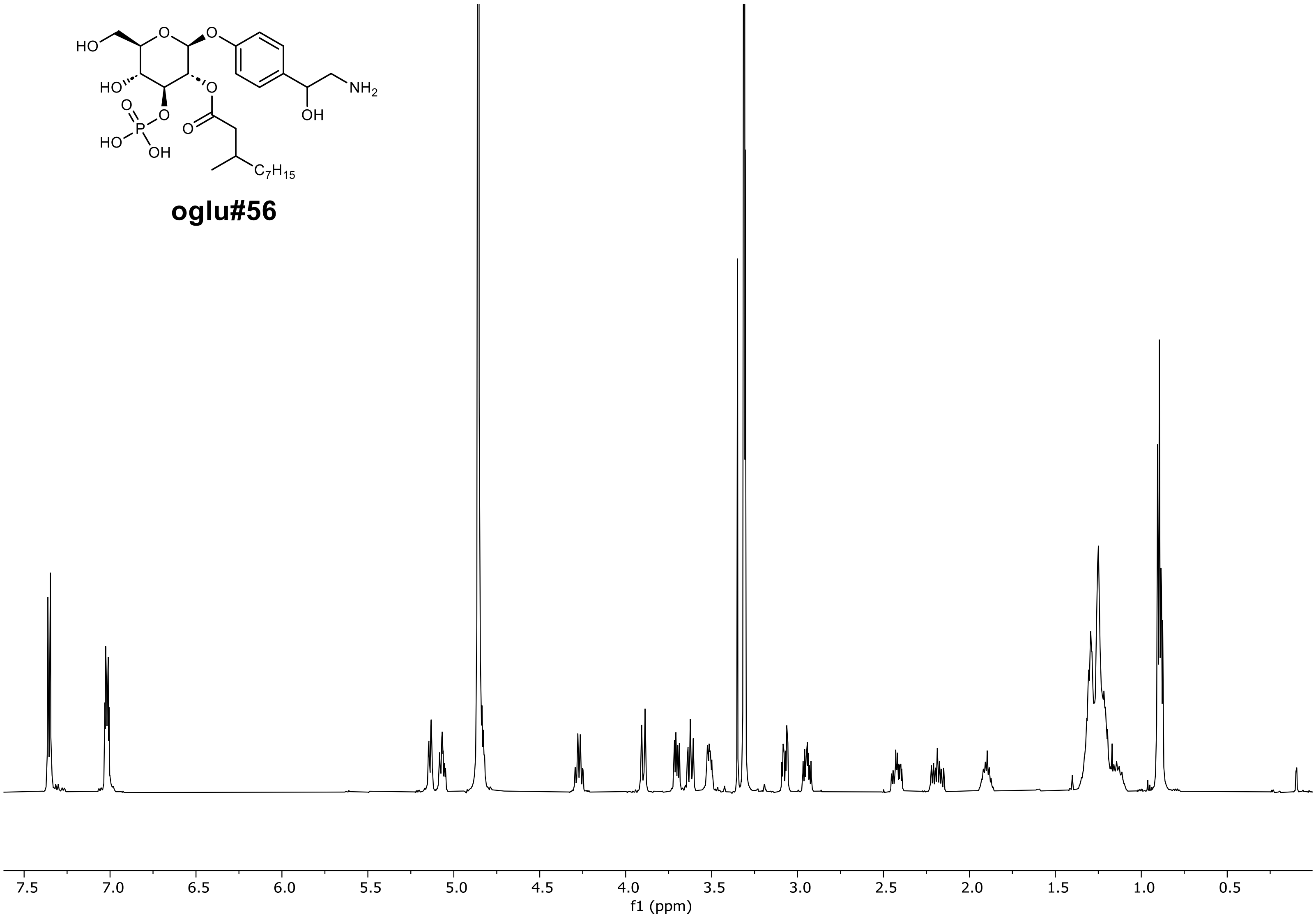

[1]H NMR of (2$S$,3$R$,4$S$,5$R$,6$R$)-2-(4-(2-amino-1-hydroxyethyl)phenoxy)-5-hydroxy-6-(hydroxymethyl)-4-(phosphonooxy)tetrahydro-2H-pyran-3-yl 3-methyldecanoate (**oglu#56**) in methanol-$d_4$ (600 MHz).

### 2.1 2D characterization of oglu#56

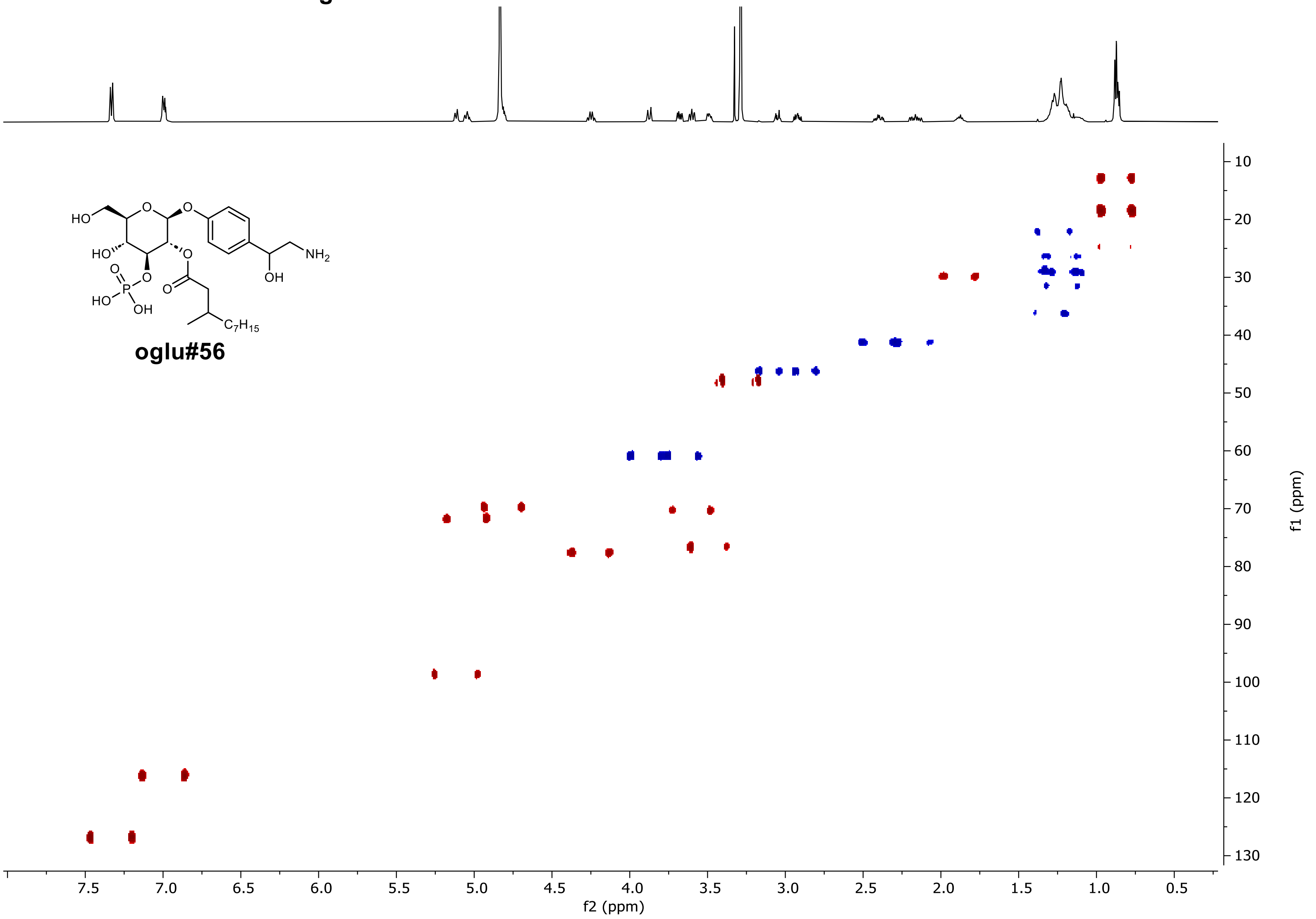

HSQC of (2*S*,3*R*,4*S*,5*R*,6*R*)-2-(4-(2-amino-1-hydroxyethyl)phenoxy)-5-hydroxy-6-(hydroxymethyl)-4-(phosphonooxy)tetrahydro-2H-pyran-3-yl 3-methyldecanoate (**oglu#56**) in methanol-$d_4$ (600 MHz).

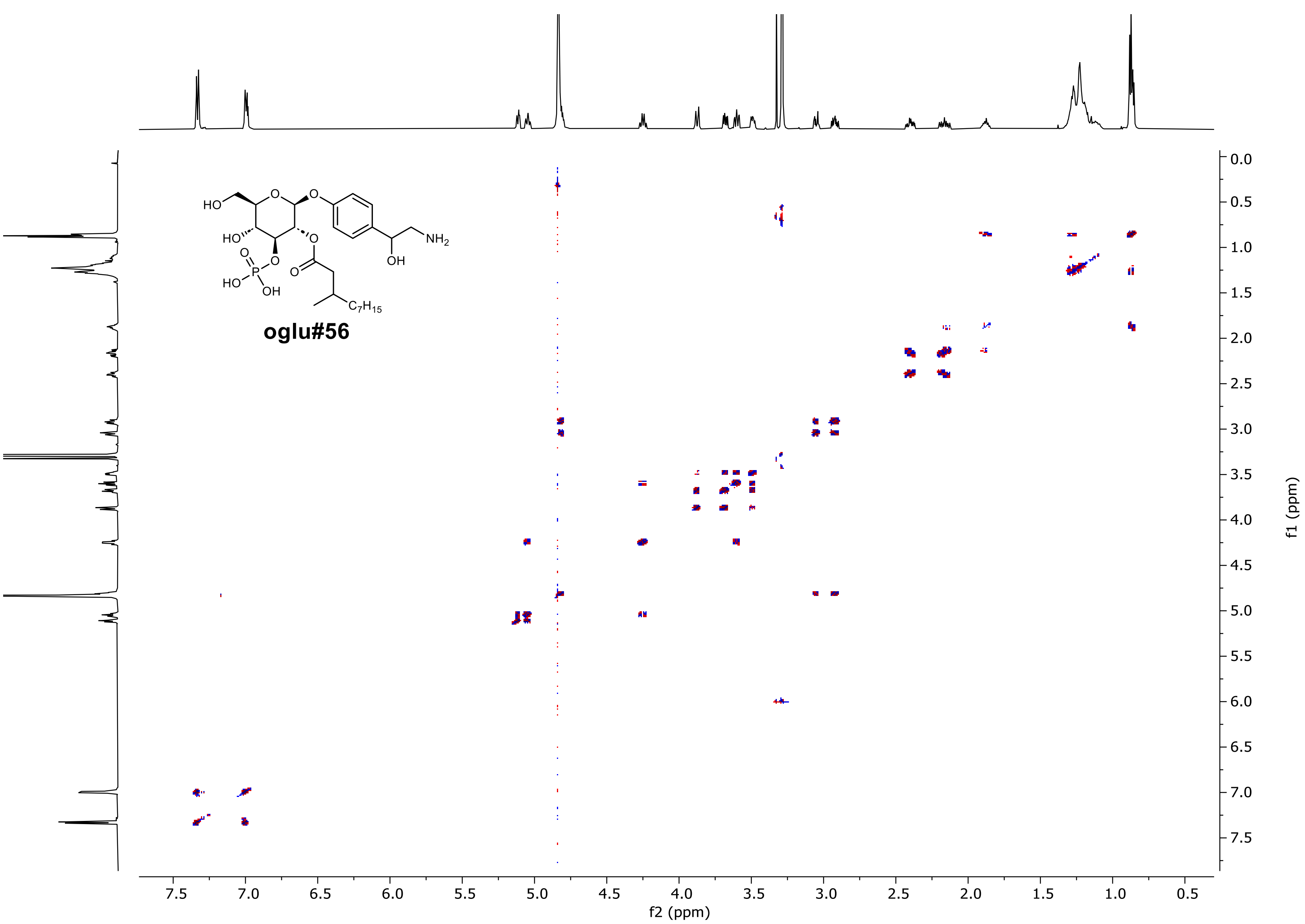
oglu#56
HO
HO
O
O
P
HO
OH
O
O
O
O
NH2
OH
C7H15
f2 (ppm)
f1 (ppm)
7.5
7.0
6.5
6.0
5.5
5.0
4.5
4.0
3.5
3.0
2.5
2.0
1.5
1.0
0.5
0.0

DQFCOSY of (2*S*,3*R*,4*S*,5*R*,6*R*)-2-(4-(2-amino-1-hydroxyethyl)phenoxy)-5-hydroxy-6-(hydroxymethyl)-4-(phosphonooxy)tetrahydro-2H-pyran-3-yl 3-methyldecanoate (**oglu#56**) in methanol-$d_4$ (600 MHz).

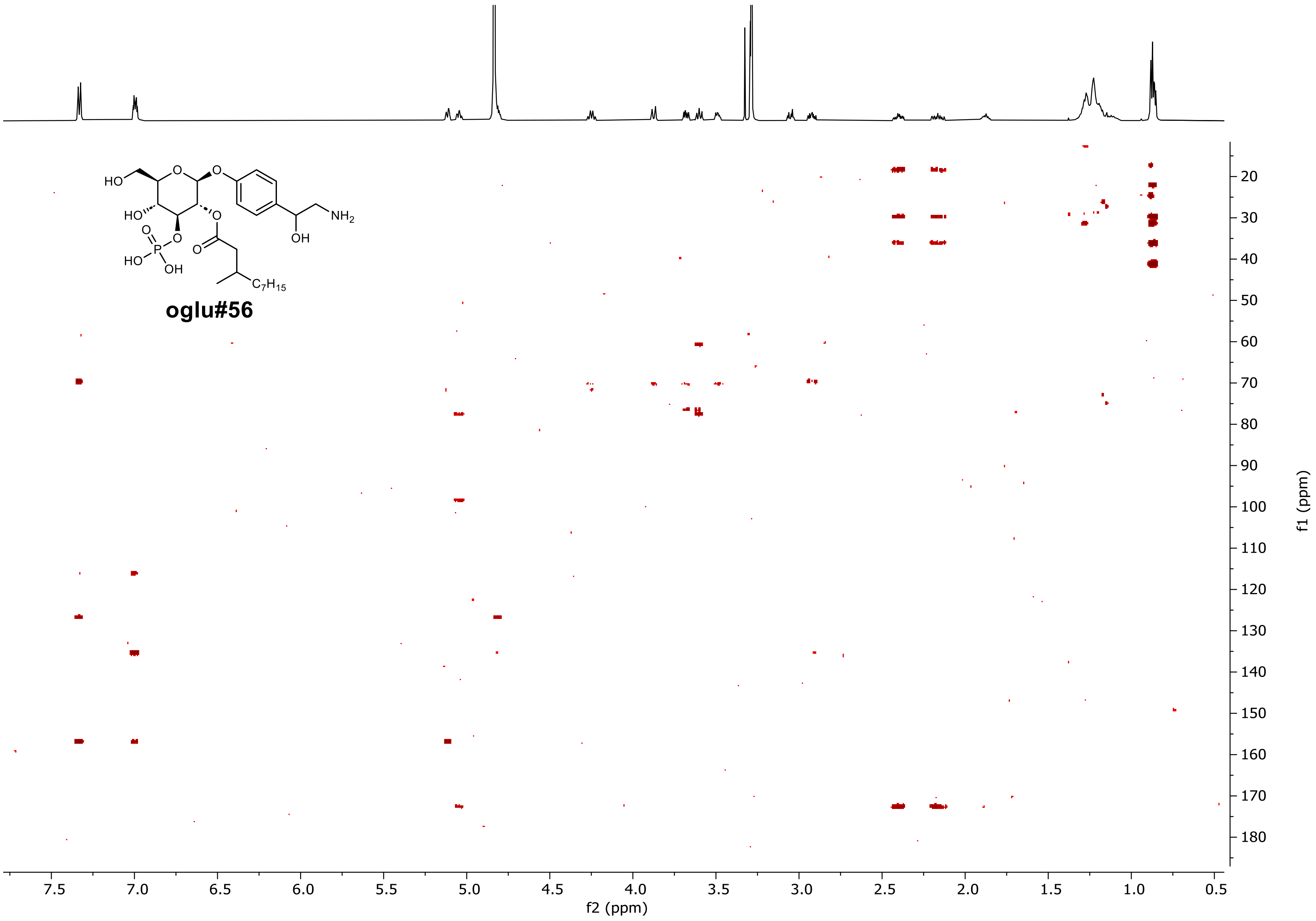

[1]H-[13]C HMBC of (2$S$,3$R$,4$S$,5$R$,6$R$)-2-(4-(2-amino-1-hydroxyethyl)phenoxy)-5-hydroxy-6-(hydroxymethyl)-4-(phosphono-oxy)tetrahydro-2H-pyran-3-yl 3-methyldecanoate (**oglu#56**) in methanol-$d_4$ (600 MHz).

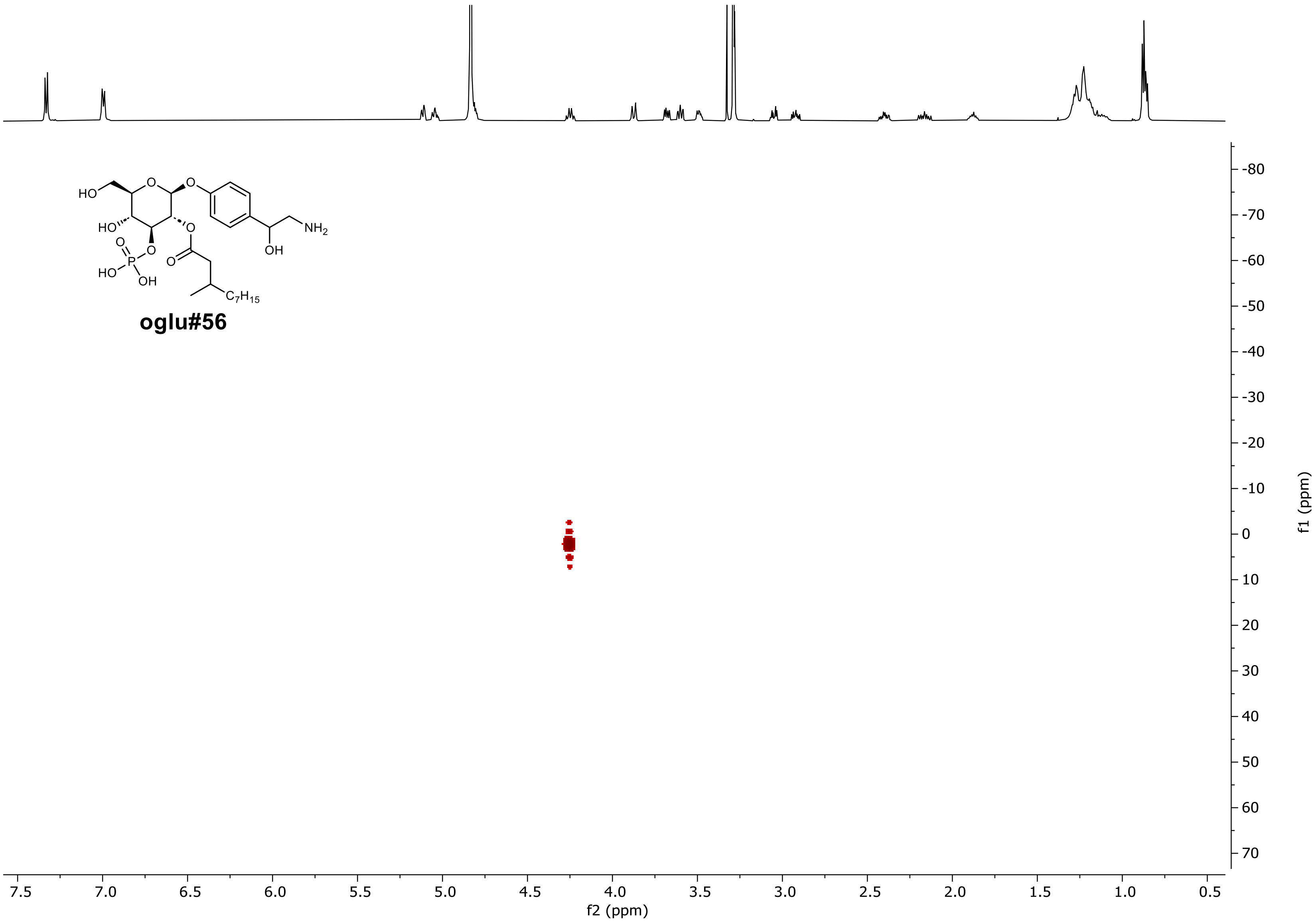

$^{1}H$-$^{31}P$ HMBC of (2*S*,3*R*,4*S*,5*R*,6*R*)-2-(4-(2-amino-1-hydroxyethyl)phenoxy)-5-hydroxy-6-(hydroxymethyl)-4-(phosphonooxy)tetrahydro-2H-pyran-3-yl 3-methyldecanoate (**oglu#56**) in methanol-$d_4$ (600 MHz).
